# Decoding the effect of temperatures on conformational stability and order of ligand unbound thermo sensing adenine riboswitch using molecular dynamics simulation

**DOI:** 10.1101/2024.12.14.628526

**Authors:** Soumi Das

## Abstract

The structure-function relationship of the riboswitch is governed mainly by two factors, ligand binding and temperature. Most of the experimental studies shed light on structural dynamics and gene regulation function of Adenine riboswitch from the aspect of ligand instead of temperature. Two unliganded Adenine riboswitch conformations (apoA and apoB) from the thermophile *Vibrio vulnificus* draw particular attention to the Biophysics research community due to their diverse and polymorphic structures. Ligand-free apoB Adenine riboswitch conformation is not able to interact with the ligand whereas ligand-free apoA Adenine riboswitch conformation adopts ligand-receptive form. The interconversion between apoA and apoB conformation is temperature-dependent and thermodynamically controlled. Therefore Adenine riboswitch is called a temperature sensing RNA. The molecular mechanism underlying the thermosensitivity of ligand free Adenine riboswitch is not well known. Hence it is essential to explore temperature-induced conformational dynamics of unliganded Adenine riboswitch. In this research work I make an attempt to examine conformational stability and order of apoA with respect to apoB Adenine riboswitch aptamer using conformational thermodynamics derived from all-atom molecular dynamics trajectories in the temperature range 283K-400K. The changes in conformational free energy and entropy of conformational degrees of freedom like pseudo-torsion angle ⍰ and θ are computed. RMSD, RMSF, R_G_, principal component analysis, hydrogen bonding interaction, and conformational thermodynamics data demonstrate that conformational stability and order of apoA with respect to apoB adenine riboswitch conformation is significant at 293K and 303K. The temperatures corresponding to the conformational order and stability of apoA adenine riboswitch with respect to apoB adenine riboswitch whole aptamer are shown in descending order 293K∼303K> 313K∼283K>373K>323K. The topological and conformational changes related to hydrogen bonding reorganization occur mostly at temperature 323K and 400K. Ligand unbound adenine riboswitch is sensitive to heat and may be inactivated at temperatures 323K and 400K.

**Highlights:**

- Molecular Dynamics Simulation of ligand unbound two different conformations of adenine riboswitch with the same sequence at different temperatures are performed.
- At low temperatures (293K and 303K) the conformational stability of apoA is more pronounced compared to apoB adenine riboswitch.
- Thermal stability of apoA compared to apoB adenine riboswitch is the least at 323K temperature.
- At higher temperature (373K) and lower temperature (283K) apoA adenine riboswitch exerts moderate conformational stability.
- The nucleotides from stem2 and junction (J_1/2_, J_2/3_) regions of adenine riboswitch contribute the most to the conformational stability and order.

## 1. Introduction

Riboswitch is a structured noncoding m-RNA found in all the three domains of the life including Bacteria, Archaea, and Eukaryota. Architecture of riboswitches are organized through a different kind of secondary structural elements such as pseudoknot, stem/helices, internal loops, kissing loops, junction region. Most studies on riboswitch till date shed light on ligand related gene expression mechanism (transcription, translation and alternative splicing) at the molecular level. Ligands accomplishes a wide variety of chemical entities such as nucleo-bases and their derivatives (e.g.; Adenine, Guanine), amino acids (e.g.; Lysine, Glycine), inorganic ions (e.g.; Mg^2+^, Mn^2+^, Ni^2+^, Co^2+^, F^-^), sugars, coenzymes and their derivatives (e.g.; S-adenosyl methionine, Flavin mono nucleotide). Riboswitch is divided into two compartments (i) evolutionary conserved sensory or aptamer domain and (ii) regulatory domain or expression platform. Riboswitch undergoes gene regulation expression by ligand dependent allosteric conformational switch mechanism. According to ligand dependent allosteric conformational switch mechanism, binding of ligand to aptamer domain causes transduction of genetic regulatory signal towards expression platform by conformational selection model. Ligand-aptamer interactions at the molecular level reveal different kinds of interactions: base: base, base: amino acid, base: sugar, base: phosphate etc. In addition to ligands or metabolites, riboswitches utilize different factors as for example temperature, uncharged t-RNA, metal ions to achieve regulation of gene expression [**1-5**].

Adenine riboswitch from Vibrio vulnificus is reported as a temperature-sensing riboswitch. A bi-stable structural element of Adenine riboswitch having two different conformations apoA and apoB with mutually exclusive secondary structures (**Figure 1.(a)**), is responsible for temperature dependent translational initiation. Experimental study explores that the conformational equilibrium shifts toward the binding incompetent conformation apoB at low temperatures (e.g.; 10° C) whereas binding competent apoA conformation populates at physiological temperatures (e.g.; 37° C). Efficient conformational switching, governed over a large temperature range, plays an important role in Vibrio biology because pathogen replicates around 10° C in its marine habitat and at 37° C in human host [**6**].

**Figure 1.**
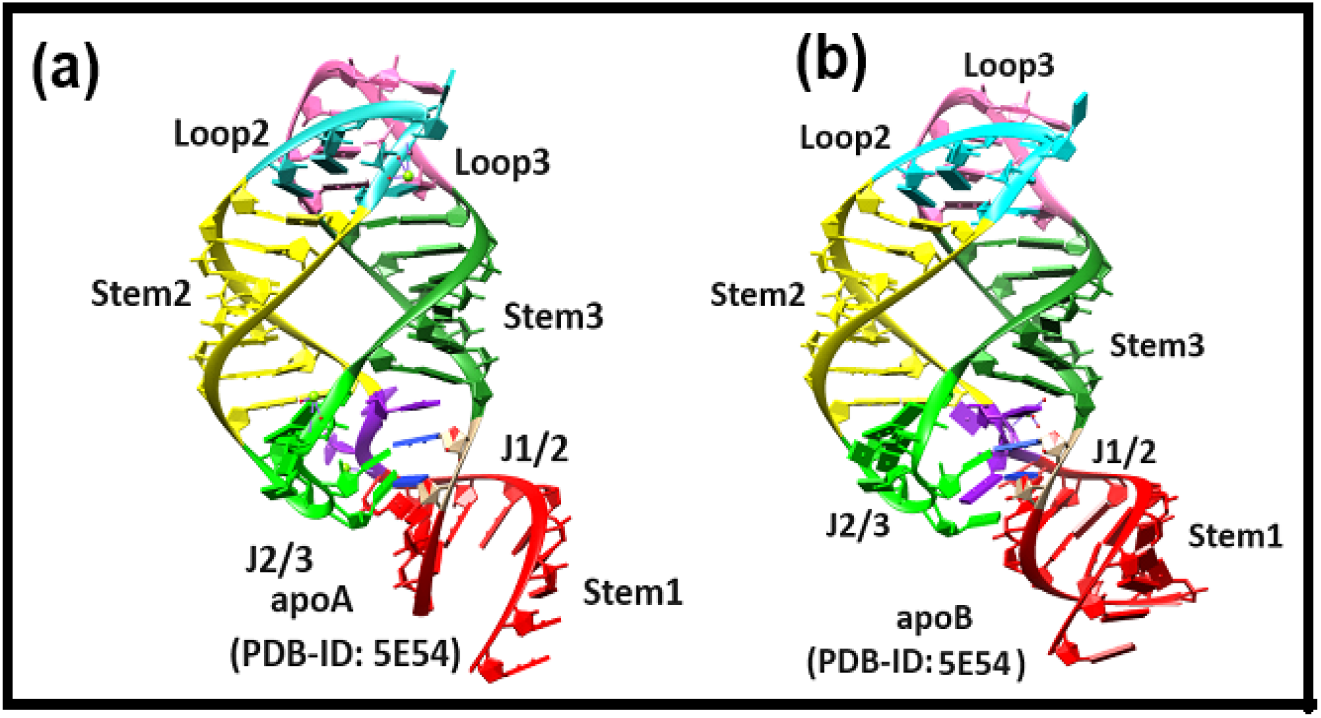
(a) Crystal structure of two different mutually exclusive conformations of ligand unbound adenine riboswitch aptamer: apoA (left) and apoB (right).

Adenine riboswitch aptamer consists of three helices [stem1, stem2, stem3], kissing loops [L2 and L3] and junction loops [J_12_, J_23_, J_31_] **(Figure 1. (a))**. Several research groups [**7-20]** shed light on structure-function relationship, folding-unfolding dynamics, and gene regulation of the adenine riboswitch from the aspects ligand instead of temperature using NMR, ITC, stopped-flow, in vivo assay and molecular dynamics simulation. However, there has yet not been any attempt to compare the temperature mediated conformational dynamics of ligand unbound apo A with respect to the apo B conformations of adenine riboswitch aptamer in the temperature range 283K-400K using molecular dynamics simulation.

In this work, thermodynamics of region-wise conformational change of apo A with respect to the apo B is extracted from the histograms of microscopic conformational variables. The histogram-based method (HBM) for conformational thermodynamics estimation which was initially developed for the proteins [**21-26]** further extended to the protein-DNA complex systems [**27-28]** as well. Here I extend the conformational thermodynamics technique to the RNA system using the microscopic conformational variables of RNA named pseudo torsion angle □ and θ at different temperatures 283K, 293K, 303K, 313K, 323K, 373K and 400K captured from molecular dynamics trajectory to examine conformational stability of apoA with respect to apoB conformations of the adenine sensing riboswitch in aqueous solution to decipher the influence of temperatures on the structure, function, and dynamics of ligand unbound adenine riboswitch. Our goal is to explore the intrinsic relationship between the structural cores, i.e. stem region, kissing loops and junctions and how they mutually affect each other at different temperatures.

Findings from my study reveal that stem2 and junction regions (J_1/2_ and J_2/3_) exhibit the highest conformational stability and order of the ligand unbound adenine riboswitch whereas stem1 is the least stable and ordered. At low temperatures (293K and 303K), apoA exerts significant conformational stability and order compared to apoB adenine riboswitch. At higher temperature (373K) and lower temperature (283K) apoA adenine riboswitch exhibits moderate conformational stability. Thermal stability of apoA compared to apoB adenine riboswitch is the least at 323K temperature.

Ligand unbound adenine riboswitch is sensitive to heat and may be inactivated at temperatures 323K and 400K. The topological and conformational changes related to hydrogen bonding reorganization occur at temperature 323K and 400K. The deterioration of the hydrogen bonding network appears as the main reason for the functional inactivation of the apo adenine riboswitch upon heating. Understanding the temperature driven changes in ligand unbound adenine riboswitch dynamics enhances our comprehension of riboswitch thermodynamics which is essential for illuminating the ‘riboswitch-thermostat’ concept at dynamic molecular level.

## 2. Methods

### 2.1 System Preparations

Two distinct conformations of ligand unbound adenine riboswitch aptamer domain (Vibrio vulnificus strain 93U204 chromosome II) extracted from chain A and chain B of PDB ID-5E54 are named apoA (ligand binding competent) and apoB (ligand binding incompetent). The lists of the ligand unbound adenine riboswitch systems studied across the varied thermal conditions are shown in the Table 1.

**Table 1:** List of systems studied.

| System | Temperature | Duration of Molecular dynamics simulation | Force field | RMSD (nm) |
| --- | --- | --- | --- | --- |
| apoA form of Adenine riboswitch aptamer (PDB-ID: 5E54)<br>ligand binding competent | 283K | 1 μs | $\chi_{OL3}$ RNA Force field | 0.35 (0.07) |
| | 293K | 1 μs | | 0.24 (0.04) |
| | 303K | 1 μs | | 0.25 (0.05) |
| | 313K | 1 μs | | 0.26 (0.07) |
| | 323K | 1 μs | | 0.36 (0.05) |
| | 373K | 1 μs | | 0.40 (0.05) |
| | 400K | 1 μs | | 0.48 (0.08) |
| apoB form of Adenine riboswitch aptamer (PDB-ID: 5E54)<br>ligand binding incompetent | 283K | 1 μs | $\chi_{OL3}$ RNA Force field | 0.25(0.05) |
| | 293K | 1 μs | | 0.27(0.04) |
| | 303K | 1 μs | | 0.32(0.06) |
| | 313K | 1 μs | | 0.27(0.05) |
| | 323K | 1 μs | | 0.29(0.05) |
| | 373K | 1 μs | | 0.37(0.06) |
| | 400K | 1 μs | | 0.54(0.13) |

### 2.2 Simulation protocol

Molecular dynamics simulations are carried out using the GROMACS version 2022.2 [**29]** to examine the possible differences in the dynamical behaviour between apoA and apoB adenine riboswitch using force field parameters Amber OL3, also known as χ_OL3_ [**30**]. Mg^2+^ in the crystal structures are kept in both the apoA and apoB systems because Mg^2+^ ions are essential to confer stability of the riboswitch. Removal of water of crystallization from two RNA systems named apoA and apoB are necessary step before performing molecular dynamics simulation. The RNA systems are solvated in a cubic box with 15 Å thick layers of TIP3P water molecules for explicit solvent calculation [**31]** and neutralized with the addition of the required number of [Na^+^] and [Cl^-^] ions. Particle-Mesh Ewald Summation (PME) is used for long-ranged electrostatic interactions with 1 Å grid spacing and 10^-6^ convergence criterion. The Lennard-Jones and the short-range electrostatic interactions are truncated at 10 Å. The LINCS constraints are applied to all the bond involving hydrogen atoms. The periodic boundary conditions are imposed in all directions. Further four steps are involved in Molecular dynamics simulations: (i) energy minimization, (ii) thermalization, (iii) equilibration, and (iv) production simulation. The RNA systems are energy minimized with the help of the Steepest Descent algorithm and the Conjugate Gradient algorithm for the removal of steric clashes. Energy minimization steps using Steepest Descent algorithm are carried out with heavy atom restraints by applying a harmonic potential with a force constant of 1000 kJ mol^-1^. Application of the restraints are not allowed in case of Conjugate Gradient algorithm. After completion of energy minimization, the RNA systems are slowly heated stepwise to each of the seven desired temperatures (283K, 293K, 303K, 313K, 323K, 373K and 400K) under NVT condition. Finally the RNA systems are equilibrated for 100 ps under the isoberic-isothermal (NPT) ensemble using a time step of 2 fs. The pressure and temperature are controlled by Parrinello-Rahman barostat [**32]** and Berendsen thermostat [**33]** respectively. The final production runs are carried out for 1 μs of each system at the seven temperatures. Integration time step of 1 fs is set for the production run. The trajectories are visualized employing VMD program and images are captured with the help of CHIMERA [**34]** and PYMOL [**35**].

### 2.3 Structure analysis

The root mean square deviation (RMSD), root mean square fluctuation (RMSF), and radius of gyration (R_g_) values are computed using GROMACS embedded tools. The equilibrations of trajectories are judged from the RMSD. The backbone phosphorous (P) atom based root mean square fluctuation (RMSF) per nucleotide is calculated to understand which regions of apoA and apoB adenine riboswitch undergo the most conformational fluctuation or structural rearrangement in response to different thermal environments. The radius of gyration (R_g_) is analyzed to distinguish the compactness of apoA and apoB adenine riboswitch under different temperatures. Base pairing information of RNA systems are detected by BPFIND [**36**]. Different pseudo-torsion angles are calculated using NUPARM software [**37**].

### 2.4 Identification of interactions

The hydrogen bond interactions are characterized by the distance and angle criteria using Gromacs Hydrogen Bond Analysis module. The hydrogen bonds are taken under consideration when the distance between the donor (D) and acceptor (A) is less or equal to 3.5 Å and the angle (D-H-A) cut-off is 160°.

### 2.5 Conformational thermodynamics

The detailed protocol of the histogram-based method (HBM) for computation of the conformational thermodynamics derived from equilibrated molecular dynamics trajectory is stated in preceding publications [**21-28**]. The histograms represent the probability of finding the system in a given conformation. The histograms and the conformational thermodynamics are interconnected with the help of Boltzmann factors and Gibbs formula corresponding to effective free energy and entropy respectively. The histograms of microscopic conformational variables are generated using the equilibrated trajectory, namely, 600 to 1000 ns. The normalized probability distribution of any microscopic conformational variable θ in the apoA state and the apoB state is given by *H*^*apoA*^ (*θ*) and *H*^*apoB*^ (*θ*) respectively. The change in free energy of any microscopic conformational variable θ of the apoA with respect to that of the apoB is defined by the following formula:

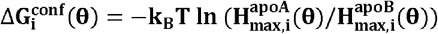

Where “max” represents the peak value of the histogram and i represents each of the RNA residues.

The change in conformational entropy of a given microscopic conformational variable θ the apoA with respect to that of the apoB is evaluated as,

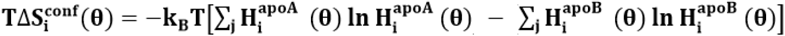

Where the sum is taken over all histogram bins j and i represent each of the RNA residues.

k_B_ And T denote Boltzmann constant and temperature respectively.

The computation of conformational stability and order are carried out in terms of conformational thermodynamics using the code located in https://github.com/snbsoftmatter/confthermo.

### 2.6 Essential Dynamics

Essential dynamics are calculated at different temperatures through principal component analysis [**38]** with the help of Gromacs analysis tool gmx covar module in which a covariance matrix of 3N*3N dimension is generated with N being P atomic fluctuation. Eigenvectors and eigenvalues are obtained from covariance matrix. Eigenvectors represent the direction of motion along the principal components whereas the eigenvalue of an eigenvector defines how much data is dispersed along the eigenvector. The dynamic cross correlated maps are generated with the help of Bio3D library of R program [**39]** to explore correlated and anti-correlated motion. The extent of correlated and anti-correlated motions which are defined correlation coefficient **C**_**(i**, **j)**_ between i^th^ and j^th^ atoms, are expressed by the following formulas:

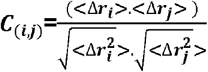

where Δ*r*_*i*_ and Δ*r*_*j*_ are the displacement of i^th^ and j^th^ atom from mean position and <> symbol represents the time average.

2D projection plots from simulated trajectory are generated using gmx anaeig module.

### 2.7 Gibbs Free Energy Landscape

Free Energy Landscape (FEL) is generated using gmx sham module to capture the lowest free energy state. The FEL landscape is projected on PC1 and PC2.

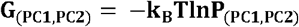

Where k_B_, T and P_(PC1,PC2)_ denote Boltzmann constant, temperature and normalized joint probability distribution respectively.

## 3 Results and Discussion

MD simulation average snapshots for apoA and apoB adenine riboswitch at different temperatures are represented at Figure S1.

### 3.1 Global and local stability of the aptamer domain

The structural stability of riboswitches in aqueous solution, in both apoA and apoB configurations, are evaluated by comparing the average RMSD (Table 2. and Supplementary Figure S2). The radius of gyration (R_g_) (Table 3.) and root-mean-square fluctuation (RMSF) (Figure S3) values are calculated over the trajectory production stage, taking crystallographic structure as a reference structure. The relative variability of the apoA and apoB systems are illustrated by the root-mean-square deviations (RMSD) of phosphodiester backbone from the corresponding initial structures as a function of time at seven different temperatures. The computed averaged RMSD are ranged between 2.5–3.5 Å approximately for the temperature 283K, 293K, 303K, 313K, and 323K. On increasing the temperature to 400K, average RMSD of whole aptamer of apo A and apo B increase to 4.8 Å and 5.4 Å respectively. The inspection over some particular regions of the RNA aptamer evidence slight differences in their dynamic behaviour. For example, the stem1 (P1) region presented higher RMSD values than that of all other substructures in both species. Stem1 of apoA exhibits significant distortion at 400K and 323K as observed from Table 1. It is worth mentioning that the nucleotides composing the Stem1 or P1 helix are located in the terminal regions of the adenine riboswitch aptamer, showing broader movements during MD simulations. Unlike the stem2 and stem3 segment, J_1/2_ (Hinge) and J_2/3_ (Latch) of apoB display higher RMSD at 400K, 303K, and 293K. No noticeable differences have been observed between the RMSD values obtained for stem2, stem3, loop2, and loop3 for both apoA and apoB adenine riboswitch at different temperatures. The higher backbone RMSD of both apoA and apoB whole aptamer at elevated temperatures indicate less stability of ligand unbound Adenine riboswitch at higher temperatures. Distinct dynamics of apoA and apoB are clearly reflected at 283K and 400K from Figure S2.

**Table 2:**
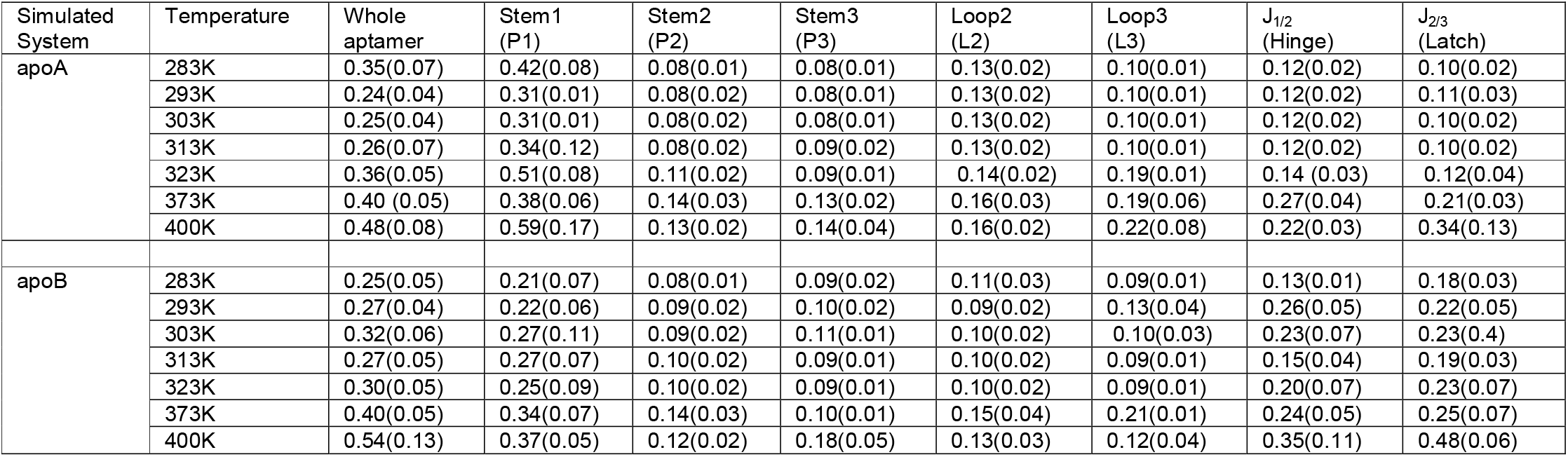
Root mean square deviations (RMSD) of apoA and apoB adenine riboswitch as a whole and for substructures (nm)

**Table 3:** Radius of gyration (R_g_) of apoA and apoB adenine riboswitch aptamer (nm)

| Simulated System | Temperature | Whole aptamer |
| --- | --- | --- |
| apoA | 283K | 1.84(0.05) |
|  | 293K | 1.88(0.03) |
|  | 303K | 1.88(0.03) |
|  | 313K | 1.87(0.03) |
|  | 323K | 1.84(0.04) |
|  | 373K | 1.78(0.04) |
|  | 400K | 1.73(0.04) |
| apoB | 283K | 1.96(0.03) |
|  | 293K | 1.96(0.03) |
|  | 303K | 1.95(0.03) |
|  | 313K | 1.93(0.04) |
|  | 323K | 1.94(0.03) |
|  | 373K | 1.85(0.04) |
|  | 400K | 1.81(0.04) |

R_g_ gives the estimation of a system’s compactness as the higher R_g_ value indicates lower compactness. The average R_g_ value (Table 3.) of apoA is lower across the temperatures indicating tighter packing of apoA adenine riboswitch compared to apoB adenine riboswitch. R_g_ values for both apoA and apoB adenine riboswitch decrease with increase the values of the temperatures. The lowest average RMSF values (Figure S3) are observed with apoA followed by apoB suggesting these two riboswitches to be more rigid at 283K and 313K. The conformation of apoA and apoB Adenine riboswitch are most affected by changes in temperature 293K and 400K reflected from RMSF. All structures showed enhanced flexibility at stem1 region located in the terminal regions of the aptamer. Another common region with enhanced flexibility was at J_2/3_ position. apoB shows enhanced flexibility compared to apoA at 293K and 303K. The most interesting observation is that the whole aptamer of apoA and apoB at 323K, 373K and 400K exhibit more significant fluctuation. RMSF results thus clearly confirm that apoA is thermodynamically more stable at 283K, 293K, 303K, 313K and 373K. In contrast both apoA and apoB is unstable at 400K.

### 3.2 Dynamics of canonical and non-canonical hydrogen bonds at different temperatures

The occupancies of the hydrogen bonds that are involved in the formation of the base pair (Table 4.) are computed in the stem1, stem2 and stem3, loops L2, loop L3 and J_2/3_ across the varied thermal condition. For apoA and apoB, almost all base pairs of stem1 at 283K, 323K and 400K result in an overall destabilized hydrogen bond network in our MD simulations, manifested in lower occupancies for the hydrogen bonds (< 50%) while at 293K, 303Kand 313K hydrogen bond networks get moderately stable. In contrast hydrogen bond occupancies of Stem1 of apoA and apoB at 373K exhibits the highest stbility. Hydrogen bond occupancies of stem2 of apoA and apoB decrease significantly at 323K (< 70%) and 400K (< 40%) whereas all base pairs of stem2 are significantly stable (hydrogen bond occupancy > 90%) at 283K, 293K, 303K, 313K, 373K. An overall destabilized hydrogen bond network of stem3 is observed at almost all temperatures (283K, 303K, 313K, 373K and 400K.) reflected from hydrogen bond occupancy. Hydrogen bonding at base pairs of stem3 at 293K (hydrogen bond occupancy > 97%) is highly stable except 58A 68U (hydrogen bond occupancy > 61%). Transient weak hydrogen bonds destabilize Loop2 and loop3 at 323K and 400K. In contrast loop2 and loop3 get stabilized by almost all hydrogen bonded both canonical and non-canonical base pairs (hydrogen bond occupancy > 90%) at temperatures 283K, 293K, 303K, 313K, 373K. All hydrogen bonds of apoA and apoB at J_2/3_ are disrupted completely at 400K. In case of 323K temperature, weak hydrogen bond occupancy (< 35%) is observed at apoB while apoA conformation exhibit complete disrupted hydrogen bond at J_2/3_. In contrast, the strength of the hydrogen-bond network of J_2/3_ of apoA and apoB at temperatures 283K, 293K, 303K, 313K, 373K are significant.

**Table 4:**
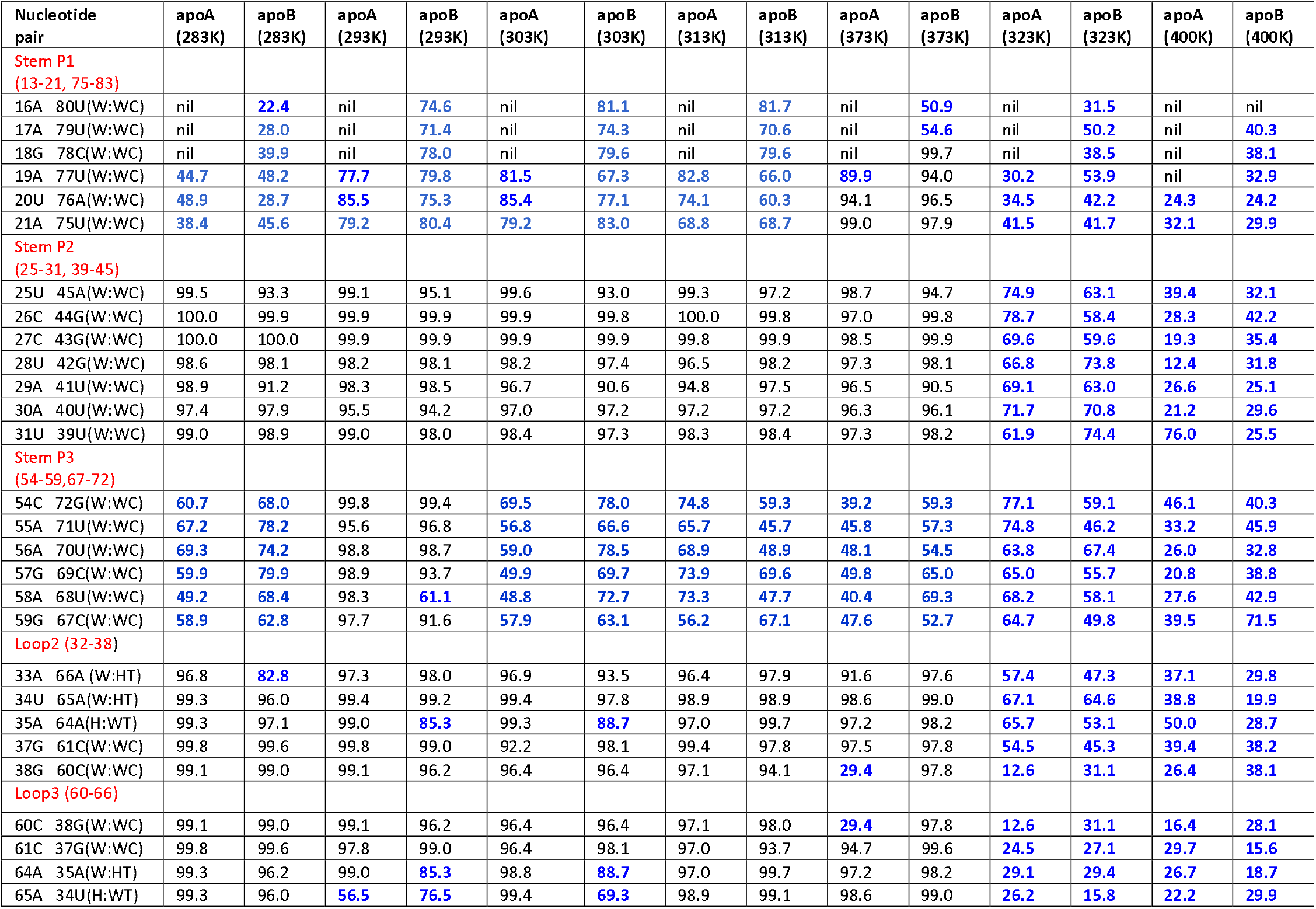
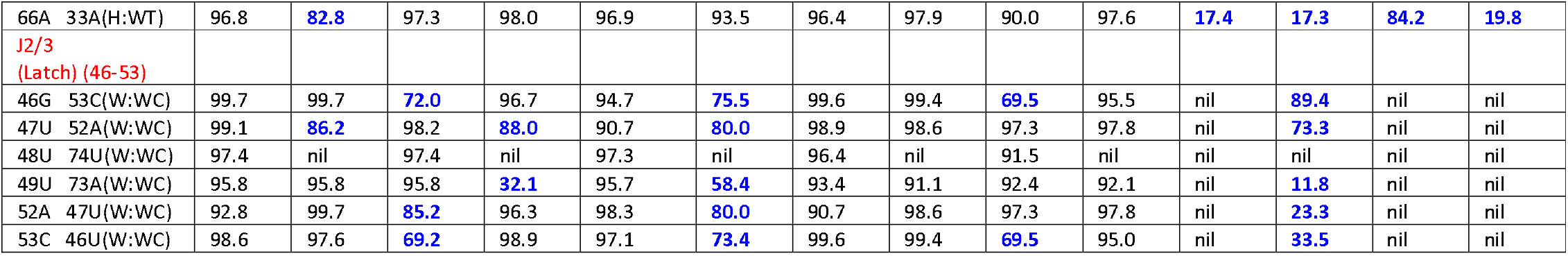
Hydrogen bond occupancy in different region of apoA and apoB adenine riboswitch under different temperatures.

### 3.3 Comparative analysis of fluctuations in pseudo torsion angle

The RNA backbone possess seven conventional torsional degrees of freedom (α, β, γ, δ, ε, ζ) which delineate rotation around the bond. Besides there are glycosidic torsion angle χ which defines the rotation between the ribose sugar and base. Five endocyclic torsion angles (ν_o_, ν_1_, ν_2_, ν_3_, ν_4_) describe the pseudo-rotation phase angle P [**40**]. Such multi dimensionality of the RNA conformational space represents severe hindrance to its systematic conformational analysis. Two new degrees of freedom called pseudo torsion angle □ and θ analogous to φ and ψ angle in protein are introduced to reduce the dimensionality of the RNA conformation [**41**].

Therefore the conformational fluctuations of apoA and apoB adenine riboswitch are quantified in terms of the distribution of microscopic conformational variables, □ and θ over the equilibrated trajectory. Pseudo torsion angle distributions corresponding to each base of apoA and apoB RNA are generated at different temperatures. Several nucleo-bases exhibit thermal sensitivity in □ and θ distribution. Few representative cases are shown in SI **(S5-S16)**.

Some representative histogram distribution of pseudo torsion angles of apoA and apoB at different temperatures are illustrated. Each peak of the histogram defines the most probable value for the corresponding variable. Increase in flexibility is observed due to the flat multimodal curves like 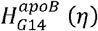 at stem1 at 313K, 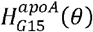 of stem1 and 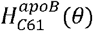 of loop3 at 323K. Sharper unimodal peaks are observed in 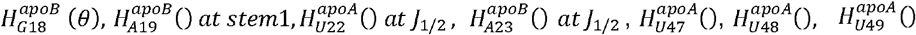 and 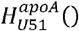 at *J*_2/3_, at temperature 283K, 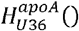 of loop2 at temperature 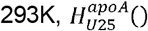 at 303K indicating rigidity. Bimodal distribution at 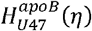 and 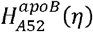 of *J*_2/3_ at 283K and 293K, 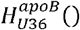 of loop2 at 323K, 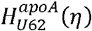 of loop3 at 323K suggest expansion of flexibility.

Alteration of the single peak into multiple peaked or broadened distribution under different thermal condition suggests enhanced flexibility in the given pseudo-torsion angle. Overall, the changes in pseudo torsion angles distribution of apoA and apoB at different temperatures indicate that stem1 and junction regions show significant flexibility at different temperatures while loop3 show sensitivity in θ distribution at 323K and 373K. Different rotameric states denoted by distribution of η and θ are prominent in junction region. The histogram of the bases of RNA for η and θ indicate that overall, the number of peaks, peak heights and widths are different in apoA and apoB states and typically sensitive to temperature.

### 3.4 Comparison of conformational stability and order of apo A with respect to apoB across varied temperatures in terms of conformational thermodynamics

We account for changes in free energy △G and entropy T△S of the apoA adenine riboswitch with respect to apoB adenine riboswitch from the distributions of the pseudo-torsion angles across varied temperature. Positive values of the changes in free energy and entropy indicate destabilization and disorder of the apoA adenine riboswitch in comparison to the apoB adenine riboswitch, while the negative values indicate conformational stabilization and order. The overall changes in conformational thermodynamics of the various domains of both apo adenine riboswitch are obtained by adding all the pseudo-torsion angles contributions from the residues of the particular region. Both the residue-wise (Table S1-S7) and the domain-wise (Table 5 (a) and (b)) free energy and entropy costs for conformational changes in the apoA with respect to the apoB under different thermal condition are computed.

**Table 5(a):** The changes (kJ/mol) in conformational thermodynamics of apoA adenine riboswitch in comparison to the apoB adenine riboswitch domain wise (based on pseudo torsion angle)

| Region | Stem1 |  | Stem2 |  | Stem3 |  |
| --- | --- | --- | --- | --- | --- | --- |
| Temperature | $\Delta G$ | $T\Delta S$ | $\Delta G$ | $T\Delta S$ | $\Delta G$ | $T\Delta S$ |
| 283K | 6.19 | 8.77 | -3.69 | -4.13 | 1.5 | 0.88 |
| 293K | -1.22 | -1.67 | -4.69 | -4.83 | -2.01 | -1.25 |
| 303K | 0.36 | -4.0 | -7.03 | -6.65 | 3.21 | 2.1 |
| 313K | 0.16 | 2.60 | -0.20 | -1.11 | 2.02 | 1.54 |
| 323K | 7.91 | 11.74 | -1.26 | -1.51 | -1.94 | 0.80 |
| 373K | -4.64 | 2.70 | -4.70 | -9.26 | 3.10 | 4.42 |
| Region | Loop2 |  | Loop3 |  |  |  |
| Temperature | $\Delta G$ | $T\Delta S$ | $\Delta G$ | $T\Delta S$ | | |
| 283K | -3.65 | -3.01 | -1.58 | -2.10 |  |  |
| 293K | -3.61 | -3.67 | -8.16 | -11.63 |  |  |
| 303K | -2.46 | -2.15 | -3.26 | -3.84 |  |  |
| 313K | -0.85 | -0.96 | 0.81 | 0.25 |  |  |
| 323K | -2.45 | -0.25 | 7.47 | 11.20 |  |  |
| 373K | -6.56 | -9.59 | 0.78 | 1.33 |  |  |
| Region | J1/2 (Hinge) |  | J2/3 (Latch) |  |  |  |

| Temperature | $\Delta G$ | $T\Delta S$ | $\Delta G$ | $T\Delta S$ |
| --- | --- | --- | --- | --- |
| 283K | -0.64 | -1.21 | -20.58 | -21.64 |
| 293K | -5.63 | -4.87 | -17.85 | -21.37 |
| 303K | -8.88 | -11.17 | -23.41 | -28.01 |
| 313K | -2.88 | -6.04 | -23.00 | -27.00 |
| 323K | -4.20 | -4.63 | -14.98 | -16.06 |
| 373K | 7.77 | 5.23 | -15.36 | -20.18 |

**Table 5(b):** The changes (kJ/mol) in conformational thermodynamics of apoA adenine riboswitch in comparison to the apoB adenine riboswitch (based on pseudo torsion angle) whole aptamer

| Temperature | $\Delta G$ | $T\Delta S$ |
| --- | --- | --- |
| 283K | -22.45 | -22.44 |
| 293K | -43.17 | -49.29 |
| 303K | -41.47 | -53.72 |
| 313K | -23.94 | -30.72 |
| 323K | -9.45 | 1.29 |
| 373K | -19.61 | -25.35 |

Stem1 of apoA with respect to apoB becomes energetically destabilized and disordered at 283K and 323K temperature. However, stem1 from the apoA remains marginally stabilized and ordered with respect to the apoB at 293K. The marginal instability and disorder is observed in the stem1 of apoA at 313K. In contrast, apoA of stem1 exhibit energetically stabilization and entropically disorderness at 373K.

It is observed that stem2 of the apoA remains energetically stabilized and ordered with reference to the apoB across the varied thermal condition. 303K temperature imparts maximum stability and order towards stem2 of the apoA whereas marginal conformational stability and order are observed at 313K and 323K.

The stem3 of the of the adenine riboswitch undergo disorder and destabilization in the apoA system compared to the apoB system at temperature 303K, 313K, and 373K. It is observed that the apoA system remains energetically stabilized and ordered with reference to the apoB system at temperature at 293K and 323K. It has been found that the overall entropy and free energy changes of the stem3 of apoA remain marginal compared to apoB.

The junction region J_2/3_ (Latch), loop2 and stem2 of apoA with respect to apoB impart substantial stability and order under different thermal condition.

It has been found that the overall entropy and free energy changes of the loop3 of apoA compared to apoB remain marginal at temperature 313K and 373K while for that of loop2, marginal stabilization and order is observed at 313K.

It may be noted that in the apoA system with respect to apoB adenine riboswitch, the J_1/2_ shows maximum disorder and destabilization at 373K whereas 283K temperature imparts marginal conformational stability and order. On the other hand J_1/2_ of apoA compared to apoB get maximum stabilization at 303K followed by 293K, 323K and 313K.

The results for changes in conformational thermodynamics are shown in Table 5. (b) suggest that conformational stability and order of apoA with respect to apoB adenine riboswitch conformation is significant at 293K and 303K. The temperatures corresponding to the conformational order and stability of apoA adenine riboswitch with respect to apoB riboswitch whole aptamer are shown in descending order 293K∼303K> 313K∼283K>373K>323K.

#### 3.4.1 Conformational thermodynamics changes of apo A with respect to apoB at Stem1

The total change in free energy and entropy of stem1 at 283K are 6.19 kJ/mol and 8.77 kJ/mol respectively. G14, A17, G18, A19, U75 and A76 impart enhanced destabilization and disorder, the maximum being in G18. The overall changes in the free energy and entropy of stem1 at 293K are marginal,-1.22 kJ/mol and-1.68 kJ/mol respectively. G14, G15, U20, A21, U75 confer marginal stability and conformational order, the maximum being observed in A21. On the other hand G14, G15, A16, A21 and U75 show stabilization and order of stem1 at 303K while A17, G18, A19 and A76 impart destabilization and disorder leading marginal change in free energy (ΔG=0.36 kJ/mol) and significant change of entropy (TΔS=-4.0 kJ/mol) of the stem1 at 303K. The marginal destabilization and disorder of stem1 at 313K are indicated by changes in the free energy 0.17 kJ/mol and entropy 2.61 kJ/mol. Almost all the bases of stem1 at 313K are responsible for decrease in stability and order except U20 and A21. It is observed that the total change in free energy and entropy of stem1 at 323K are 7.91 kJ/mol and 11.74 kJ/mol. Bases G14, G15, A17, G18, A19, U75 and A76 account for the decrease in stability and order, the highest destabilized and disordered base being G15. However conformational stability of stem1 at 373K is indicated by changes in the free energy -4.64 kJ/mol while disorderness is conferred by changes in the entropy 2.7 kJ/mol. Base G14 accounts for the maximum stability while G18 impart the highest disorderness.

#### 3.4.2 Conformational thermodynamics changes of apo A with respect to apoB at Stem2

Let us now discuss on conformational stability and order of stem2 across the varied temperatures. It is found that stem2 attains the maximum stability at 303K and maximum order at 373K. The total change in free energy and entropy of stem2 at 303K are -7.03 kJ/mol and -6.65 kJ/mol respectively. U25 confer a major increase in stability and order at 303K whereas in the rest of the bases of stem2, the change in free energy and entropy are marginal. The total change in free energy and entropy of stem2 at 373K are -4.70 kJ/mol and -9.26 kJ/mol. U31, U39 and U40 of stem2 undergo a maximum stability and order at 373K. Stem2 is marginally stable and ordered at 313K and 323K. The total change in free energy of Stem2 at 313K and 323K are -0.20 kJ/mol respectively and -1.26 kJ/mol whereas the total change in entropy are -1.11 kJ/mol and -1.51 kJ/mol respectively. U25 is responsible for the maximum stability and order. On the other hand, U39 and A45 of stem2 impart marginal increase in stability and order at 313K and 323K temperatures. The extent of stability and order of stem2 are nearly uniform at both 283K and 293K temperatures. -3.69 kJ/mol and -4.69 kJ/mol are the total change in free energy of stem2 at 283K and 293K temperatures respectively. Bases U28, A29 at 283K and U25 at 293K contribute enhanced stability and order in stem2.

#### 3.4.3 Conformational thermodynamics changes of apo A with respect to apoB at Stem3

Stem3 is marginally unstable and disordered across the varied temperature except at 293K. The total change in free energy of Stem2 at 293K are -2.01 kJ/mol and -1.25 kJ/mol. It is found that G57 and U68 account for the major stability and order of stem3 at 293K whereas in rest of the bases of stem3, the change of free energy and entropy are marginal. Stem3 shows maximum destabilization (ΔG=3.10 kJ/mol) and disorder (TΔS=4.42 kJ/mol) at 373K. 70U and 71U of stem3 undergo maximum destabilization and disorder at 373K.

#### 3.4.4 Conformational thermodynamics changes of apo A with respect to apoB at Loop2

Loop2 attains maximum stability (ΔG=-6.56 kJ/mol) and order (TΔS=-9.59 kJ/mol) at 373K. G32, A33, and G38 of loop2 at 373K exhibit an increase in order and stability. However, the changes in free energy (ΔG=-0.85 kJ/mol) and entropy (TΔS=-0.96 kJ/mol) of loop2 are marginal at 313K. The extent of stability (ΔG=-3.6 kJ/mol) and order (TΔS=-3.0 kJ/mol) of loop2 are nearly uniform at 283K and 293K. G32 and G37 impart maximum stability and order at 283K and 293K respectively. The extent of stability of loop2 are nearly equal at 303K (ΔG=-2.46 kJ/mol) and 323K (ΔG=-2.46 kJ/mol). It is found that G37 and U36 exhibit enhanced stability and order of loop2 at 303K and 323K respectively.

#### 3.4.5 Conformational thermodynamics changes of apo A with respect to apoB at Loop3

The total changes in free energy and entropy of loop3 at 293K are -8.16 kJ/mol and -11.63 kJ/mol respectively suggesting maximum conformational stability and order. C61, U62, U63, and A65 of loop3 impart enhanced stability and order, the maximum being U62 at 293K. However, loop3 is marginally unstable and disordered at 313K and 373K. In contrast, loop3 is marginally stable at 283K. The total change in free energy and entropy of loop3 at 303K are -3.26 kJ/mol and -3.84 kJ/mol. The stability and order of loop3 decreases with increase of temperature. The maximum destabilization (ΔG=7.47 kJ/mol) and disorder (TΔS=11.20 kJ/mol) of loop3 is observed at 323K. C61 and U62 of loop3 at 323K confer a major decrease in stability and order whereas in rest of the bases of loop3, the free energy and entropy change is marginal.

#### 3.4.6 Conformational thermodynamics changes of apo A with respect to apoB at J_1/2_ (Hinge)

Hinge J_1/2_ attains maximum stability and order at 303K. -8.88 kJ/mol and -11.2 kJ/mol are the total changes in free energy and entropy of J_1/2_ at 303K. Marginal stability (ΔG=-0.64 kJ/mol) and order (TΔS=-1.21 kJ/mol) are found in J_1/2_ at 283K. The extent of stability and order of J_1/2_ are nearly uniform at 293K (ΔG=-5.63 kJ/mol, TΔS= -4.87 kJ/mol) and 323K (ΔG=-4.20 kJ/mol, TΔS=-4.63 kJ/mol) respectively. The total change in free energy and entropy of J_1/2_ at 313K are -2.88 kJ/mol and -6.04 kJ/mol. On the other hand, maximum destabilization and disorder are observed in J_1/2_ at 373K, the highest destabilized and disordered base being A23. The total change in free energy and entropy of J_1/2_ at 373K are 7.77 kJ/mol and 5.23 kJ/mol.

#### 3.4.7 Conformational thermodynamics changes of apo A with respect to apoB at J_2/3_ (Latch)

Latch J_2/3_ is extremely stable and ordered among all domain of apo Adenine riboswitch under different thermal condition. The extent of conformational stability and order of J_2/3_ are nearly equal and maximum at both temperature 303K (ΔG= -23.41 kJ/mol, TΔS= -28.01 kJ/mol) and (ΔG= -23.00 kJ/mol, TΔS= -27.00 kJ/mol) 313K. Almost all the bases of J_2/3_ show enhanced stability and order, the maximum being U48 at both 303K and 313K. The total change in free energy and entropy of J_2/3_ at temperature 283K are -20.58 kJ/mol and -21.64 kJ/mol. U48 confer the conformational stability and order at 283K. However, the stability and order of J_2/3_ are decreased some extent at 323K (ΔG= -15.00 kJ/mol, TΔS= -16.06 kJ/mol), 373K (ΔG= -15.40 kJ/mol, TΔS= -20.18 kJ/mol) and 293K (ΔG= -17.85 kJ/mol, TΔS= -21.37 kJ/mol). 46G and 47U are responsible for slight decrease in stability and order of J_2/3_ at 373K. On the other hand U47, U48 and U49 impart stability and order of J_2/3_ at 323K. On the other hand G46, C50, and A52 are responsible for marginal increase in destabilization and disorder of J_2/3_ at 323K.

### 3.5 Principal component analysis (PCA) demonstrates different dominant motions in apo state

The practical application of PCA is to reduce the dimensionality of the huge data preserving the most of the variation in the data. Essential dynamics analysis (EDA) is utilized here to study the essential motion of the sampled conformations generated from molecular dynamics trajectory among seven different temperatures. If a riboswitch displays large conformational changes, then the distributions are also scattered in the conformational space. It is observed from EDA (Figure 2.) that both the apoA and apoB form of adenine riboswitch’s distribution are more widely dispersed at elevated temperatures 400K, 373K and 323K. However, the distribution of ligand unbound states of adenine riboswitch in the conformational space is limited at physiological (303K and 313K) and low temperatures (283K and 293K).

**Figure 2.**
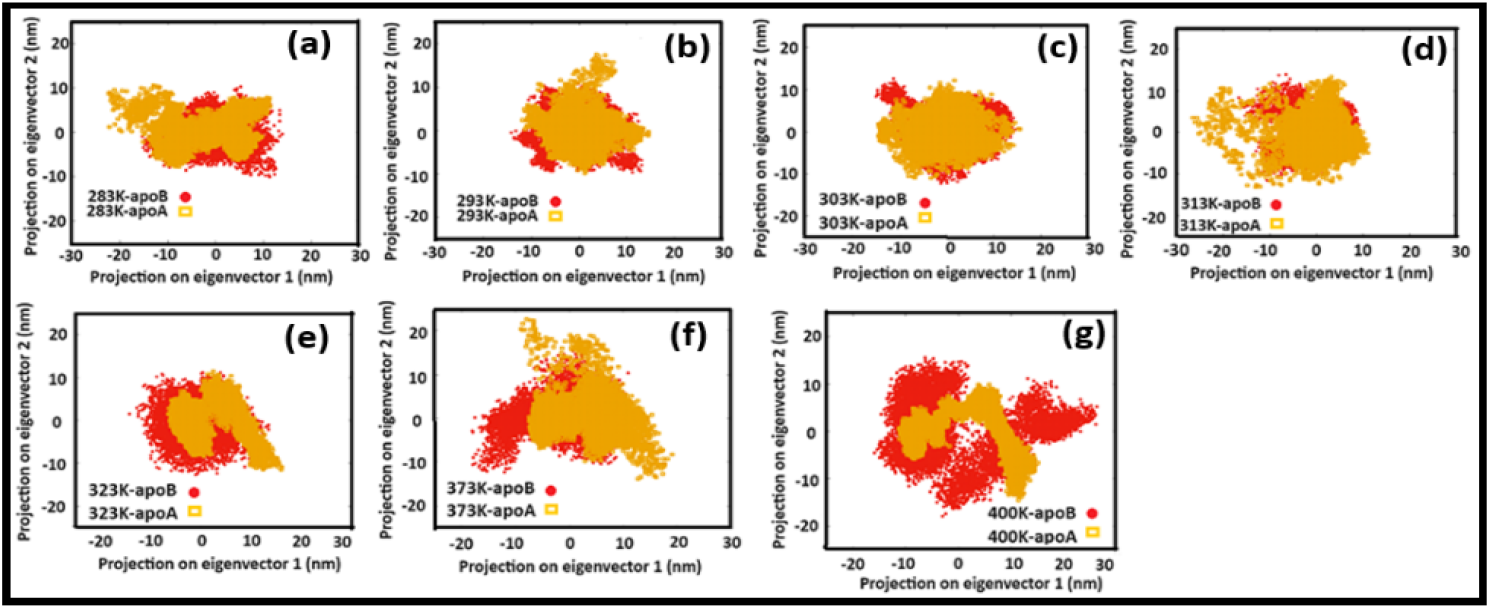
2D projection of simulated trajectories of apoA (yellow) and apoB (red) Adenine riboswitch

Figure 2. represents that apoB occupies larger and distinct conformational subspace than apoA at 323K and 400K temperature pointing toward the change in the conformational subspace of the riboswitch due to temperature. On the other hand, apoB occupies smaller conformational subspace than apoA at 283K and 313K (Figure 2.).

To evaluate the contrast in the structure and dynamics of the apoA adenine riboswitch in comparison to apoB adenine riboswitch, comparative dynamic cross correlation matrix (DCCM) analysis is carried out at different temperatures to figure out the correlation or anti correlation between nucleotides of the different regions of ligand unound adenine riboswitch (Figure S4). Correlation coefficients ranges between the value +1 and -1 representing both positive and negative correlation reflected from colour gradients. The blue colour suggests positive correlation, pink represents negative correlation and white indicates uncorrelated fluctuations.

Junction J_2/3_ shows positively correlated movements with stem1 for apoA and apoB at 283K as well as apoA at 303K illustrating the nucleotides are hydrogen bonded and tend to move together. Correlated motion of nucleotides between nucleotides of apoA and apoB Adenine riboswitches decrease significantly with increase of temperatures suggesting interaction decreases with increase of temperatures.

The x- and y-axis represent the first two eigenvectors, PC1 and PC2, of the P atomic fluctuation. The z-axis represents the free energy in kJ mol^−1^. The color conventions are depicted as red (energy maxima) and blue (energy minima). The first two principal components, PC1 and PC2 of apoA and apoB Adenine riboswitch are employed to obtain information on the global minimum-energy (stable) conformations responsible for the essential dynamics, at different temperatures, which are depicted in the form of Gibbs free energy landscapes (FEL) shown in Figure 3.

**Figure 3.**
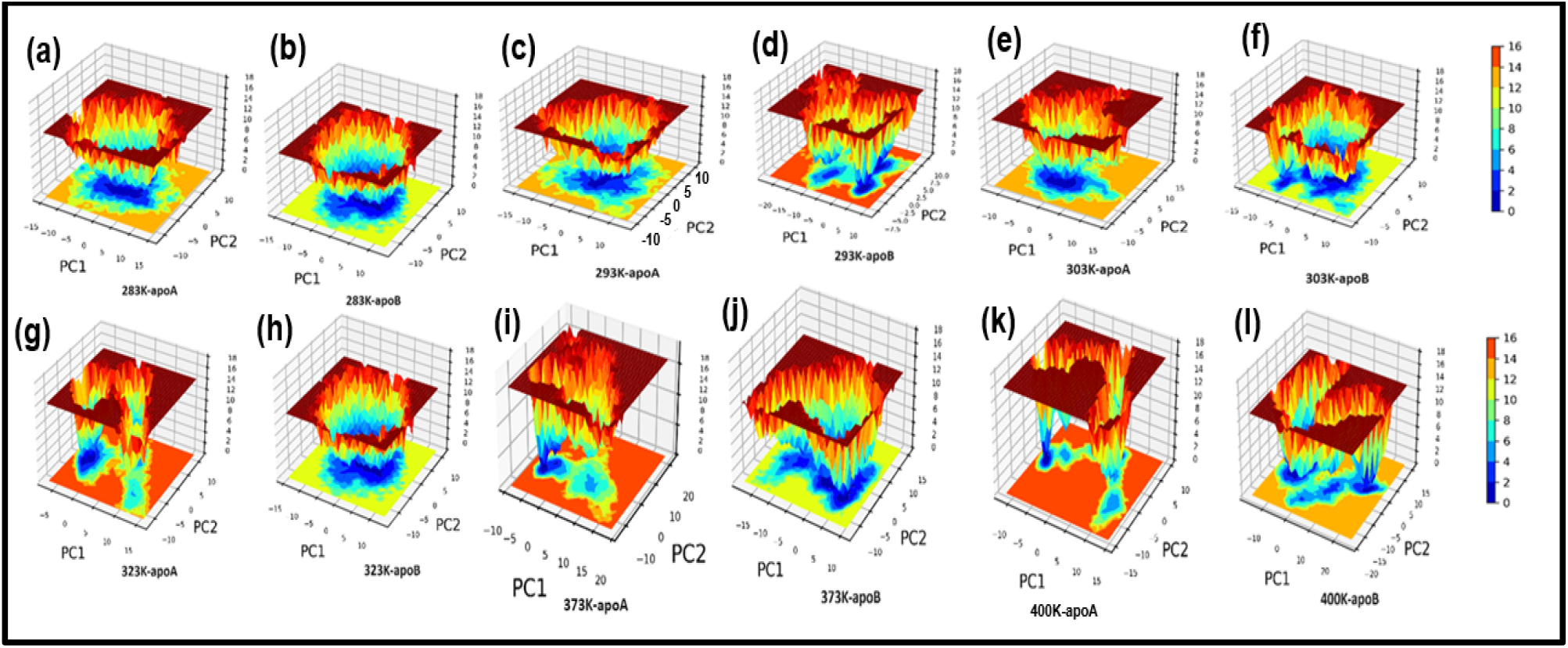
Gibbs free energy landscapes of PC1 & PC2 (initial two principal components) of projection of simulated trajectories apoA and apoB adenine riboswitches

The lowest energy states or stable conformations represented by blue coloured maps whereas red coloured maps corresponded to high-energy conformations. Each riboswitch produced different FEL pattern at different temperatures. FEL of the riboswitch (apoA and apoB at 283K, apoA 293K) depicted broader minima with little energy barrier suggesting large number of stable conformations. Single dominant basins (apoA at 303K) in the FEL plots indicate stable form while multiple basins in the FEL indicate destabilization (apoB at 293K, apoA and apoB at 400K). Minima with flat ends indicated clustering of the stable or low energy conformations.

The shallow and narrow energy basin observed during the simulation revealed the low stability of apoA at 323K and 373K. The stable conformations indicated by the minima are responsible for essential dynamics in the riboswitch, and as the number of stable conformations decreases flexibility of the system increases and vice versa. Consequently, FEL pattern served to figure out the effect of the temperatures on the sampled essential subspace.

## 4. Conclusion

In summary, the result of my current research work suggests that temperature induces conformational changes on the two different structures of apo adenine riboswitch (possessing the same sequence) that remodel the internal hydrogen bonding network as well as affect the conformational stability and order. It is observed from the conformational thermodynamics view that temperature affects different core regions of the apo adenine riboswitch diversely. Nucleotides of the stem2 and junction regions (**J**_**1/2**_, **J**_**2/3**_) contribute the most conformational stability and order whereas stem1 contributes the least conformational stability and order towards apo adenine riboswitch. The conformational stability of apoA is more pronounced compared to apoB adenine riboswitch at low temperatures (293K and 303K). Thermal stability of apoA compared to apoB adenine riboswitch is the least at 323K temperature. At higher temperature (373K) and lower temperature (283K) apoA adenine riboswitch exerts moderate conformational stability.

Such conformational thermodynamics study based on the probability distribution of proper conformational variable delineates a new approach to explore the stability and order of the variable ribo nucleobase geometry of the riboswitch both residue-wise and region-wise due to thermal variation. Understanding the intrinsic relationships among basic structural motifs of apo adenine riboswitch may be beneficial to elucidate structural dynamics and function as well as unravel the stability and dynamics folding of the full length RNA. The distinct thermo-sensitivity of adenine riboswitch suggested in this work may contribute to the research field of thermometer RNA and RNA sensors.

## Author Contributions

Soumi Das conceptualized, curated, analyzed, interpreted the data, and wrote the manuscript.

## Acknowledgments

Soumi Das is thankful to the Technical Research Centre, S. N. Bose National Centre for Basic Sciences, Kolkata for the computational facilities, and the Department of Science and Technology (DST) for funding.

