## Supplementary Information for "Decoding the effect of temperatures on conformational stability and order of ligand unbound thermo sensing adenine riboswitch using molecular dynamics simulation"

**Soumi Das^1*^**

**^1^Department of Physics of Complex Systems, S. N. Bose National Centre for Basic Sciences, Block-JD, Sector-III, Salt Lake, Kolkata 700106, India.**

**Correspondence: Soumi Das**

**Supplementary figure:**

**Figure S1: MD simulation average snapshots for apoA and apoB adenine riboswitch at different temperatures**

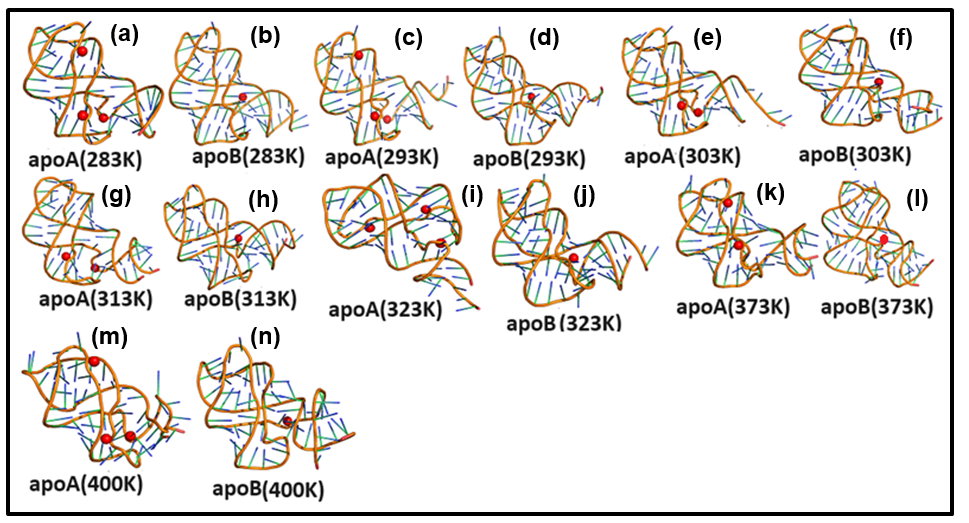

**Figure S2: Root mean square deviation (RMSD) of backbone atoms of apoA (violet) and apoB (green) adenine riboswitch as a function of simulation time 1µs at different temperatures considering cryatallographic structure as a reference structure.**

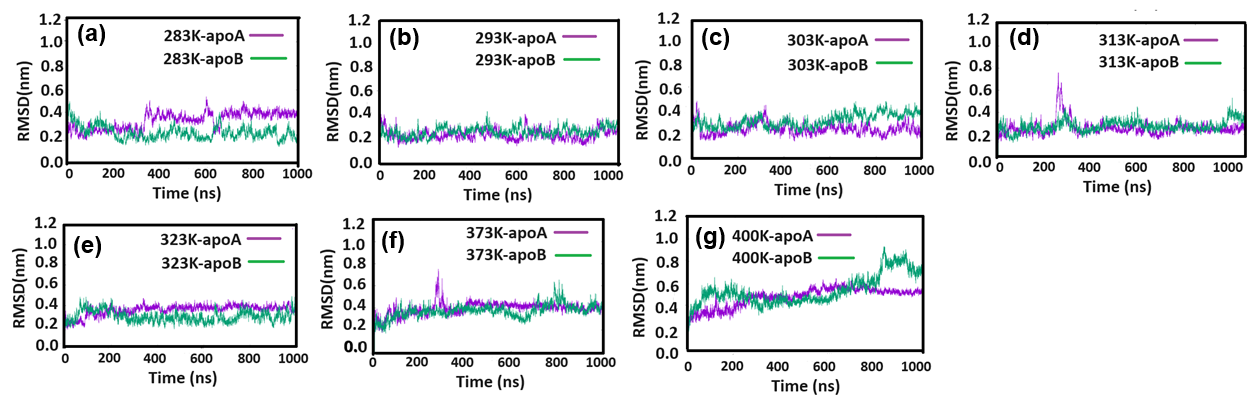

**Figure S3: Root mean square fluctuation (RMSF) of simulated systems apoA (red) and apoB (blue) adenine riboswitch at different temperatures**

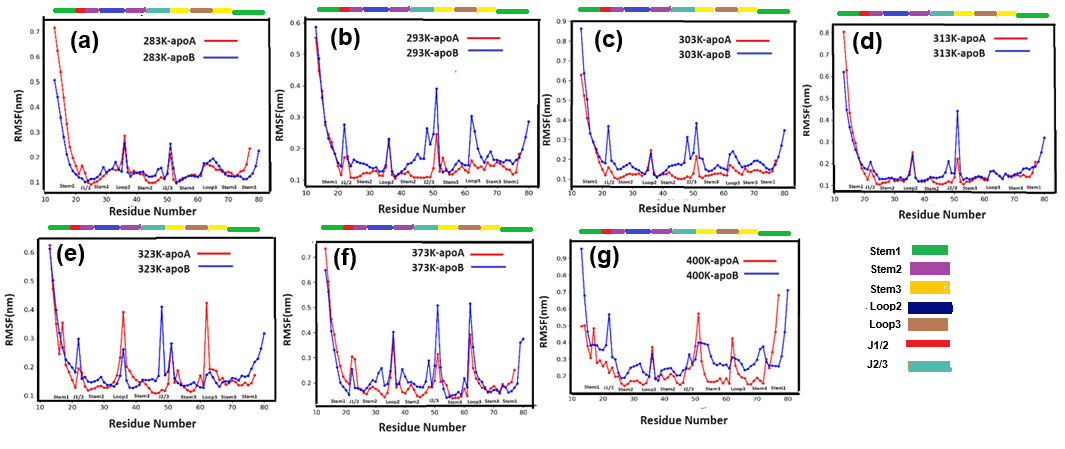

**Figure S4: Dynamic cross correlation matrix (DCCM) of backbone atoms of apoA (red) and apoB (blue) adenine riboswitch at different temperatures**

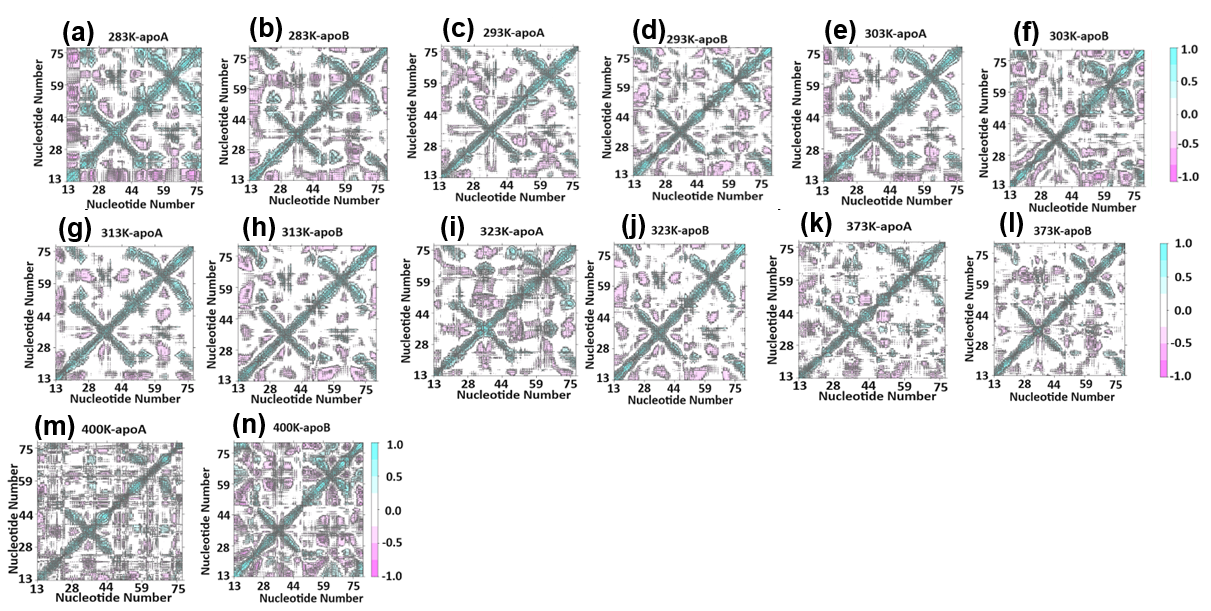

**Figure S5: Some representative Histogram distribution of pseudo-torsion angle (η) at 283K (apoA in blue and apoB in orange)**

(a) 283K-apoAB-eta-A19-Stem1

H(η)

| η | NF-apoA | NF-apoB |
| --- | --- | --- |
| 0 | 0 | 0 |
| 5 | 0 | 0 |
| 10 | 0 | 0 |
| 15 | 0 | 0 |
| 20 | 0 | 0 |
| 25 | 0 | 0 |
| 30 | 0 | 0 |
| 35 | 0 | 0 |
| 40 | 0 | 0 |
| 45 | 0 | 0 |
| 50 | 0 | 0 |
| 55 | 0 | 0 |
| 60 | 0 | 0 |
| 65 | 0 | 0 |
| 70 | 0 | 0 |
| 75 | 0 | 0 |
| 80 | 0 | 0 |
| 85 | 0 | 0 |
| 90 | 0 | 0 |
| 95 | 0 | 0 |
| 100 | 0 | 0 |
| 105 | 0 | 0 |
| 110 | 0 | 0 |
| 115 | 0 | 0 |
| 120 | 0 | 0 |
| 125 | 9.999e-05 | 0 |
| 130 | 9.999e-05 | 0 |
| 135 | 0.00059994 | 0 |
| 140 | 0.00519948 | 0 |
| 145 | 0.019898 | 0 |
| 150 | 0.0709929 | 0.00039996 |
| 155 | 0.144086 | 0.00459954 |
| 160 | 0.19608 | 0.0260974 |
| 165 | 0.187281 | 0.0971903 |
| 170 | 0.123188 | 0.212479 |
| 175 | 0.0535946 | 0.284472 |
| 180 | 0.0241976 | 0.232777 |
| 185 | 0.0143986 | 0.106789 |
| 190 | 0.0140986 | 0.0282972 |
| 195 | 0.0147985 | 0.00649935 |
| 200 | 0.0208979 | 0.00039996 |
| 205 | 0.0258974 | 0 |
| 210 | 0.0286971 | 0 |
| 215 | 0.0243976 | 0 |
| 220 | 0.0161984 | 0 |
| 225 | 0.010099 | 0 |
| 230 | 0.00339966 | 0 |
| 235 | 0.00089991 | 0 |
| 240 | 0.00079992 | 0 |
| 245 | 9.999e-05 | 0 |
| 250 | 0 | 0 |
| 255 | 0 | 0 |
| 260 | 0 | 0 |
| 265 | 0 | 0 |
| 270 | 0 | 0 |
| 275 | 0 | 0 |
| 280 | 0 | 0 |
| 285 | 0 | 0 |
| 290 | 0 | 0 |
| 295 | 0 | 0 |
| 300 | 0 | 0 |
| 305 | 0 | 0 |
| 310 | 0 | 0 |
| 315 | 0 | 0 |
| 320 | 0 | 0 |
| 325 | 0 | 0 |
| 330 | 0 | 0 |
| 335 | 0 | 0 |
| 340 | 0 | 0 |
| 345 | 0 | 0 |
| 350 | 0 | 0 |
| 355 | 0 | 0 |
| 360 | 0 | 0 |

(b) 283K-apoAB-eta-A21-Stem1

H(η)

| η | NF-apoA | NF-apoB |
| --- | --- | --- |
| 0 | 0 | 0 |
| 5 | 0 | 0 |
| 10 | 0 | 0 |
| 15 | 0 | 0 |
| 20 | 0 | 0 |
| 25 | 0 | 0 |
| 30 | 0 | 0 |
| 35 | 0 | 0 |
| 40 | 0 | 0 |
| 45 | 0 | 0 |
| 50 | 0 | 0 |
| 55 | 0 | 0 |
| 60 | 0 | 0 |
| 65 | 0 | 0 |
| 70 | 0 | 0 |
| 75 | 0 | 0 |
| 80 | 0 | 0 |
| 85 | 0 | 0 |
| 90 | 0 | 0 |
| 95 | 0 | 9.999e-05 |
| 100 | 0 | 0 |
| 105 | 0 | 9.999e-05 |
| 110 | 0 | 0.00029997 |
| 115 | 0 | 0.00039996 |
| 120 | 0 | 0.00039996 |
| 125 | 0 | 0.00069993 |
| 130 | 0 | 0.00069993 |
| 135 | 0 | 0.00079992 |
| 140 | 0 | 0.00089991 |
| 145 | 9.999e-05 | 0.00069993 |
| 150 | 0 | 0.0009999 |
| 155 | 0.00069993 | 0.00089991 |
| 160 | 0.00829917 | 0.00169983 |
| 165 | 0.0461954 | 0.00279972 |
| 170 | 0.166383 | 0.00809919 |
| 175 | 0.306169 | 0.0368963 |
| 180 | 0.30487 | 0.115288 |
| 185 | 0.135086 | 0.213579 |
| 190 | 0.0292971 | 0.242976 |
| 195 | 0.00279972 | 0.177982 |
| 200 | 9.999e-05 | 0.107089 |
| 205 | 0 | 0.0542946 |
| 210 | 0 | 0.0225977 |
| 215 | 0 | 0.00839916 |
| 220 | 0 | 0.00119988 |
| 225 | 0 | 9.999e-05 |
| 230 | 0 | 0 |
| 235 | 0 | 0 |
| 240 | 0 | 0 |
| 245 | 0 | 0 |
| 250 | 0 | 0 |
| 255 | 0 | 0 |
| 260 | 0 | 0 |
| 265 | 0 | 0 |
| 270 | 0 | 0 |
| 275 | 0 | 0 |
| 280 | 0 | 0 |
| 285 | 0 | 0 |
| 290 | 0 | 0 |
| 295 | 0 | 0 |
| 300 | 0 | 0 |
| 305 | 0 | 0 |
| 310 | 0 | 0 |
| 315 | 0 | 0 |
| 320 | 0 | 0 |
| 325 | 0 | 0 |
| 330 | 0 | 0 |
| 335 | 0 | 0 |
| 340 | 0 | 0 |
| 345 | 0 | 0 |
| 350 | 0 | 0 |
| 355 | 0 | 0 |
| 360 | 0 | 0 |

(c) 283K-apoAB-eta-U22-J1/2

H(η)

| η | NF-apoA | NF-apoB |
| --- | --- | --- |
| 0 | 0 | 0.00039996 |
| 5 | 0 | 0.00779922 |
| 10 | 0 | 0.00189981 |
| 15 | 0 | 0.00049995 |
| 20 | 0 | 0.00019998 |
| 25 | 0 | 9.999e-05 |
| 30 | 0 | 0 |
| 35 | 0 | 0 |
| 40 | 0 | 9.999e-05 |
| 45 | 0 | 0.00029997 |
| 50 | 0 | 0.00019998 |
| 55 | 0 | 0 |
| 60 | 0 | 0 |
| 65 | 0 | 0 |
| 70 | 0 | 0 |
| 75 | 0 | 0 |
| 80 | 0 | 0 |
| 85 | 0 | 0 |
| 90 | 0 | 0 |
| 95 | 0 | 0 |
| 100 | 0 | 0 |
| 105 | 0 | 0 |
| 110 | 0 | 0 |
| 115 | 0 | 0 |
| 120 | 0 | 0 |
| 125 | 0 | 0 |
| 130 | 0 | 0 |
| 135 | 0 | 0 |
| 140 | 0 | 0 |
| 145 | 0 | 0 |
| 150 | 0 | 0 |
| 155 | 0 | 0 |
| 160 | 0 | 0 |
| 165 | 0 | 0 |
| 170 | 0 | 0 |
| 175 | 0 | 0 |
| 180 | 0 | 0 |
| 185 | 0 | 0 |
| 190 | 0 | 0 |
| 195 | 0 | 0 |
| 200 | 0 | 0 |
| 205 | 9.999e-05 | 0 |
| 210 | 0.00369963 | 0 |
| 215 | 0.0559944 | 0 |
| 220 | 0.236776 | 0 |
| 225 | 0.369363 | 0 |
| 230 | 0.237576 | 0 |
| 235 | 0.079692 | 0 |
| 240 | 0.0142986 | 0 |
| 245 | 0.00239976 | 0 |
| 250 | 9.999e-05 | 0 |
| 255 | 0 | 0 |
| 260 | 0 | 0 |
| 265 | 0 | 0 |
| 270 | 0 | 0 |
| 275 | 0 | 0 |
| 280 | 0 | 0 |
| 285 | 0 | 0 |
| 290 | 0 | 0 |
| 295 | 0 | 0 |
| 300 | 0 | 0 |
| 305 | 0 | 0 |
| 310 | 0 | 0.00079992 |
| 315 | 0 | 0.00109989 |
| 320 | 0 | 0.00239976 |
| 325 | 0 | 0.00459954 |
| 330 | 0 | 0.0235976 |
| 335 | 0 | 0.0991901 |
| 340 | 0 | 0.229977 |
| 345 | 0 | 0.29987 |
| 350 | 0 | 0.19958 |
| 355 | 0 | 0.0941906 |
| 360 | 0 | 0.0331967 |

(d) 283K-apoAB-eta-A23-J1/2

H(η)

| η | NF-apoA | NF-apoB |
| --- | --- | --- |
| 0 | 0 | 0 |
| 5 | 0 | 0 |
| 10 | 0 | 0 |
| 15 | 0 | 0 |
| 20 | 0 | 0 |
| 25 | 0 | 0 |
| 30 | 0 | 0 |
| 35 | 0 | 0 |
| 40 | 0 | 0 |
| 45 | 0 | 0 |
| 50 | 0 | 0 |
| 55 | 0 | 0 |
| 60 | 0 | 0 |
| 65 | 0 | 0 |
| 70 | 0 | 0 |
| 75 | 0 | 0 |
| 80 | 0 | 0 |
| 85 | 0 | 0 |
| 90 | 0 | 0 |
| 95 | 0 | 0 |
| 100 | 0 | 0 |
| 105 | 0 | 0 |
| 110 | 0 | 0 |
| 115 | 0 | 0 |
| 120 | 0 | 0 |
| 125 | 0 | 0 |
| 130 | 0 | 0 |
| 135 | 0 | 0 |
| 140 | 0 | 0 |
| 145 | 0 | 0 |
| 150 | 0 | 0 |
| 155 | 0 | 0 |
| 160 | 0 | 0 |
| 165 | 0.00029997 | 0 |
| 170 | 0.00629937 | 0 |
| 175 | 0.0329967 | 0 |
| 180 | 0.0905909 | 0 |
| 185 | 0.169683 | 0 |
| 190 | 0.206979 | 0 |
| 195 | 0.228077 | 0 |
| 200 | 0.165883 | 0 |
| 205 | 0.0782922 | 0 |
| 210 | 0.0176982 | 0 |
| 215 | 0.0029997 | 0 |
| 220 | 0.00019998 | 0 |
| 225 | 0 | 0 |
| 230 | 0 | 0 |
| 235 | 0 | 0 |
| 240 | 0 | 0 |
| 245 | 0 | 0.00019998 |
| 250 | 0 | 0.00349965 |
| 255 | 0 | 0.0249975 |
| 260 | 0 | 0.121488 |
| 265 | 0 | 0.290771 |
| 270 | 0 | 0.343666 |
| 275 | 0 | 0.171183 |
| 280 | 0 | 0.0352965 |
| 285 | 0 | 0.00349965 |
| 290 | 0 | 0.00229977 |
| 295 | 0 | 0.00119988 |
| 300 | 0 | 0.00129987 |
| 305 | 0 | 0.00039996 |
| 310 | 0 | 0.00019998 |
| 315 | 0 | 0 |
| 320 | 0 | 0 |
| 325 | 0 | 0 |
| 330 | 0 | 0 |
| 335 | 0 | 0 |
| 340 | 0 | 0 |
| 345 | 0 | 0 |
| 350 | 0 | 0 |
| 355 | 0 | 0 |
| 360 | 0 | 0 |

(e) 283K-apoAB-eta-A24-J1/2

H(η)

| η | NF-apoA | NF-apoB |
| --- | --- | --- |
| 0 | 0 | 0 |
| 5 | 0 | 0 |
| 10 | 0 | 0 |
| 15 | 0 | 0 |
| 20 | 0 | 0 |
| 25 | 0 | 0 |
| 30 | 0 | 0 |
| 35 | 0 | 0 |
| 40 | 0 | 0 |
| 45 | 0 | 0 |
| 50 | 0 | 0 |
| 55 | 0 | 0 |
| 60 | 0 | 0 |
| 65 | 0 | 0 |
| 70 | 0 | 0 |
| 75 | 0 | 0 |
| 80 | 0 | 0 |
| 85 | 0 | 0 |
| 90 | 0 | 0 |
| 95 | 0 | 0 |
| 100 | 0 | 0 |
| 105 | 0 | 0 |
| 110 | 0 | 0 |
| 115 | 0 | 0 |
| 120 | 0 | 0 |
| 125 | 0 | 0 |
| 130 | 0 | 0 |
| 135 | 0 | 0 |
| 140 | 0 | 0 |
| 145 | 0 | 0 |
| 150 | 0 | 0 |
| 155 | 0 | 0 |
| 160 | 0 | 0 |
| 165 | 0 | 0 |
| 170 | 0 | 0 |
| 175 | 0 | 0.00029997 |
| 180 | 0 | 0.00129987 |
| 185 | 0 | 0.00329967 |
| 190 | 0 | 0.00329967 |
| 195 | 0 | 0.0112989 |
| 200 | 0 | 0.0551945 |
| 205 | 0 | 0.167983 |
| 210 | 0 | 0.277472 |
| 215 | 0 | 0.277272 |
| 220 | 0 | 0.145985 |
| 225 | 0 | 0.0435956 |
| 230 | 0 | 0.00969903 |
| 235 | 0 | 0.00249975 |
| 240 | 0.00049995 | 0.00049995 |
| 245 | 0.00419958 | 0.00029997 |
| 250 | 0.0277972 | 0 |
| 255 | 0.114089 | 0 |
| 260 | 0.276272 | 0 |
| 265 | 0.313869 | 0 |
| 270 | 0.183482 | 0 |
| 275 | 0.0648935 | 0 |
| 280 | 0.0128987 | 0 |
| 285 | 0.00179982 | 0 |
| 290 | 9.999e-05 | 0 |
| 295 | 9.999e-05 | 0 |
| 300 | 0 | 0 |
| 305 | 0 | 0 |
| 310 | 0 | 0 |
| 315 | 0 | 0 |
| 320 | 0 | 0 |
| 325 | 0 | 0 |
| 330 | 0 | 0 |
| 335 | 0 | 0 |
| 340 | 0 | 0 |
| 345 | 0 | 0 |
| 350 | 0 | 0 |
| 355 | 0 | 0 |
| 360 | 0 | 0 |

(f) 283K-apoAB-eta-U25-Stem2

H(η)

| η | NF-apoA | NF-apoB |
| --- | --- | --- |
| 0 | 0 | 0 |
| 5 | 0 | 0 |
| 10 | 0 | 0 |
| 15 | 0 | 0 |
| 20 | 0 | 0 |
| 25 | 0 | 0 |
| 30 | 0 | 0 |
| 35 | 0 | 0 |
| 40 | 0 | 0 |
| 45 | 0 | 0 |
| 50 | 0 | 0 |
| 55 | 0 | 0 |
| 60 | 0 | 0 |
| 65 | 0 | 0 |
| 70 | 0 | 0 |
| 75 | 0 | 0 |
| 80 | 0 | 0 |
| 85 | 0 | 0 |
| 90 | 0 | 0 |
| 95 | 0 | 0 |
| 100 | 0 | 0 |
| 105 | 0 | 0 |
| 110 | 0 | 0 |
| 115 | 0 | 0 |
| 120 | 0 | 0 |
| 125 | 0 | 0 |
| 130 | 0 | 0 |
| 135 | 0 | 0 |
| 140 | 0 | 0 |
| 145 | 0 | 0 |
| 150 | 0.00179982 | 0 |
| 155 | 0.020398 | 0 |
| 160 | 0.118188 | 0 |
| 165 | 0.278572 | 0.00019998 |
| 170 | 0.318468 | 0.0009999 |
| 175 | 0.188381 | 0.00269973 |
| 180 | 0.0612939 | 0.00839916 |
| 185 | 0.0106989 | 0.0232977 |
| 190 | 0.00169983 | 0.0477952 |
| 195 | 0.00049995 | 0.109189 |
| 200 | 0 | 0.244376 |
| 205 | 0 | 0.312669 |
| 210 | 0 | 0.181382 |
| 215 | 0 | 0.0576942 |
| 220 | 0 | 0.00879912 |
| 225 | 0 | 0.00209979 |
| 230 | 0 | 0.00039996 |
| 235 | 0 | 0 |
| 240 | 0 | 0 |
| 245 | 0 | 0 |
| 250 | 0 | 0 |
| 255 | 0 | 0 |
| 260 | 0 | 0 |
| 265 | 0 | 0 |
| 270 | 0 | 0 |
| 275 | 0 | 0 |
| 280 | 0 | 0 |
| 285 | 0 | 0 |
| 290 | 0 | 0 |
| 295 | 0 | 0 |
| 300 | 0 | 0 |
| 305 | 0 | 0 |
| 310 | 0 | 0 |
| 315 | 0 | 0 |
| 320 | 0 | 0 |
| 325 | 0 | 0 |
| 330 | 0 | 0 |
| 335 | 0 | 0 |
| 340 | 0 | 0 |
| 345 | 0 | 0 |
| 350 | 0 | 0 |
| 355 | 0 | 0 |
| 360 | 0 | 0 |

(g) 283K-apoAB-eta-U36-Loop2

H(η)

| η | NF-apoA | NF-apoB |
| --- | --- | --- |
| 0 | 0.00379962 | 0.00239976 |
| 5 | 0.192481 | 0.121788 |
| 10 | 0.179582 | 0.10019 |
| 15 | 0.129587 | 0.0667933 |
| 20 | 0.0780922 | 0.0752925 |
| 25 | 0.0437956 | 0.0944906 |
| 30 | 0.0182982 | 0.112589 |
| 35 | 0.0113989 | 0.106589 |
| 40 | 0.00429957 | 0.0769923 |
| 45 | 0.0019998 | 0.0375962 |
| 50 | 0.00119988 | 0.0111989 |
| 55 | 0.00019998 | 0.00179982 |
| 60 | 9.999e-05 | 0.00039996 |
| 65 | 0 | 0 |
| 70 | 0 | 0 |
| 75 | 0 | 0 |
| 80 | 0 | 0 |
| 85 | 0 | 0 |
| 90 | 0 | 0 |
| 95 | 0 | 0 |
| 100 | 0 | 0 |
| 105 | 0 | 0 |
| 110 | 0 | 0 |
| 115 | 0 | 0 |
| 120 | 0 | 0 |
| 125 | 0 | 0 |
| 130 | 0 | 0 |
| 135 | 0 | 0 |
| 140 | 0 | 0 |
| 145 | 0 | 0 |
| 150 | 0 | 0 |
| 155 | 0 | 0 |
| 160 | 0 | 0 |
| 165 | 0 | 0 |
| 170 | 0 | 0 |
| 175 | 0 | 0 |
| 180 | 0 | 0 |
| 185 | 0 | 0 |
| 190 | 0 | 0 |
| 195 | 0 | 0 |
| 200 | 0 | 0 |
| 205 | 0 | 0 |
| 210 | 0 | 0 |
| 215 | 0 | 0 |
| 220 | 0 | 0 |
| 225 | 0 | 0 |
| 230 | 0 | 0 |
| 235 | 0 | 0 |
| 240 | 0 | 0 |
| 245 | 0 | 0 |
| 250 | 0 | 0 |
| 255 | 0 | 0 |
| 260 | 0 | 0 |
| 265 | 0 | 0 |
| 270 | 0 | 0 |
| 275 | 0 | 0 |
| 280 | 0 | 0 |
| 285 | 0 | 0 |
| 290 | 0 | 0 |
| 295 | 0 | 0 |
| 300 | 0 | 0 |
| 305 | 0 | 0 |
| 310 | 0 | 0 |
| 315 | 0 | 0 |
| 320 | 0 | 0 |
| 325 | 0 | 0 |
| 330 | 0.00029997 | 0 |
| 335 | 0.00159984 | 0.00029997 |
| 340 | 0.0059994 | 0.00079992 |
| 345 | 0.0175982 | 0.0039996 |
| 350 | 0.0533947 | 0.0211979 |
| 355 | 0.10069 | 0.0581942 |
| 360 | 0.155584 | 0.107389 |

(h) 283K-apoAB-eta-G37-Loop2

H(η)

| η | NF-apoA | NF-apoB |
| --- | --- | --- |
| 0 | 0 | 0 |
| 5 | 0 | 0 |
| 10 | 0 | 0 |
| 15 | 0 | 0 |
| 20 | 0 | 0 |
| 25 | 0 | 0 |
| 30 | 0 | 0 |
| 35 | 0 | 0 |
| 40 | 0 | 0 |
| 45 | 0 | 0 |
| 50 | 0 | 0 |
| 55 | 0 | 0 |
| 60 | 0 | 0 |
| 65 | 0 | 0 |
| 70 | 0 | 0 |
| 75 | 0 | 0 |
| 80 | 0 | 0 |
| 85 | 0 | 0 |
| 90 | 0 | 0 |
| 95 | 0 | 0 |
| 100 | 0 | 0 |
| 105 | 0 | 0 |
| 110 | 0 | 0 |
| 115 | 0 | 0 |
| 120 | 0 | 0 |
| 125 | 0 | 0 |
| 130 | 0 | 9.999e-05 |
| 135 | 0 | 9.999e-05 |
| 140 | 0 | 0.0009999 |
| 145 | 0 | 0.00259974 |
| 150 | 0 | 0.00869913 |
| 155 | 0 | 0.0224978 |
| 160 | 0 | 0.0533947 |
| 165 | 0.00049995 | 0.0962904 |
| 170 | 0.00449955 | 0.136986 |
| 175 | 0.0157984 | 0.163884 |
| 180 | 0.0443956 | 0.169083 |
| 185 | 0.0825917 | 0.161984 |
| 190 | 0.137386 | 0.10409 |
| 195 | 0.174383 | 0.0550945 |
| 200 | 0.192281 | 0.0190981 |
| 205 | 0.167283 | 0.00479952 |
| 210 | 0.107789 | 0.00029997 |
| 215 | 0.0524948 | 0 |
| 220 | 0.0178982 | 0 |
| 225 | 0.00229977 | 0 |
| 230 | 0.00039996 | 0 |
| 235 | 0 | 0 |
| 240 | 0 | 0 |
| 245 | 0 | 0 |
| 250 | 0 | 0 |
| 255 | 0 | 0 |
| 260 | 0 | 0 |
| 265 | 0 | 0 |
| 270 | 0 | 0 |
| 275 | 0 | 0 |
| 280 | 0 | 0 |
| 285 | 0 | 0 |
| 290 | 0 | 0 |
| 295 | 0 | 0 |
| 300 | 0 | 0 |
| 305 | 0 | 0 |
| 310 | 0 | 0 |
| 315 | 0 | 0 |
| 320 | 0 | 0 |
| 325 | 0 | 0 |
| 330 | 0 | 0 |
| 335 | 0 | 0 |
| 340 | 0 | 0 |
| 345 | 0 | 0 |
| 350 | 0 | 0 |
| 355 | 0 | 0 |
| 360 | 0 | 0 |

(i) 283K-apoAB-eta-U47-J2/3

H(η)

|  | NF-apoA | NF-apoB |
| --- | --- | --- |
| 0 | 0 | 0 |
| 5 | 0 | 0 |
| 10 | 0 | 0 |
| 15 | 0 | 0 |
| 20 | 0 | 0 |
| 25 | 0 | 0 |
| 30 | 0 | 0 |
| 35 | 0 | 0 |
| 40 | 0 | 0 |
| 45 | 0 | 0 |
| 50 | 0 | 0 |
| 55 | 0 | 0 |
| 60 | 0 | 0 |
| 65 | 0 | 0 |
| 70 | 0 | 0 |
| 75 | 0 | 0 |
| 80 | 0 | 0 |
| 85 | 0 | 0 |
| 90 | 0 | 0 |
| 95 | 0 | 0 |
| 100 | 0 | 0 |
| 105 | 0 | 0 |
| 110 | 0 | 0 |
| 115 | 0 | 0 |
| 120 | 0 | 0 |
| 125 | 0 | 0 |
| 130 | 0 | 0 |
| 135 | 0 | 0 |
| 140 | 0 | 9.999e-05 |
| 145 | 0 | 0.00119988 |
| 150 | 0 | 0.00969903 |
| 155 | 0 | 0.0278972 |
| 160 | 0 | 0.0651935 |
| 165 | 9.999e-05 | 0.107889 |
| 170 | 9.999e-05 | 0.108289 |
| 175 | 0 | 0.0671933 |
| 180 | 0.00109989 | 0.0271973 |
| 185 | 0.0116988 | 0.0156984 |
| 190 | 0.0735926 | 0.0375962 |
| 195 | 0.236376 | 0.0835916 |
| 200 | 0.339766 | 0.130087 |
| 205 | 0.235676 | 0.122188 |
| 210 | 0.0821918 | 0.0839916 |
| 215 | 0.0162984 | 0.0481952 |
| 220 | 0.00239976 | 0.030097 |
| 225 | 0.00069993 | 0.0167983 |
| 230 | 0 | 0.00889911 |
| 235 | 0 | 0.00459954 |
| 240 | 0 | 0.00169983 |
| 245 | 0 | 0.00129987 |
| 250 | 0 | 0.00049995 |
| 255 | 0 | 0 |
| 260 | 0 | 9.999e-05 |
| 265 | 0 | 0 |
| 270 | 0 | 0 |
| 275 | 0 | 0 |
| 280 | 0 | 0 |
| 285 | 0 | 0 |
| 290 | 0 | 0 |
| 295 | 0 | 0 |
| 300 | 0 | 0 |
| 305 | 0 | 0 |
| 310 | 0 | 0 |
| 315 | 0 | 0 |
| 320 | 0 | 0 |
| 325 | 0 | 0 |
| 330 | 0 | 0 |
| 335 | 0 | 0 |
| 340 | 0 | 0 |
| 345 | 0 | 0 |
| 350 | 0 | 0 |
| 355 | 0 | 0 |
| 360 | 0 | 0 |

(j) 283K-apoAB-eta-U48-J2/3

H(η)

| η | NF-apoA | NF-apoB |
| --- | --- | --- |
| 0 | 0.00109989 | 0.00069993 |
| 5 | 0.107989 | 0.0291971 |
| 10 | 0.237276 | 0.0182982 |
| 15 | 0.289871 | 0.0176982 |
| 20 | 0.214579 | 0.0146985 |
| 25 | 0.0842916 | 0.0107989 |
| 30 | 0.0219978 | 0.009999 |
| 35 | 0.00439956 | 0.00849915 |
| 40 | 0.00019998 | 0.0059994 |
| 45 | 0.00019998 | 0.0059994 |
| 50 | 0.00029997 | 0.00629937 |
| 55 | 0.00039996 | 0.00659934 |
| 60 | 9.999e-05 | 0.0140986 |
| 65 | 9.999e-05 | 0.0235976 |
| 70 | 0 | 0.0513949 |
| 75 | 0 | 0.0777922 |
| 80 | 0 | 0.0828917 |
| 85 | 0 | 0.0672933 |
| 90 | 0 | 0.0356964 |
| 95 | 0 | 0.0107989 |
| 100 | 0 | 0.00219978 |
| 105 | 0 | 0.00029997 |
| 110 | 0 | 9.999e-05 |
| 115 | 0 | 0 |
| 120 | 0 | 0 |
| 125 | 0 | 0 |
| 130 | 0 | 0 |
| 135 | 0 | 0 |
| 140 | 0 | 0 |
| 145 | 0 | 0 |
| 150 | 0 | 0 |
| 155 | 0 | 0 |
| 160 | 0 | 0 |
| 165 | 0 | 0 |
| 170 | 0 | 0 |
| 175 | 0 | 0 |
| 180 | 0 | 0 |
| 185 | 0 | 0 |
| 190 | 0 | 0 |
| 195 | 0 | 0 |
| 200 | 0 | 0 |
| 205 | 0 | 0 |
| 210 | 0 | 0 |
| 215 | 0 | 0 |
| 220 | 0 | 0 |
| 225 | 0 | 0 |
| 230 | 0 | 0 |
| 235 | 0 | 0 |
| 240 | 0 | 0 |
| 245 | 0 | 0 |
| 250 | 0 | 0 |
| 255 | 0 | 0 |
| 260 | 0 | 0 |
| 265 | 0 | 0 |
| 270 | 0 | 0 |
| 275 | 0 | 0 |
| 280 | 0 | 0 |
| 285 | 0 | 0 |
| 290 | 0 | 9.999e-05 |
| 295 | 0 | 0.00049995 |
| 300 | 0 | 0.00159984 |
| 305 | 0 | 0.00709929 |
| 310 | 0 | 0.0226977 |
| 315 | 0 | 0.0470953 |
| 320 | 0 | 0.0624938 |
| 325 | 0 | 0.0681932 |
| 330 | 0 | 0.0590941 |
| 335 | 0 | 0.0422958 |
| 340 | 0 | 0.039796 |
| 345 | 0 | 0.0358964 |
| 350 | 0.00069993 | 0.0413959 |
| 355 | 0.00609939 | 0.0372963 |
| 360 | 0.030397 | 0.0334967 |

(k) 283K-apoAB-eta-U49-J2/3

H(η)

| η | NF-apoA | NF-apoB |
| --- | --- | --- |
| 0 | 0 | 0 |
| 5 | 0 | 0 |
| 10 | 0 | 0 |
| 15 | 0 | 0 |
| 20 | 0 | 0 |
| 25 | 0 | 0 |
| 30 | 0 | 0 |
| 35 | 0 | 0 |
| 40 | 0 | 0 |
| 45 | 0 | 0 |
| 50 | 0 | 0 |
| 55 | 0 | 0 |
| 60 | 0 | 0 |
| 65 | 0 | 0 |
| 70 | 0 | 0 |
| 75 | 0 | 0 |
| 80 | 0 | 0 |
| 85 | 0 | 0 |
| 90 | 0 | 0 |
| 95 | 0 | 0 |
| 100 | 0 | 0 |
| 105 | 0 | 0 |
| 110 | 0 | 0 |
| 115 | 0 | 0 |
| 120 | 0 | 0 |
| 125 | 0 | 0 |
| 130 | 0 | 9.999e-05 |
| 135 | 0 | 0.00019998 |
| 140 | 0 | 9.999e-05 |
| 145 | 0 | 0 |
| 150 | 0 | 0.00129987 |
| 155 | 0 | 0.00259974 |
| 160 | 0 | 0.00569943 |
| 165 | 0 | 0.00909909 |
| 170 | 0 | 0.0155984 |
| 175 | 0 | 0.0307969 |
| 180 | 0 | 0.0443956 |
| 185 | 0.00119988 | 0.060194 |
| 190 | 0.0107989 | 0.070193 |
| 195 | 0.0889911 | 0.0868913 |
| 200 | 0.282472 | 0.130487 |
| 205 | 0.376462 | 0.174883 |
| 210 | 0.19558 | 0.181382 |
| 215 | 0.0394961 | 0.113689 |
| 220 | 0.00459954 | 0.0523948 |
| 225 | 0.00029997 | 0.0150985 |
| 230 | 9.999e-05 | 0.00369963 |
| 235 | 0 | 0.00079992 |
| 240 | 0 | 0.00039996 |
| 245 | 0 | 0 |
| 250 | 0 | 0 |
| 255 | 0 | 0 |
| 260 | 0 | 0 |
| 265 | 0 | 0 |
| 270 | 0 | 0 |
| 275 | 0 | 0 |
| 280 | 0 | 0 |
| 285 | 0 | 0 |
| 290 | 0 | 0 |
| 295 | 0 | 0 |
| 300 | 0 | 0 |
| 305 | 0 | 0 |
| 310 | 0 | 0 |
| 315 | 0 | 0 |
| 320 | 0 | 0 |
| 325 | 0 | 0 |
| 330 | 0 | 0 |
| 335 | 0 | 0 |
| 340 | 0 | 0 |
| 345 | 0 | 0 |
| 350 | 0 | 0 |
| 355 | 0 | 0 |
| 360 | 0 | 0 |

(l) 283K-apoAB-eta-U51-J2/3

H(η)

| η | NF-apoA | NF-apoB |
| --- | --- | --- |
| 0 | 0.0029997 | 0.00219978 |
| 5 | 0.169083 | 0.0967903 |
| 10 | 0.150985 | 0.0824918 |
| 15 | 0.114689 | 0.0690931 |
| 20 | 0.0688931 | 0.0778922 |
| 25 | 0.0347965 | 0.0891911 |
| 30 | 0.0140986 | 0.0962904 |
| 35 | 0.00479952 | 0.105289 |
| 40 | 0.00149985 | 0.0935906 |
| 45 | 0 | 0.0524948 |
| 50 | 0 | 0.0131987 |
| 55 | 0 | 0.00369963 |
| 60 | 0 | 0.00019998 |
| 65 | 0 | 9.999e-05 |
| 70 | 0 | 0 |
| 75 | 0 | 0 |
| 80 | 0 | 0 |
| 85 | 0 | 0 |
| 90 | 0 | 0 |
| 95 | 0 | 0 |
| 100 | 0 | 0 |
| 105 | 0 | 0 |
| 110 | 0 | 0 |
| 115 | 0 | 0 |
| 120 | 0 | 0 |
| 125 | 0 | 0 |
| 130 | 0 | 0 |
| 135 | 0 | 0 |
| 140 | 0 | 0 |
| 145 | 0 | 0 |
| 150 | 0 | 0 |
| 155 | 0 | 0 |
| 160 | 0 | 0 |
| 165 | 0 | 0 |
| 170 | 0 | 0 |
| 175 | 0 | 0 |
| 180 | 0 | 0 |
| 185 | 0 | 0 |
| 190 | 0 | 0 |
| 195 | 0 | 0 |
| 200 | 0 | 0 |
| 205 | 0 | 0 |
| 210 | 0 | 0 |
| 215 | 0 | 0 |
| 220 | 0 | 0 |
| 225 | 0 | 0 |
| 230 | 0 | 0 |
| 235 | 0 | 0 |
| 240 | 0 | 0 |
| 245 | 0 | 0 |
| 250 | 0 | 0 |
| 255 | 0 | 0 |
| 260 | 0 | 0 |
| 265 | 0 | 0 |
| 270 | 0 | 0 |
| 275 | 0 | 0 |
| 280 | 0 | 0 |
| 285 | 0 | 9.999e-05 |
| 290 | 9.999e-05 | 0.00019998 |
| 295 | 9.999e-05 | 0.00069993 |
| 300 | 0.00059994 | 0.00089991 |
| 305 | 0.00089991 | 0.00259974 |
| 310 | 0.00139986 | 0.00389961 |
| 315 | 0.00249975 | 0.00289971 |
| 320 | 0.00079992 | 0.00429957 |
| 325 | 0.00179982 | 0.00469953 |
| 330 | 0.00379962 | 0.00409959 |
| 335 | 0.00979902 | 0.00519948 |
| 340 | 0.0184982 | 0.00859914 |
| 345 | 0.0466953 | 0.0153985 |
| 350 | 0.0745925 | 0.0310969 |
| 355 | 0.122288 | 0.0559944 |
| 360 | 0.154285 | 0.0767923 |

(m) 283K-apoAB-eta-A52-J2/3

H(η)

| η | NF-apoA | NF-apoB |
| --- | --- | --- |
| 0 | 0 | 0 |
| 5 | 0 | 0 |
| 10 | 0 | 0 |
| 15 | 0 | 0 |
| 20 | 0 | 0 |
| 25 | 0 | 0 |
| 30 | 0 | 0 |
| 35 | 0 | 0 |
| 40 | 0 | 0 |
| 45 | 0 | 0 |
| 50 | 0 | 0 |
| 55 | 0 | 0 |
| 60 | 0 | 0 |
| 65 | 0 | 0 |
| 70 | 0 | 0 |
| 75 | 0 | 0 |
| 80 | 0 | 0 |
| 85 | 0 | 0 |
| 90 | 0 | 0 |
| 95 | 0 | 0 |
| 100 | 0 | 0 |
| 105 | 0 | 0 |
| 110 | 0 | 0 |
| 115 | 0 | 0 |
| 120 | 0 | 0.00019998 |
| 125 | 0 | 0.00039996 |
| 130 | 0 | 0.00149985 |
| 135 | 0 | 0.00549945 |
| 140 | 0 | 0.0150985 |
| 145 | 0 | 0.0292971 |
| 150 | 0 | 0.0446955 |
| 155 | 0 | 0.0629937 |
| 160 | 0 | 0.090391 |
| 165 | 9.999e-05 | 0.118388 |
| 170 | 0.00079992 | 0.138386 |
| 175 | 0.00669933 | 0.154285 |
| 180 | 0.0285971 | 0.129387 |
| 185 | 0.0790921 | 0.0983902 |
| 190 | 0.157984 | 0.0558944 |
| 195 | 0.20228 | 0.0290971 |
| 200 | 0.20008 | 0.0155984 |
| 205 | 0.154485 | 0.00739926 |
| 210 | 0.0961904 | 0.00179982 |
| 215 | 0.0490951 | 0.00109989 |
| 220 | 0.0184982 | 9.999e-05 |
| 225 | 0.00479952 | 9.999e-05 |
| 230 | 0.00119988 | 0 |
| 235 | 9.999e-05 | 0 |
| 240 | 0 | 0 |
| 245 | 0 | 0 |
| 250 | 0 | 0 |
| 255 | 0 | 0 |
| 260 | 0 | 0 |
| 265 | 0 | 0 |
| 270 | 0 | 0 |
| 275 | 0 | 0 |
| 280 | 0 | 0 |
| 285 | 0 | 0 |
| 290 | 0 | 0 |
| 295 | 0 | 0 |
| 300 | 0 | 0 |
| 305 | 0 | 0 |
| 310 | 0 | 0 |
| 315 | 0 | 0 |
| 320 | 0 | 0 |
| 325 | 0 | 0 |
| 330 | 0 | 0 |
| 335 | 0 | 0 |
| 340 | 0 | 0 |
| 345 | 0 | 0 |
| 350 | 0 | 0 |
| 355 | 0 | 0 |
| 360 | 0 | 0 |

(n) 283K-apoAB-eta-U74-J3/1

H(η)

| η | NF-apoA | NF-apoB |
| --- | --- | --- |
| 0 | 0 | 0 |
| 5 | 0 | 0 |
| 10 | 0 | 0 |
| 15 | 0 | 0 |
| 20 | 0 | 0 |
| 25 | 0 | 0 |
| 30 | 0 | 0 |
| 35 | 0 | 0 |
| 40 | 0 | 0 |
| 45 | 0 | 0 |
| 50 | 0 | 0 |
| 55 | 0 | 0 |
| 60 | 0 | 0 |
| 65 | 0 | 0 |
| 70 | 0 | 0 |
| 75 | 0 | 0 |
| 80 | 0 | 0 |
| 85 | 0 | 0 |
| 90 | 0 | 0 |
| 95 | 0 | 0 |
| 100 | 0 | 0 |
| 105 | 0 | 0 |
| 110 | 0 | 0 |
| 115 | 0 | 0 |
| 120 | 0 | 0.00029997 |
| 125 | 0 | 0.00169983 |
| 130 | 0 | 0.00549945 |
| 135 | 0 | 0.0129987 |
| 140 | 0.00019998 | 0.0354965 |
| 145 | 0.00029997 | 0.0994901 |
| 150 | 0.00509949 | 0.186381 |
| 155 | 0.0277972 | 0.249875 |
| 160 | 0.10159 | 0.214679 |
| 165 | 0.248675 | 0.123688 |
| 170 | 0.29587 | 0.050495 |
| 175 | 0.206779 | 0.0144986 |
| 180 | 0.0851915 | 0.00409959 |
| 185 | 0.0236976 | 0.00059994 |
| 190 | 0.00439956 | 0.00019998 |
| 195 | 0.00039996 | 0 |
| 200 | 0 | 0 |
| 205 | 0 | 0 |
| 210 | 0 | 0 |
| 215 | 0 | 0 |
| 220 | 0 | 0 |
| 225 | 0 | 0 |
| 230 | 0 | 0 |
| 235 | 0 | 0 |
| 240 | 0 | 0 |
| 245 | 0 | 0 |
| 250 | 0 | 0 |
| 255 | 0 | 0 |
| 260 | 0 | 0 |
| 265 | 0 | 0 |
| 270 | 0 | 0 |
| 275 | 0 | 0 |
| 280 | 0 | 0 |
| 285 | 0 | 0 |
| 290 | 0 | 0 |
| 295 | 0 | 0 |
| 300 | 0 | 0 |
| 305 | 0 | 0 |
| 310 | 0 | 0 |
| 315 | 0 | 0 |
| 320 | 0 | 0 |
| 325 | 0 | 0 |
| 330 | 0 | 0 |
| 335 | 0 | 0 |
| 340 | 0 | 0 |
| 345 | 0 | 0 |
| 350 | 0 | 0 |
| 355 | 0 | 0 |
| 360 | 0 | 0 |

**Figure S6: Some representative Histogram distribution of pseudo-torsion angle (θ) at 283K (apoA in blue and apoB in orange))**

(a) 283K-apoAB-theta-G18-Stem1

H(θ)

| θ | NF-apoA | NF-apoB |
| --- | --- | --- |
| 0 | 0 | 0 |
| 5 | 0 | 0 |
| 10 | 0 | 0 |
| 15 | 0 | 0 |
| 20 | 0 | 0 |
| 25 | 0 | 0 |
| 30 | 0 | 0 |
| 35 | 0 | 0 |
| 40 | 0 | 0 |
| 45 | 0 | 0 |
| 50 | 0 | 0 |
| 55 | 0 | 0 |
| 60 | 0 | 0 |
| 65 | 0 | 0 |
| 70 | 0 | 0 |
| 75 | 0 | 0 |
| 80 | 0 | 0 |
| 85 | 0 | 0 |
| 90 | 0 | 0 |
| 95 | 0 | 0 |
| 100 | 0 | 0 |
| 105 | 0 | 0 |
| 110 | 0 | 0 |
| 115 | 0 | 0 |
| 120 | 0 | 0 |
| 125 | 0 | 0 |
| 130 | 0 | 0 |
| 135 | 0 | 0 |
| 140 | 0 | 0 |
| 145 | 0 | 0 |
| 150 | 0 | 0 |
| 155 | 0 | 0 |
| 160 | 0.00069993 | 0 |
| 165 | 0.00109989 | 0 |
| 170 | 0.0029997 | 0 |
| 175 | 0.010099 | 0 |
| 180 | 0.0251975 | 9.999e-05 |
| 185 | 0.0537946 | 0.00079992 |
| 190 | 0.0822918 | 0.00369963 |
| 195 | 0.107489 | 0.0105989 |
| 200 | 0.121088 | 0.0414959 |
| 205 | 0.116488 | 0.10229 |
| 210 | 0.080192 | 0.182782 |
| 215 | 0.0590941 | 0.229577 |
| 220 | 0.0451955 | 0.219878 |
| 225 | 0.0343966 | 0.134287 |
| 230 | 0.0286971 | 0.0563944 |
| 235 | 0.0251975 | 0.0123988 |
| 240 | 0.0192981 | 0.00409959 |
| 245 | 0.0141986 | 0.00139986 |
| 250 | 0.0150985 | 0.00019998 |
| 255 | 0.0138986 | 0 |
| 260 | 0.0264974 | 0 |
| 265 | 0.0378962 | 0 |
| 270 | 0.0361964 | 0 |
| 275 | 0.0228977 | 0 |
| 280 | 0.0110989 | 0 |
| 285 | 0.00589941 | 0 |
| 290 | 0.00229977 | 0 |
| 295 | 0.00069993 | 0 |
| 300 | 0 | 0 |
| 305 | 0 | 0 |
| 310 | 0 | 0 |
| 315 | 0 | 0 |
| 320 | 0 | 0 |
| 325 | 0 | 0 |
| 330 | 0 | 0 |
| 335 | 0 | 0 |
| 340 | 0 | 0 |
| 345 | 0 | 0 |
| 350 | 0 | 0 |
| 355 | 0 | 0 |
| 360 | 0 | 0 |

(b) 283K-apoAB-theta-A19-Stem1

H(θ)

| θ | NF-apoA | NF-apoB |
| --- | --- | --- |
| 0 | 0 | 0 |
| 5 | 0 | 0 |
| 10 | 0 | 0 |
| 15 | 0 | 0 |
| 20 | 0 | 0 |
| 25 | 0 | 0 |
| 30 | 0 | 0 |
| 35 | 0 | 0 |
| 40 | 0 | 0 |
| 45 | 0 | 0 |
| 50 | 0 | 0 |
| 55 | 0 | 0 |
| 60 | 0 | 0 |
| 65 | 0 | 0 |
| 70 | 0 | 0 |
| 75 | 0 | 0 |
| 80 | 0 | 0 |
| 85 | 9.999e-05 | 0 |
| 90 | 0.00019998 | 0 |
| 95 | 0.00049995 | 0 |
| 100 | 0.00159984 | 0 |
| 105 | 0.00179982 | 0 |
| 110 | 0.00429957 | 0 |
| 115 | 0.010099 | 0 |
| 120 | 0.0107989 | 0 |
| 125 | 0.0153985 | 0 |
| 130 | 0.0192981 | 0 |
| 135 | 0.0243976 | 0 |
| 140 | 0.0219978 | 0 |
| 145 | 0.020398 | 0 |
| 150 | 0.0144986 | 0 |
| 155 | 0.0114989 | 0 |
| 160 | 0.00649935 | 0 |
| 165 | 0.00559944 | 0 |
| 170 | 0.00669933 | 0 |
| 175 | 0.00719928 | 0 |
| 180 | 0.00929907 | 0.00029997 |
| 185 | 0.0171983 | 0.00029997 |
| 190 | 0.0288971 | 0.00239976 |
| 195 | 0.0379962 | 0.0108989 |
| 200 | 0.0490951 | 0.0321968 |
| 205 | 0.0579942 | 0.0981902 |
| 210 | 0.0706929 | 0.172083 |
| 215 | 0.090491 | 0.215378 |
| 220 | 0.127487 | 0.20058 |
| 225 | 0.134187 | 0.139386 |
| 230 | 0.10269 | 0.0756924 |
| 235 | 0.0571943 | 0.0344966 |
| 240 | 0.0239976 | 0.0130987 |
| 245 | 0.00759924 | 0.00409959 |
| 250 | 0.00179982 | 0.00049995 |
| 255 | 0.00039996 | 0.00039996 |
| 260 | 9.999e-05 | 0 |
| 265 | 0 | 0 |
| 270 | 0 | 0 |
| 275 | 0 | 0 |
| 280 | 0 | 0 |
| 285 | 0 | 0 |
| 290 | 0 | 0 |
| 295 | 0 | 0 |
| 300 | 0 | 0 |
| 305 | 0 | 0 |
| 310 | 0 | 0 |
| 315 | 0 | 0 |
| 320 | 0 | 0 |
| 325 | 0 | 0 |
| 330 | 0 | 0 |
| 335 | 0 | 0 |
| 340 | 0 | 0 |
| 345 | 0 | 0 |
| 350 | 0 | 0 |
| 355 | 0 | 0 |
| 360 | 0 | 0 |

(c) 283K-apoAB-theta-A21-Stem1

H(θ)

| θ | NF-apoA | NF-apoB |
| --- | --- | --- |
| 0 | 0.00649935 | 0 |
| 5 | 0.274573 | 0 |
| 10 | 0.150685 | 0 |
| 15 | 0.0461954 | 0 |
| 20 | 0.00649935 | 0 |
| 25 | 0.00069993 | 0 |
| 30 | 0 | 0 |
| 35 | 0 | 0 |
| 40 | 0 | 0 |
| 45 | 0 | 0 |
| 50 | 0 | 0 |
| 55 | 0 | 0 |
| 60 | 0 | 0 |
| 65 | 0 | 0 |
| 70 | 0 | 0 |
| 75 | 0 | 0 |
| 80 | 0 | 0 |
| 85 | 0 | 0 |
| 90 | 0 | 0 |
| 95 | 0 | 0 |
| 100 | 0 | 0 |
| 105 | 0 | 0 |
| 110 | 0 | 0 |
| 115 | 0 | 0 |
| 120 | 0 | 0 |
| 125 | 0 | 0 |
| 130 | 0 | 0 |
| 135 | 0 | 0 |
| 140 | 0 | 0 |
| 145 | 0 | 0 |
| 150 | 0 | 0 |
| 155 | 0 | 0 |
| 160 | 0 | 0 |
| 165 | 0 | 0 |
| 170 | 0 | 0 |
| 175 | 0 | 0 |
| 180 | 0 | 0 |
| 185 | 0 | 0 |
| 190 | 0 | 0 |
| 195 | 0 | 0 |
| 200 | 0 | 0 |
| 205 | 0 | 0 |
| 210 | 0 | 0 |
| 215 | 0 | 0 |
| 220 | 0 | 0 |
| 225 | 0 | 0.00039996 |
| 230 | 0 | 0.00019998 |
| 235 | 0 | 9.999e-05 |
| 240 | 0 | 9.999e-05 |
| 245 | 0 | 0.00039996 |
| 250 | 0 | 0.00019998 |
| 255 | 0 | 0.00119988 |
| 260 | 0 | 0.00109989 |
| 265 | 0 | 0.00159984 |
| 270 | 0 | 0.00129987 |
| 275 | 0 | 0.00209979 |
| 280 | 0 | 0.0029997 |
| 285 | 0 | 0.00659934 |
| 290 | 0 | 0.0214979 |
| 295 | 0 | 0.0650935 |
| 300 | 0 | 0.192181 |
| 305 | 0 | 0.342566 |
| 310 | 0 | 0.257774 |
| 315 | 9.999e-05 | 0.0780922 |
| 320 | 0.00049995 | 0.0171983 |
| 325 | 0.00089991 | 0.00549945 |
| 330 | 0.00089991 | 0.00149985 |
| 335 | 0.00069993 | 0.00029997 |
| 340 | 0.00449955 | 0 |
| 345 | 0.0149985 | 0 |
| 350 | 0.0605939 | 0 |
| 355 | 0.162084 | 0 |
| 360 | 0.269573 | 0 |

(d) 283K-apoAB-theta-U22-J1/2

H(θ)

| θ | NF-apoA | NF-apoB |
| --- | --- | --- |
| 0 | 0 | 0 |
| 5 | 0 | 0 |
| 10 | 0 | 0 |
| 15 | 0 | 0 |
| 20 | 0 | 0 |
| 25 | 0 | 0 |
| 30 | 0 | 0 |
| 35 | 0 | 0 |
| 40 | 0 | 0 |
| 45 | 0 | 0 |
| 50 | 0 | 0 |
| 55 | 0 | 0 |
| 60 | 0 | 0 |
| 65 | 0 | 0 |
| 70 | 0 | 0 |
| 75 | 0 | 0 |
| 80 | 0 | 0 |
| 85 | 0 | 0 |
| 90 | 0 | 0 |
| 95 | 0 | 0 |
| 100 | 0.00469953 | 0 |
| 105 | 0.0611939 | 0 |
| 110 | 0.233477 | 0 |
| 115 | 0.380462 | 0 |
| 120 | 0.248175 | 0 |
| 125 | 0.0631937 | 0 |
| 130 | 0.00849915 | 0 |
| 135 | 0.00029997 | 0 |
| 140 | 0 | 0 |
| 145 | 0 | 0 |
| 150 | 0 | 0 |
| 155 | 0 | 0 |
| 160 | 0 | 0 |
| 165 | 0 | 0 |
| 170 | 0 | 0 |
| 175 | 0 | 0 |
| 180 | 0 | 0 |
| 185 | 0 | 0 |
| 190 | 0 | 0 |
| 195 | 0 | 0 |
| 200 | 0 | 0 |
| 205 | 0 | 0 |
| 210 | 0 | 0 |
| 215 | 0 | 0 |
| 220 | 0 | 0 |
| 225 | 0 | 0 |
| 230 | 0 | 0 |
| 235 | 0 | 0 |
| 240 | 0 | 0 |
| 245 | 0 | 0 |
| 250 | 0 | 0 |
| 255 | 0 | 0 |
| 260 | 0 | 0 |
| 265 | 0 | 0 |
| 270 | 0 | 0 |
| 275 | 0 | 0 |
| 280 | 0 | 0.00079992 |
| 285 | 0 | 0.00249975 |
| 290 | 0 | 0.00949905 |
| 295 | 0 | 0.0326967 |
| 300 | 0 | 0.0918908 |
| 305 | 0 | 0.190281 |
| 310 | 0 | 0.261874 |
| 315 | 0 | 0.232177 |
| 320 | 0 | 0.125887 |
| 325 | 0 | 0.0424958 |
| 330 | 0 | 0.00849915 |
| 335 | 0 | 0.00119988 |
| 340 | 0 | 9.999e-05 |
| 345 | 0 | 9.999e-05 |
| 350 | 0 | 0 |
| 355 | 0 | 0 |
| 360 | 0 | 0 |

(e) 283K-apoAB-theta-A23-J1/2

H(θ)

| θ | NF-apoA | NF-apoB |
| --- | --- | --- |
| 0 | 0.00449955 | 0 |
| 5 | 0.184782 | 0 |
| 10 | 0.110589 | 0 |
| 15 | 0.0533947 | 0 |
| 20 | 0.0214979 | 0 |
| 25 | 0.00639936 | 0 |
| 30 | 0.00089991 | 0 |
| 35 | 0.00039996 | 0 |
| 40 | 0 | 0 |
| 45 | 9.999e-05 | 0 |
| 50 | 0 | 0 |
| 55 | 0 | 0 |
| 60 | 0 | 0 |
| 65 | 0 | 0 |
| 70 | 0 | 0 |
| 75 | 0 | 0 |
| 80 | 0 | 0 |
| 85 | 0 | 0 |
| 90 | 0 | 0 |
| 95 | 0 | 0 |
| 100 | 0 | 0 |
| 105 | 0 | 0 |
| 110 | 0 | 0 |
| 115 | 0 | 0 |
| 120 | 0 | 0 |
| 125 | 0 | 0 |
| 130 | 0 | 0 |
| 135 | 0 | 0.00049995 |
| 140 | 0 | 0.00769923 |
| 145 | 0 | 0.0407959 |
| 150 | 0 | 0.123188 |
| 155 | 0 | 0.268773 |
| 160 | 0 | 0.285971 |
| 165 | 0 | 0.180582 |
| 170 | 0 | 0.0636936 |
| 175 | 0 | 0.0150985 |
| 180 | 0 | 0.00529947 |
| 185 | 0 | 0.00169983 |
| 190 | 0 | 0.00249975 |
| 195 | 0 | 0.00329967 |
| 200 | 0 | 0.00069993 |
| 205 | 0 | 9.999e-05 |
| 210 | 0 | 9.999e-05 |
| 215 | 0 | 0 |
| 220 | 0 | 0 |
| 225 | 0 | 0 |
| 230 | 0 | 0 |
| 235 | 0 | 0 |
| 240 | 0 | 0 |
| 245 | 0 | 0 |
| 250 | 0 | 0 |
| 255 | 0 | 0 |
| 260 | 0 | 0 |
| 265 | 0 | 0 |
| 270 | 0 | 0 |
| 275 | 0 | 0 |
| 280 | 0 | 0 |
| 285 | 0 | 0 |
| 290 | 0 | 0 |
| 295 | 0 | 0 |
| 300 | 0 | 0 |
| 305 | 0 | 0 |
| 310 | 0 | 0 |
| 315 | 0 | 0 |
| 320 | 0 | 0 |
| 325 | 0 | 0 |
| 330 | 0.00019998 | 0 |
| 335 | 0.00219978 | 0 |
| 340 | 0.0149985 | 0 |
| 345 | 0.0505949 | 0 |
| 350 | 0.128087 | 0 |
| 355 | 0.20288 | 0 |
| 360 | 0.218478 | 0 |

(f) 283K-apoAB-theta-A24-J1/2

H(θ)

| θ | NF-apoA | NF-apoB |
| --- | --- | --- |
| 0 | 0 | 0 |
| 5 | 0 | 0 |
| 10 | 0 | 0 |
| 15 | 0 | 0 |
| 20 | 0 | 0 |
| 25 | 0 | 0 |
| 30 | 0 | 0 |
| 35 | 0.00029997 | 0 |
| 40 | 0.00879912 | 0 |
| 45 | 0.0712929 | 0.00029997 |
| 50 | 0.262374 | 0.00119988 |
| 55 | 0.412459 | 0.00279972 |
| 60 | 0.207179 | 0.00589941 |
| 65 | 0.0336966 | 0.00779922 |
| 70 | 0.00349965 | 0.00729927 |
| 75 | 0.00039996 | 0.0105989 |
| 80 | 0 | 0.029997 |
| 85 | 0 | 0.105889 |
| 90 | 0 | 0.247175 |
| 95 | 0 | 0.322568 |
| 100 | 0 | 0.19748 |
| 105 | 0 | 0.0531947 |
| 110 | 0 | 0.00709929 |
| 115 | 0 | 0.00059994 |
| 120 | 0 | 9.999e-05 |
| 125 | 0 | 0 |
| 130 | 0 | 0 |
| 135 | 0 | 0 |
| 140 | 0 | 0 |
| 145 | 0 | 0 |
| 150 | 0 | 0 |
| 155 | 0 | 0 |
| 160 | 0 | 0 |
| 165 | 0 | 0 |
| 170 | 0 | 0 |
| 175 | 0 | 0 |
| 180 | 0 | 0 |
| 185 | 0 | 0 |
| 190 | 0 | 0 |
| 195 | 0 | 0 |
| 200 | 0 | 0 |
| 205 | 0 | 0 |
| 210 | 0 | 0 |
| 215 | 0 | 0 |
| 220 | 0 | 0 |
| 225 | 0 | 0 |
| 230 | 0 | 0 |
| 235 | 0 | 0 |
| 240 | 0 | 0 |
| 245 | 0 | 0 |
| 250 | 0 | 0 |
| 255 | 0 | 0 |
| 260 | 0 | 0 |
| 265 | 0 | 0 |
| 270 | 0 | 0 |
| 275 | 0 | 0 |
| 280 | 0 | 0 |
| 285 | 0 | 0 |
| 290 | 0 | 0 |
| 295 | 0 | 0 |
| 300 | 0 | 0 |
| 305 | 0 | 0 |
| 310 | 0 | 0 |
| 315 | 0 | 0 |
| 320 | 0 | 0 |
| 325 | 0 | 0 |
| 330 | 0 | 0 |
| 335 | 0 | 0 |
| 340 | 0 | 0 |
| 345 | 0 | 0 |
| 350 | 0 | 0 |
| 355 | 0 | 0 |
| 360 | 0 | 0 |

(g) 283K-apoAB-theta-A35-Loop2

H(θ)

| θ | NF-apoA | NF-apoB |
| --- | --- | --- |
| 0 | 0 | 0 |
| 5 | 0 | 0 |
| 10 | 0 | 0 |
| 15 | 0 | 0 |
| 20 | 0 | 0 |
| 25 | 0 | 0 |
| 30 | 0 | 0 |
| 35 | 0 | 0 |
| 40 | 0 | 0 |
| 45 | 0 | 0 |
| 50 | 0 | 0 |
| 55 | 0 | 0 |
| 60 | 0 | 0 |
| 65 | 0 | 0 |
| 70 | 0 | 9.999e-05 |
| 75 | 0 | 0.00079992 |
| 80 | 0 | 0.00089991 |
| 85 | 0 | 0.00509949 |
| 90 | 9.999e-05 | 0.0250975 |
| 95 | 0.0009999 | 0.0776922 |
| 100 | 0.00279972 | 0.139686 |
| 105 | 0.00539946 | 0.171783 |
| 110 | 0.0122988 | 0.170083 |
| 115 | 0.0205979 | 0.184482 |
| 120 | 0.0334967 | 0.144886 |
| 125 | 0.059594 | 0.060294 |
| 130 | 0.105689 | 0.0147985 |
| 135 | 0.146885 | 0.00339966 |
| 140 | 0.167183 | 0.00069993 |
| 145 | 0.177882 | 0.00019998 |
| 150 | 0.141586 | 0 |
| 155 | 0.0789921 | 0 |
| 160 | 0.0342966 | 0 |
| 165 | 0.00969903 | 0 |
| 170 | 0.00219978 | 0 |
| 175 | 0.00029997 | 0 |
| 180 | 0 | 0 |
| 185 | 0 | 0 |
| 190 | 0 | 0 |
| 195 | 0 | 0 |
| 200 | 0 | 0 |
| 205 | 0 | 0 |
| 210 | 0 | 0 |
| 215 | 0 | 0 |
| 220 | 0 | 0 |
| 225 | 0 | 0 |
| 230 | 0 | 0 |
| 235 | 0 | 0 |
| 240 | 0 | 0 |
| 245 | 0 | 0 |
| 250 | 0 | 0 |
| 255 | 0 | 0 |
| 260 | 0 | 0 |
| 265 | 0 | 0 |
| 270 | 0 | 0 |
| 275 | 0 | 0 |
| 280 | 0 | 0 |
| 285 | 0 | 0 |
| 290 | 0 | 0 |
| 295 | 0 | 0 |
| 300 | 0 | 0 |
| 305 | 0 | 0 |
| 310 | 0 | 0 |
| 315 | 0 | 0 |
| 320 | 0 | 0 |
| 325 | 0 | 0 |
| 330 | 0 | 0 |
| 335 | 0 | 0 |
| 340 | 0 | 0 |
| 345 | 0 | 0 |
| 350 | 0 | 0 |
| 355 | 0 | 0 |
| 360 | 0 | 0 |

(h) 283K-apoAB-theta-A29-Stem2

H(θ)

| θ | NF-apoA | NF-apoB |
| --- | --- | --- |
| 0 | 0 | 0 |
| 5 | 0 | 0 |
| 10 | 0 | 0 |
| 15 | 0 | 0 |
| 20 | 0 | 0 |
| 25 | 0 | 0 |
| 30 | 0 | 0 |
| 35 | 0 | 0 |
| 40 | 0 | 0 |
| 45 | 0 | 0 |
| 50 | 0 | 0 |
| 55 | 0 | 0 |
| 60 | 0 | 0 |
| 65 | 0 | 0 |
| 70 | 0 | 0 |
| 75 | 0 | 0 |
| 80 | 0 | 0 |
| 85 | 0 | 0 |
| 90 | 0 | 0 |
| 95 | 0 | 0 |
| 100 | 0 | 0 |
| 105 | 0 | 0 |
| 110 | 0 | 0 |
| 115 | 0 | 0 |
| 120 | 0 | 0 |
| 125 | 0 | 0 |
| 130 | 0 | 0 |
| 135 | 0 | 0 |
| 140 | 0 | 0 |
| 145 | 0 | 0 |
| 150 | 0 | 0 |
| 155 | 0 | 0 |
| 160 | 0 | 0 |
| 165 | 0 | 0 |
| 170 | 0 | 0 |
| 175 | 0 | 0 |
| 180 | 0 | 0 |
| 185 | 0 | 0.00039996 |
| 190 | 0.00239976 | 0.00179982 |
| 195 | 0.0167983 | 0.0115988 |
| 200 | 0.0768923 | 0.0428957 |
| 205 | 0.193381 | 0.106289 |
| 210 | 0.253975 | 0.149185 |
| 215 | 0.223978 | 0.164584 |
| 220 | 0.142986 | 0.154185 |
| 225 | 0.0610939 | 0.148485 |
| 230 | 0.0193981 | 0.111489 |
| 235 | 0.00539946 | 0.0585941 |
| 240 | 0.00279972 | 0.0278972 |
| 245 | 0.00069993 | 0.0137986 |
| 250 | 0.00019998 | 0.00619938 |
| 255 | 0 | 0.00209979 |
| 260 | 0 | 0.00049995 |
| 265 | 0 | 0 |
| 270 | 0 | 0 |
| 275 | 0 | 0 |
| 280 | 0 | 0 |
| 285 | 0 | 0 |
| 290 | 0 | 0 |
| 295 | 0 | 0 |
| 300 | 0 | 0 |
| 305 | 0 | 0 |
| 310 | 0 | 0 |
| 315 | 0 | 0 |
| 320 | 0 | 0 |
| 325 | 0 | 0 |
| 330 | 0 | 0 |
| 335 | 0 | 0 |
| 340 | 0 | 0 |
| 345 | 0 | 0 |
| 350 | 0 | 0 |
| 355 | 0 | 0 |
| 360 | 0 | 0 |

(i) 283K-apoAB-theta-U47-J2/3

H(θ)

| θ | NF-apoA | NF-apoB |
| --- | --- | --- |
| 0 | 0 | 0 |
| 5 | 0 | 0 |
| 10 | 0 | 0 |
| 15 | 0 | 0 |
| 20 | 0 | 0 |
| 25 | 0 | 0 |
| 30 | 0 | 0 |
| 35 | 0 | 0 |
| 40 | 0 | 0 |
| 45 | 0 | 0 |
| 50 | 0 | 0 |
| 55 | 0 | 0 |
| 60 | 0 | 0 |
| 65 | 0 | 0 |
| 70 | 0 | 0 |
| 75 | 0 | 0 |
| 80 | 0 | 0 |
| 85 | 0 | 0 |
| 90 | 0 | 0 |
| 95 | 0 | 0 |
| 100 | 0 | 0 |
| 105 | 0 | 0 |
| 110 | 0 | 0 |
| 115 | 0 | 0 |
| 120 | 0 | 0 |
| 125 | 0 | 0 |
| 130 | 0 | 0 |
| 135 | 0 | 0 |
| 140 | 0 | 0 |
| 145 | 0 | 0 |
| 150 | 0 | 0 |
| 155 | 0 | 0.00019998 |
| 160 | 0 | 0.00169983 |
| 165 | 0 | 0.0069993 |
| 170 | 0 | 0.0216978 |
| 175 | 0 | 0.0392961 |
| 180 | 0 | 0.0612939 |
| 185 | 0 | 0.0680932 |
| 190 | 0 | 0.0544946 |
| 195 | 9.999e-05 | 0.0354965 |
| 200 | 9.999e-05 | 0.0323968 |
| 205 | 0.00079992 | 0.0210979 |
| 210 | 0.00119988 | 0.0177982 |
| 215 | 0.00629937 | 0.0141986 |
| 220 | 0.0188981 | 0.0143986 |
| 225 | 0.059894 | 0.0151985 |
| 230 | 0.122788 | 0.0183982 |
| 235 | 0.184682 | 0.0266973 |
| 240 | 0.214679 | 0.0293971 |
| 245 | 0.182782 | 0.0348965 |
| 250 | 0.118288 | 0.0372963 |
| 255 | 0.060194 | 0.0417958 |
| 260 | 0.0217978 | 0.0427957 |
| 265 | 0.00549945 | 0.0394961 |
| 270 | 0.00149985 | 0.0380962 |
| 275 | 0.00049995 | 0.0351965 |
| 280 | 0 | 0.0309969 |
| 285 | 0 | 0.0315968 |
| 290 | 0 | 0.0286971 |
| 295 | 0 | 0.029597 |
| 300 | 0 | 0.0324968 |
| 305 | 0 | 0.0315968 |
| 310 | 0 | 0.0280972 |
| 315 | 0 | 0.0170983 |
| 320 | 0 | 0.0121988 |
| 325 | 0 | 0.00639936 |
| 330 | 0 | 0.00219978 |
| 335 | 0 | 0.00049995 |
| 340 | 0 | 9.999e-05 |
| 345 | 0 | 0 |
| 350 | 0 | 0 |
| 355 | 0 | 0 |
| 360 | 0 | 0 |

(j) 283K-apoAB-theta-U48-J2/3

H(θ)

| θ | NF-apoA | NF-apoB |
| --- | --- | --- |
| 0 | 0 | 0 |
| 5 | 0 | 0 |
| 10 | 0 | 0 |
| 15 | 0 | 0 |
| 20 | 0 | 0 |
| 25 | 0 | 0 |
| 30 | 0 | 0 |
| 35 | 0 | 0 |
| 40 | 0 | 0 |
| 45 | 0 | 0 |
| 50 | 0 | 0 |
| 55 | 0 | 0 |
| 60 | 0 | 0 |
| 65 | 0 | 0 |
| 70 | 0 | 0 |
| 75 | 0 | 0 |
| 80 | 0 | 0 |
| 85 | 0 | 0 |
| 90 | 0 | 0 |
| 95 | 0 | 0 |
| 100 | 0 | 0 |
| 105 | 0 | 0 |
| 110 | 0 | 0 |
| 115 | 0 | 0 |
| 120 | 0 | 0 |
| 125 | 0 | 0 |
| 130 | 0 | 9.999e-05 |
| 135 | 0 | 0 |
| 140 | 0 | 0.00029997 |
| 145 | 0 | 0 |
| 150 | 0 | 0.00029997 |
| 155 | 0 | 0.00149985 |
| 160 | 0 | 0.00619938 |
| 165 | 0 | 0.020198 |
| 170 | 0 | 0.0407959 |
| 175 | 0 | 0.0578942 |
| 180 | 0 | 0.0549945 |
| 185 | 9.999e-05 | 0.0442956 |
| 190 | 9.999e-05 | 0.0307969 |
| 195 | 0.0029997 | 0.0359964 |
| 200 | 0.0359964 | 0.0382962 |
| 205 | 0.20208 | 0.050095 |
| 210 | 0.417658 | 0.0536946 |
| 215 | 0.270073 | 0.0489951 |
| 220 | 0.0633937 | 0.0544946 |
| 225 | 0.00729927 | 0.0547945 |
| 230 | 0.00019998 | 0.0530947 |
| 235 | 0 | 0.0463954 |
| 240 | 9.999e-05 | 0.0382962 |
| 245 | 0 | 0.029597 |
| 250 | 0 | 0.0266973 |
| 255 | 0 | 0.0347965 |
| 260 | 0 | 0.0453955 |
| 265 | 0 | 0.0452955 |
| 270 | 0 | 0.0361964 |
| 275 | 0 | 0.0226977 |
| 280 | 0 | 0.0138986 |
| 285 | 0 | 0.00819918 |
| 290 | 0 | 0.00369963 |
| 295 | 0 | 0.00169983 |
| 300 | 0 | 0.00019998 |
| 305 | 0 | 9.999e-05 |
| 310 | 0 | 0 |
| 315 | 0 | 0 |
| 320 | 0 | 0 |
| 325 | 0 | 0 |
| 330 | 0 | 0 |
| 335 | 0 | 0 |
| 340 | 0 | 0 |
| 345 | 0 | 0 |
| 350 | 0 | 0 |
| 355 | 0 | 0 |
| 360 | 0 | 0 |

(k) 283K-apoAB-theta-A52-J2/3

H(θ)

| θ | NF-apoA | NF-apoB |
| --- | --- | --- |
| 0 | 0 | 0 |
| 5 | 0 | 0 |
| 10 | 0 | 0 |
| 15 | 0 | 0 |
| 20 | 0 | 0 |
| 25 | 0 | 0 |
| 30 | 0 | 0 |
| 35 | 0 | 0 |
| 40 | 0 | 0 |
| 45 | 0 | 0 |
| 50 | 0 | 0 |
| 55 | 0 | 0 |
| 60 | 0 | 0 |
| 65 | 0 | 0 |
| 70 | 0 | 0 |
| 75 | 0 | 0 |
| 80 | 0 | 0 |
| 85 | 0 | 0 |
| 90 | 0 | 0 |
| 95 | 0 | 0 |
| 100 | 0 | 0 |
| 105 | 0 | 0 |
| 110 | 0 | 0 |
| 115 | 0 | 0 |
| 120 | 0 | 0 |
| 125 | 0 | 0 |
| 130 | 0 | 0 |
| 135 | 0 | 0 |
| 140 | 0 | 0 |
| 145 | 0 | 0 |
| 150 | 0 | 9.999e-05 |
| 155 | 0 | 9.999e-05 |
| 160 | 0 | 0.00109989 |
| 165 | 0 | 0.00279972 |
| 170 | 0 | 0.00929907 |
| 175 | 0 | 0.0179982 |
| 180 | 0 | 0.0321968 |
| 185 | 0 | 0.0493951 |
| 190 | 0 | 0.0674933 |
| 195 | 0 | 0.0859914 |
| 200 | 9.999e-05 | 0.110089 |
| 205 | 0.00129987 | 0.132887 |
| 210 | 0.00979902 | 0.137086 |
| 215 | 0.0487951 | 0.124288 |
| 220 | 0.149685 | 0.0926907 |
| 225 | 0.260274 | 0.0653935 |
| 230 | 0.240976 | 0.0364964 |
| 235 | 0.157584 | 0.0188981 |
| 240 | 0.0825917 | 0.010199 |
| 245 | 0.0340966 | 0.00319968 |
| 250 | 0.010199 | 0.00119988 |
| 255 | 0.00249975 | 0.00079992 |
| 260 | 0.00079992 | 0.00019998 |
| 265 | 0.00129987 | 9.999e-05 |
| 270 | 0 | 0 |
| 275 | 0 | 0 |
| 280 | 0 | 0 |
| 285 | 0 | 0 |
| 290 | 0 | 0 |
| 295 | 0 | 0 |
| 300 | 0 | 0 |
| 305 | 0 | 0 |
| 310 | 0 | 0 |
| 315 | 0 | 0 |
| 320 | 0 | 0 |
| 325 | 0 | 0 |
| 330 | 0 | 0 |
| 335 | 0 | 0 |
| 340 | 0 | 0 |
| 345 | 0 | 0 |
| 350 | 0 | 0 |
| 355 | 0 | 0 |
| 360 | 0 | 0 |

(l) 283K-apoAB-theta-A73-J3/1

H(θ)

| θ | NF-apoA | NF-apoB |
| --- | --- | --- |
| 0 | 0 | 0 |
| 5 | 0 | 0 |
| 10 | 0 | 0 |
| 15 | 0 | 0 |
| 20 | 0 | 0 |
| 25 | 0 | 0 |
| 30 | 0 | 0 |
| 35 | 0 | 0 |
| 40 | 0 | 0 |
| 45 | 0 | 0 |
| 50 | 0 | 0 |
| 55 | 0 | 0 |
| 60 | 0 | 0 |
| 65 | 0 | 0 |
| 70 | 0 | 0 |
| 75 | 0 | 0 |
| 80 | 0 | 0 |
| 85 | 0 | 0 |
| 90 | 0 | 0 |
| 95 | 0 | 0 |
| 100 | 0 | 0 |
| 105 | 0 | 0 |
| 110 | 0 | 0 |
| 115 | 0 | 0 |
| 120 | 0 | 0 |
| 125 | 0 | 0 |
| 130 | 0 | 0 |
| 135 | 0 | 0 |
| 140 | 0 | 0 |
| 145 | 0 | 0 |
| 150 | 0 | 0 |
| 155 | 0 | 0 |
| 160 | 0 | 0 |
| 165 | 0 | 0 |
| 170 | 9.999e-05 | 0 |
| 175 | 0.00029997 | 0 |
| 180 | 0.00269973 | 0 |
| 185 | 0.0144986 | 0 |
| 190 | 0.030297 | 0 |
| 195 | 0.0493951 | 0.00059994 |
| 200 | 0.0530947 | 0.00039996 |
| 205 | 0.0744926 | 0.0009999 |
| 210 | 0.116888 | 0.0019998 |
| 215 | 0.188581 | 0.00679932 |
| 220 | 0.229077 | 0.0208979 |
| 225 | 0.160884 | 0.0483952 |
| 230 | 0.0618938 | 0.10049 |
| 235 | 0.0153985 | 0.144186 |
| 240 | 0.00219978 | 0.173283 |
| 245 | 0.00019998 | 0.189281 |
| 250 | 0 | 0.163484 |
| 255 | 0 | 0.10039 |
| 260 | 0 | 0.0368963 |
| 265 | 0 | 0.0106989 |
| 270 | 0 | 0.0009999 |
| 275 | 0 | 0.00019998 |
| 280 | 0 | 0 |
| 285 | 0 | 0 |
| 290 | 0 | 0 |
| 295 | 0 | 0 |
| 300 | 0 | 0 |
| 305 | 0 | 0 |
| 310 | 0 | 0 |
| 315 | 0 | 0 |
| 320 | 0 | 0 |
| 325 | 0 | 0 |
| 330 | 0 | 0 |
| 335 | 0 | 0 |
| 340 | 0 | 0 |
| 345 | 0 | 0 |
| 350 | 0 | 0 |
| 355 | 0 | 0 |
| 360 | 0 | 0 |

**Figure S7: Some representative Histogram distribution of pseudo-torsion angle (η) at 293K (apoA in blue and apoB in orange))**

(a) 293K-apoAB-eta-U22-J1/2

H(η)

| η | NF-apoA | NF-apoB |
| --- | --- | --- |
| 0 | 0 | 0 |
| 5 | 0 | 0 |
| 10 | 0 | 0 |
| 15 | 0 | 0 |
| 20 | 0 | 0 |
| 25 | 0 | 0 |
| 30 | 0 | 0 |
| 35 | 0 | 0 |
| 40 | 0 | 0 |
| 45 | 0 | 0 |
| 50 | 0 | 0 |
| 55 | 0 | 0 |
| 60 | 0 | 0 |
| 65 | 0 | 0 |
| 70 | 0 | 0 |
| 75 | 0 | 0 |
| 80 | 0 | 0 |
| 85 | 0 | 0 |
| 90 | 0 | 0 |
| 95 | 0 | 0 |
| 100 | 0 | 0 |
| 105 | 0 | 0 |
| 110 | 0 | 0 |
| 115 | 0 | 0 |
| 120 | 0 | 0 |
| 125 | 0 | 0 |
| 130 | 0 | 0 |
| 135 | 0 | 0 |
| 140 | 0 | 0 |
| 145 | 0 | 0 |
| 150 | 0 | 0 |
| 155 | 0 | 0 |
| 160 | 0 | 0 |
| 165 | 0 | 0 |
| 170 | 0 | 0 |
| 175 | 0 | 0 |
| 180 | 0 | 0 |
| 185 | 0 | 0 |
| 190 | 0 | 0 |
| 195 | 0 | 0 |
| 200 | 0 | 0 |
| 205 | 0 | 0 |
| 210 | 0.00259974 | 0 |
| 215 | 0.0354965 | 0 |
| 220 | 0.155984 | 0 |
| 225 | 0.30007 | 0 |
| 230 | 0.271173 | 0.00029997 |
| 235 | 0.160484 | 0.00139986 |
| 240 | 0.060294 | 0.00409959 |
| 245 | 0.0119988 | 0.0168983 |
| 250 | 0.00189981 | 0.0551945 |
| 255 | 0 | 0.111289 |
| 260 | 0 | 0.142386 |
| 265 | 0 | 0.142686 |
| 270 | 0 | 0.138386 |
| 275 | 0 | 0.146585 |
| 280 | 0 | 0.123888 |
| 285 | 0 | 0.070493 |
| 290 | 0 | 0.029897 |
| 295 | 0 | 0.00929907 |
| 300 | 0 | 0.00329967 |
| 305 | 0 | 0.00109989 |
| 310 | 0 | 0.00059994 |
| 315 | 0 | 0.00049995 |
| 320 | 0 | 0.00049995 |
| 325 | 0 | 0.00089991 |
| 330 | 0 | 0.00029997 |
| 335 | 0 | 0 |
| 340 | 0 | 0 |
| 345 | 0 | 0 |
| 350 | 0 | 0 |
| 355 | 0 | 0 |
| 360 | 0 | 0 |

(b) 293K-apoAB-eta-U25-Stem2

H(η)

| η | NF-apoA | NF-apoB |
| --- | --- | --- |
| 0 | 0 | 0 |
| 5 | 0 | 0 |
| 10 | 0 | 0 |
| 15 | 0 | 0 |
| 20 | 0 | 0 |
| 25 | 0 | 0 |
| 30 | 0 | 0 |
| 35 | 0 | 0 |
| 40 | 0 | 0 |
| 45 | 0 | 0 |
| 50 | 0 | 0 |
| 55 | 0 | 0 |
| 60 | 0 | 0 |
| 65 | 0 | 0 |
| 70 | 0 | 0 |
| 75 | 0 | 0 |
| 80 | 0 | 0 |
| 85 | 0 | 0 |
| 90 | 0 | 0 |
| 95 | 0 | 0 |
| 100 | 0 | 0 |
| 105 | 0 | 0 |
| 110 | 0 | 0 |
| 115 | 0 | 0 |
| 120 | 0 | 0 |
| 125 | 0 | 0 |
| 130 | 0 | 0 |
| 135 | 0 | 0 |
| 140 | 0 | 0 |
| 145 | 0.00049995 | 0 |
| 150 | 0.00129987 | 0 |
| 155 | 0.0132987 | 0.00049995 |
| 160 | 0.0706929 | 0.00529947 |
| 165 | 0.184082 | 0.0179982 |
| 170 | 0.290971 | 0.0534947 |
| 175 | 0.240076 | 0.113489 |
| 180 | 0.118588 | 0.170783 |
| 185 | 0.0392961 | 0.164084 |
| 190 | 0.0193981 | 0.143986 |
| 195 | 0.0133987 | 0.10229 |
| 200 | 0.00649935 | 0.0771923 |
| 205 | 0.00159984 | 0.0609939 |
| 210 | 0.00029997 | 0.0481952 |
| 215 | 0 | 0.0278972 |
| 220 | 0 | 0.010399 |
| 225 | 0 | 0.00239976 |
| 230 | 0 | 0.00089991 |
| 235 | 0 | 9.999e-05 |
| 240 | 0 | 0 |
| 245 | 0 | 0 |
| 250 | 0 | 0 |
| 255 | 0 | 0 |
| 260 | 0 | 0 |
| 265 | 0 | 0 |
| 270 | 0 | 0 |
| 275 | 0 | 0 |
| 280 | 0 | 0 |
| 285 | 0 | 0 |
| 290 | 0 | 0 |
| 295 | 0 | 0 |
| 300 | 0 | 0 |
| 305 | 0 | 0 |
| 310 | 0 | 0 |
| 315 | 0 | 0 |
| 320 | 0 | 0 |
| 325 | 0 | 0 |
| 330 | 0 | 0 |
| 335 | 0 | 0 |
| 340 | 0 | 0 |
| 345 | 0 | 0 |
| 350 | 0 | 0 |
| 355 | 0 | 0 |
| 360 | 0 | 0 |

(c) 293K-apoAB-eta-U36-Loop2

H(η)

| η | NF-apoA | NF-apoB |
| --- | --- | --- |
| 0 | 0.00309969 | 0.00149985 |
| 5 | 0.19918 | 0.0755924 |
| 10 | 0.176182 | 0.0721928 |
| 15 | 0.128087 | 0.0749925 |
| 20 | 0.0749925 | 0.0928907 |
| 25 | 0.0376962 | 0.115988 |
| 30 | 0.0160984 | 0.127687 |
| 35 | 0.00709929 | 0.119888 |
| 40 | 0.00289971 | 0.0891911 |
| 45 | 0.0009999 | 0.0367963 |
| 50 | 0.00019998 | 0.0115988 |
| 55 | 9.999e-05 | 0.00439956 |
| 60 | 0 | 0.00179982 |
| 65 | 0 | 0.00079992 |
| 70 | 0 | 0.00059994 |
| 75 | 0 | 0 |
| 80 | 0 | 0 |
| 85 | 0 | 0 |
| 90 | 0 | 0 |
| 95 | 0 | 0 |
| 100 | 0 | 0 |
| 105 | 0 | 0 |
| 110 | 0 | 0 |
| 115 | 0 | 0 |
| 120 | 0 | 0 |
| 125 | 0 | 0 |
| 130 | 0 | 0 |
| 135 | 0 | 0 |
| 140 | 0 | 0 |
| 145 | 0 | 0 |
| 150 | 0 | 0 |
| 155 | 0 | 0 |
| 160 | 0 | 0 |
| 165 | 0 | 0 |
| 170 | 0 | 0 |
| 175 | 0 | 0 |
| 180 | 0 | 0 |
| 185 | 0 | 0 |
| 190 | 0 | 0 |
| 195 | 0 | 0 |
| 200 | 0 | 0 |
| 205 | 0 | 0 |
| 210 | 0 | 0 |
| 215 | 0 | 0 |
| 220 | 0 | 0 |
| 225 | 0 | 0 |
| 230 | 0 | 0 |
| 235 | 0 | 0 |
| 240 | 0 | 0 |
| 245 | 0 | 0 |
| 250 | 0 | 0 |
| 255 | 0 | 0 |
| 260 | 0 | 0 |
| 265 | 0 | 0 |
| 270 | 0 | 0 |
| 275 | 0 | 0 |
| 280 | 0 | 0 |
| 285 | 0 | 0 |
| 290 | 0 | 0 |
| 295 | 0 | 0 |
| 300 | 0 | 0 |
| 305 | 0 | 0 |
| 310 | 0 | 0 |
| 315 | 0 | 0 |
| 320 | 0 | 0 |
| 325 | 9.999e-05 | 0.00029997 |
| 330 | 9.999e-05 | 0.00089991 |
| 335 | 0.0009999 | 0.00259974 |
| 340 | 0.00609939 | 0.00479952 |
| 345 | 0.0182982 | 0.0153985 |
| 350 | 0.050495 | 0.030197 |
| 355 | 0.108789 | 0.0516948 |
| 360 | 0.168483 | 0.0681932 |

(d) 293K-apoAB-eta-G37-Loop2

H(η)

| η | NF-apoA | NF-apoB |
| --- | --- | --- |
| 0 | 0 | 0 |
| 5 | 0 | 0 |
| 10 | 0 | 0 |
| 15 | 0 | 0 |
| 20 | 0 | 0 |
| 25 | 0 | 0 |
| 30 | 0 | 0 |
| 35 | 0 | 0 |
| 40 | 0 | 0 |
| 45 | 0 | 0 |
| 50 | 0 | 0 |
| 55 | 0 | 0 |
| 60 | 0 | 0 |
| 65 | 0 | 0 |
| 70 | 0 | 0 |
| 75 | 0 | 0 |
| 80 | 0 | 0 |
| 85 | 0 | 0 |
| 90 | 0 | 0 |
| 95 | 0 | 0 |
| 100 | 0 | 0 |
| 105 | 0 | 0 |
| 110 | 0 | 0 |
| 115 | 0 | 9.999e-05 |
| 120 | 0 | 9.999e-05 |
| 125 | 0 | 0.00029997 |
| 130 | 0 | 0.00079992 |
| 135 | 0 | 0.00319968 |
| 140 | 0 | 0.00849915 |
| 145 | 0 | 0.0229977 |
| 150 | 0 | 0.0440956 |
| 155 | 0 | 0.0806919 |
| 160 | 0 | 0.110789 |
| 165 | 9.999e-05 | 0.150685 |
| 170 | 0.00129987 | 0.153485 |
| 175 | 0.00889911 | 0.142086 |
| 180 | 0.0329967 | 0.115888 |
| 185 | 0.0820918 | 0.0839916 |
| 190 | 0.136386 | 0.0489951 |
| 195 | 0.193081 | 0.0243976 |
| 200 | 0.210079 | 0.00789921 |
| 205 | 0.170583 | 0.00089991 |
| 210 | 0.0979902 | 9.999e-05 |
| 215 | 0.0464954 | 0 |
| 220 | 0.0153985 | 0 |
| 225 | 0.00369963 | 0 |
| 230 | 0.00089991 | 0 |
| 235 | 0 | 0 |
| 240 | 0 | 0 |
| 245 | 0 | 0 |
| 250 | 0 | 0 |
| 255 | 0 | 0 |
| 260 | 0 | 0 |
| 265 | 0 | 0 |
| 270 | 0 | 0 |
| 275 | 0 | 0 |
| 280 | 0 | 0 |
| 285 | 0 | 0 |
| 290 | 0 | 0 |
| 295 | 0 | 0 |
| 300 | 0 | 0 |
| 305 | 0 | 0 |
| 310 | 0 | 0 |
| 315 | 0 | 0 |
| 320 | 0 | 0 |
| 325 | 0 | 0 |
| 330 | 0 | 0 |
| 335 | 0 | 0 |
| 340 | 0 | 0 |
| 345 | 0 | 0 |
| 350 | 0 | 0 |
| 355 | 0 | 0 |
| 360 | 0 | 0 |

(e) 293K-apoAB-eta-U47-J2/3

H(η)

| η | NF-apoA | NF-apoB |
| --- | --- | --- |
| 0 | 0 | 0 |
| 5 | 0 | 0 |
| 10 | 0 | 0 |
| 15 | 0 | 0 |
| 20 | 0 | 0 |
| 25 | 0 | 0 |
| 30 | 0 | 0 |
| 35 | 0 | 0 |
| 40 | 0 | 0 |
| 45 | 0 | 0 |
| 50 | 0 | 0 |
| 55 | 0 | 0 |
| 60 | 0 | 0 |
| 65 | 0 | 0 |
| 70 | 0 | 0 |
| 75 | 0 | 0 |
| 80 | 0 | 0 |
| 85 | 0 | 0 |
| 90 | 0 | 0 |
| 95 | 0 | 0 |
| 100 | 0 | 0 |
| 105 | 0 | 0 |
| 110 | 0 | 0 |
| 115 | 0 | 0 |
| 120 | 0 | 0 |
| 125 | 0 | 0 |
| 130 | 0 | 0.00039996 |
| 135 | 0 | 0.00139986 |
| 140 | 0 | 0.00689931 |
| 145 | 0 | 0.0214979 |
| 150 | 0 | 0.0558944 |
| 155 | 0 | 0.10129 |
| 160 | 9.999e-05 | 0.164784 |
| 165 | 9.999e-05 | 0.193781 |
| 170 | 0.00019998 | 0.172483 |
| 175 | 0.00019998 | 0.10149 |
| 180 | 0.00159984 | 0.0457954 |
| 185 | 0.0149985 | 0.0227977 |
| 190 | 0.0735926 | 0.0213979 |
| 195 | 0.211079 | 0.0290971 |
| 200 | 0.313069 | 0.0311969 |
| 205 | 0.241876 | 0.0205979 |
| 210 | 0.111289 | 0.0069993 |
| 215 | 0.0271973 | 0.00209979 |
| 220 | 0.00409959 | 9.999e-05 |
| 225 | 0.00059994 | 0 |
| 230 | 0 | 0 |
| 235 | 0 | 0 |
| 240 | 0 | 0 |
| 245 | 0 | 0 |
| 250 | 0 | 0 |
| 255 | 0 | 0 |
| 260 | 0 | 0 |
| 265 | 0 | 0 |
| 270 | 0 | 0 |
| 275 | 0 | 0 |
| 280 | 0 | 0 |
| 285 | 0 | 0 |
| 290 | 0 | 0 |
| 295 | 0 | 0 |
| 300 | 0 | 0 |
| 305 | 0 | 0 |
| 310 | 0 | 0 |
| 315 | 0 | 0 |
| 320 | 0 | 0 |
| 325 | 0 | 0 |
| 330 | 0 | 0 |
| 335 | 0 | 0 |
| 340 | 0 | 0 |
| 345 | 0 | 0 |
| 350 | 0 | 0 |
| 355 | 0 | 0 |
| 360 | 0 | 0 |

(f) 293K-apoAB-eta-U48-J2/3

H(η)

| η | NF-apoA | NF-apoB |
| --- | --- | --- |
| 0 | 0.00139986 | 0.00029997 |
| 5 | 0.123688 | 0.0109989 |
| 10 | 0.227277 | 0.0135986 |
| 15 | 0.282272 | 0.0131987 |
| 20 | 0.20138 | 0.0109989 |
| 25 | 0.0790921 | 0.0111989 |
| 30 | 0.020398 | 0.010299 |
| 35 | 0.00379962 | 0.00889911 |
| 40 | 0.00069993 | 0.0121988 |
| 45 | 0.00029997 | 0.0190981 |
| 50 | 0.00019998 | 0.0331967 |
| 55 | 0.00019998 | 0.0576942 |
| 60 | 9.999e-05 | 0.090391 |
| 65 | 0 | 0.127187 |
| 70 | 0.00019998 | 0.141886 |
| 75 | 9.999e-05 | 0.143986 |
| 80 | 0 | 0.114189 |
| 85 | 0 | 0.0673933 |
| 90 | 9.999e-05 | 0.0316968 |
| 95 | 0 | 0.00909909 |
| 100 | 0 | 0.00249975 |
| 105 | 0 | 0.00049995 |
| 110 | 0 | 0 |
| 115 | 0 | 0 |
| 120 | 0 | 0 |
| 125 | 0 | 0 |
| 130 | 0 | 0 |
| 135 | 0 | 0 |
| 140 | 0 | 0 |
| 145 | 0 | 0 |
| 150 | 0 | 0 |
| 155 | 0 | 0 |
| 160 | 0 | 0 |
| 165 | 0 | 0 |
| 170 | 0 | 0 |
| 175 | 0 | 0 |
| 180 | 0 | 0 |
| 185 | 0 | 0 |
| 190 | 0 | 0 |
| 195 | 0 | 0 |
| 200 | 0 | 0 |
| 205 | 0 | 0 |
| 210 | 0 | 0 |
| 215 | 0 | 0 |
| 220 | 0 | 0 |
| 225 | 0 | 0 |
| 230 | 0 | 0 |
| 235 | 0 | 0 |
| 240 | 0 | 0 |
| 245 | 0 | 0 |
| 250 | 0 | 0 |
| 255 | 0 | 0 |
| 260 | 0 | 0 |
| 265 | 0 | 0 |
| 270 | 0 | 0 |
| 275 | 0 | 0 |
| 280 | 0 | 0 |
| 285 | 0 | 0 |
| 290 | 0 | 0 |
| 295 | 0 | 0 |
| 300 | 0 | 0.00019998 |
| 305 | 0 | 0.00029997 |
| 310 | 9.999e-05 | 0.00109989 |
| 315 | 0 | 0.00209979 |
| 320 | 9.999e-05 | 0.00319968 |
| 325 | 0 | 0.0059994 |
| 330 | 0 | 0.00489951 |
| 335 | 9.999e-05 | 0.00569943 |
| 340 | 0.00039996 | 0.0089991 |
| 345 | 0.00109989 | 0.00739926 |
| 350 | 0.00309969 | 0.00849915 |
| 355 | 0.0117988 | 0.0105989 |
| 360 | 0.0420958 | 0.010499 |

(g) 293K-apoAB-eta-U51-J2/3

H(η)

| η | NF-apoA | NF-apoB |
| --- | --- | --- |
| 0 | 0.00289971 | 0.00039996 |
| 5 | 0.155584 | 0.0217978 |
| 10 | 0.144186 | 0.0208979 |
| 15 | 0.115988 | 0.0194981 |
| 20 | 0.079792 | 0.0189981 |
| 25 | 0.0554945 | 0.0237976 |
| 30 | 0.0373963 | 0.0293971 |
| 35 | 0.0184982 | 0.0422958 |
| 40 | 0.0069993 | 0.039596 |
| 45 | 0.00249975 | 0.0216978 |
| 50 | 0.00029997 | 0.00809919 |
| 55 | 9.999e-05 | 0.00149985 |
| 60 | 0 | 0 |
| 65 | 0 | 0 |
| 70 | 0 | 0 |
| 75 | 0 | 0 |
| 80 | 0 | 0 |
| 85 | 0 | 0 |
| 90 | 0 | 0 |
| 95 | 0 | 0 |
| 100 | 0 | 0 |
| 105 | 0 | 0 |
| 110 | 0 | 0 |
| 115 | 0 | 0 |
| 120 | 0 | 0 |
| 125 | 0 | 0 |
| 130 | 0 | 0 |
| 135 | 0 | 0 |
| 140 | 0 | 0 |
| 145 | 0 | 0 |
| 150 | 0 | 0 |
| 155 | 0 | 0 |
| 160 | 0 | 0 |
| 165 | 0 | 0 |
| 170 | 0 | 0 |
| 175 | 0 | 0 |
| 180 | 0 | 0 |
| 185 | 0 | 0 |
| 190 | 0 | 0 |
| 195 | 0 | 0 |
| 200 | 0 | 0 |
| 205 | 0 | 0 |
| 210 | 0 | 9.999e-05 |
| 215 | 0 | 0 |
| 220 | 0 | 0.00019998 |
| 225 | 0 | 0.00029997 |
| 230 | 0 | 0.00069993 |
| 235 | 0 | 0.00109989 |
| 240 | 0 | 0.0009999 |
| 245 | 0 | 0.00049995 |
| 250 | 0 | 0.00109989 |
| 255 | 0 | 0.0009999 |
| 260 | 0 | 0.00489951 |
| 265 | 0 | 0.0123988 |
| 270 | 0 | 0.0220978 |
| 275 | 0 | 0.0222978 |
| 280 | 0 | 0.0166983 |
| 285 | 0 | 0.00989901 |
| 290 | 0 | 0.00979902 |
| 295 | 0 | 0.0147985 |
| 300 | 0 | 0.0248975 |
| 305 | 0 | 0.0380962 |
| 310 | 0 | 0.0556944 |
| 315 | 9.999e-05 | 0.0707929 |
| 320 | 0.00029997 | 0.0805919 |
| 325 | 0.00089991 | 0.0776922 |
| 330 | 0.00379962 | 0.0724928 |
| 335 | 0.00989901 | 0.059694 |
| 340 | 0.0222978 | 0.0434957 |
| 345 | 0.0378962 | 0.0359964 |
| 350 | 0.0621938 | 0.0282972 |
| 355 | 0.10379 | 0.0225977 |
| 360 | 0.139086 | 0.0227977 |

(h) 293K-apoAB-eta-A52-J2/3

H(η)

| η | NF-apoA | NF-apoB |
| --- | --- | --- |
| 0 | 0 | 0 |
| 5 | 0 | 0 |
| 10 | 0 | 0 |
| 15 | 0 | 0 |
| 20 | 0 | 0 |
| 25 | 0 | 0 |
| 30 | 0 | 0 |
| 35 | 0 | 0 |
| 40 | 0 | 0 |
| 45 | 0 | 0 |
| 50 | 0 | 0 |
| 55 | 0 | 0 |
| 60 | 0 | 0 |
| 65 | 0 | 0 |
| 70 | 0 | 0 |
| 75 | 0 | 0 |
| 80 | 0 | 0 |
| 85 | 0 | 0 |
| 90 | 0 | 0 |
| 95 | 0 | 0 |
| 100 | 0 | 9.999e-05 |
| 105 | 0 | 0.00049995 |
| 110 | 0 | 0.00119988 |
| 115 | 0 | 0.00279972 |
| 120 | 0 | 0.00489951 |
| 125 | 0 | 0.00819918 |
| 130 | 0 | 0.00949905 |
| 135 | 0 | 0.0122988 |
| 140 | 0 | 0.0170983 |
| 145 | 0 | 0.019598 |
| 150 | 0 | 0.0235976 |
| 155 | 0 | 0.0310969 |
| 160 | 0 | 0.0443956 |
| 165 | 9.999e-05 | 0.0565943 |
| 170 | 0.00149985 | 0.0654935 |
| 175 | 0.009999 | 0.0755924 |
| 180 | 0.0374963 | 0.069793 |
| 185 | 0.0970903 | 0.0627937 |
| 190 | 0.168383 | 0.0459954 |
| 195 | 0.20038 | 0.040096 |
| 200 | 0.182982 | 0.0429957 |
| 205 | 0.140086 | 0.059894 |
| 210 | 0.0929907 | 0.0707929 |
| 215 | 0.0458954 | 0.0785921 |
| 220 | 0.0171983 | 0.0630937 |
| 225 | 0.00489951 | 0.0457954 |
| 230 | 0.00059994 | 0.0247975 |
| 235 | 0.00039996 | 0.0148985 |
| 240 | 0 | 0.00539946 |
| 245 | 0 | 0.00149985 |
| 250 | 0 | 0.00039996 |
| 255 | 0 | 0.00019998 |
| 260 | 0 | 0 |
| 265 | 0 | 0 |
| 270 | 0 | 0 |
| 275 | 0 | 0 |
| 280 | 0 | 0 |
| 285 | 0 | 0 |
| 290 | 0 | 0 |
| 295 | 0 | 0 |
| 300 | 0 | 0 |
| 305 | 0 | 0 |
| 310 | 0 | 0 |
| 315 | 0 | 0 |
| 320 | 0 | 0 |
| 325 | 0 | 0 |
| 330 | 0 | 0 |
| 335 | 0 | 0 |
| 340 | 0 | 0 |
| 345 | 0 | 0 |
| 350 | 0 | 0 |
| 355 | 0 | 0 |
| 360 | 0 | 0 |

**Figure S8: Some representative Histogram distribution of pseudo-torsion angle (θ) at 293K (apoA in blue and apoB in orange))**

(a) 293K-apoAB-theta-U22-J1/2

H(θ)

| θ | NF-apoA | NF-apoB |
| --- | --- | --- |
| 0 | 0 | 0.00279972 |
| 5 | 0 | 0.142286 |
| 10 | 0 | 0.154485 |
| 15 | 0 | 0.150785 |
| 20 | 0 | 0.123588 |
| 25 | 0 | 0.0634937 |
| 30 | 0 | 0.0172983 |
| 35 | 0 | 0.0049995 |
| 40 | 0 | 0.00089991 |
| 45 | 0 | 9.999e-05 |
| 50 | 0 | 0 |
| 55 | 0 | 0 |
| 60 | 0 | 0 |
| 65 | 0 | 0 |
| 70 | 0 | 0 |
| 75 | 0 | 0 |
| 80 | 0 | 0 |
| 85 | 0 | 0 |
| 90 | 0 | 0 |
| 95 | 0.00019998 | 0 |
| 100 | 0.00569943 | 0 |
| 105 | 0.0573943 | 0 |
| 110 | 0.227077 | 0 |
| 115 | 0.363164 | 0 |
| 120 | 0.253975 | 0 |
| 125 | 0.0785921 | 0 |
| 130 | 0.0129987 | 0 |
| 135 | 0.00079992 | 0 |
| 140 | 9.999e-05 | 0 |
| 145 | 0 | 0 |
| 150 | 0 | 0 |
| 155 | 0 | 0 |
| 160 | 0 | 0 |
| 165 | 0 | 0 |
| 170 | 0 | 0 |
| 175 | 0 | 0 |
| 180 | 0 | 0 |
| 185 | 0 | 0 |
| 190 | 0 | 0 |
| 195 | 0 | 0 |
| 200 | 0 | 0 |
| 205 | 0 | 0 |
| 210 | 0 | 0 |
| 215 | 0 | 0 |
| 220 | 0 | 0 |
| 225 | 0 | 0 |
| 230 | 0 | 0 |
| 235 | 0 | 0 |
| 240 | 0 | 0 |
| 245 | 0 | 0 |
| 250 | 0 | 0 |
| 255 | 0 | 0 |
| 260 | 0 | 0 |
| 265 | 0 | 0 |
| 270 | 0 | 0 |
| 275 | 0 | 0 |
| 280 | 0 | 0 |
| 285 | 0 | 0 |
| 290 | 0 | 0 |
| 295 | 0 | 0.00019998 |
| 300 | 0 | 0.00029997 |
| 305 | 0 | 0.00019998 |
| 310 | 0 | 0.00029997 |
| 315 | 0 | 0.00019998 |
| 320 | 0 | 0.00109989 |
| 325 | 0 | 9.999e-05 |
| 330 | 0 | 0.00049995 |
| 335 | 0 | 0.0009999 |
| 340 | 0 | 0.00469953 |
| 345 | 0 | 0.0156984 |
| 350 | 0 | 0.0472953 |
| 355 | 0 | 0.114389 |
| 360 | 0 | 0.153285 |

(b) 293K-apoAB-theta-A23-J1/2

H(θ)

| θ | NF-apoA | NF-apoB |
| --- | --- | --- |
| 0 | 0.00329967 | 0 |
| 5 | 0.114889 | 0 |
| 10 | 0.0590941 | 0 |
| 15 | 0.0247975 | 0 |
| 20 | 0.00989901 | 0 |
| 25 | 0.00379962 | 0 |
| 30 | 0.00069993 | 0 |
| 35 | 0.00019998 | 0 |
| 40 | 0 | 0 |
| 45 | 0 | 0 |
| 50 | 0 | 0 |
| 55 | 0 | 0 |
| 60 | 0 | 0 |
| 65 | 0 | 0 |
| 70 | 0 | 0 |
| 75 | 0 | 0 |
| 80 | 0 | 0 |
| 85 | 0 | 0 |
| 90 | 0 | 0 |
| 95 | 0 | 0 |
| 100 | 0 | 0 |
| 105 | 0 | 0 |
| 110 | 0 | 0 |
| 115 | 0 | 0 |
| 120 | 0 | 0 |
| 125 | 0 | 0 |
| 130 | 0 | 0 |
| 135 | 0 | 0 |
| 140 | 0 | 0 |
| 145 | 0 | 0.00019998 |
| 150 | 0 | 0.00179982 |
| 155 | 0 | 0.00459954 |
| 160 | 0 | 0.00739926 |
| 165 | 0 | 0.00849915 |
| 170 | 0 | 0.0240976 |
| 175 | 0 | 0.0780922 |
| 180 | 0 | 0.171583 |
| 185 | 0 | 0.231277 |
| 190 | 0 | 0.188781 |
| 195 | 0 | 0.141086 |
| 200 | 0 | 0.0844916 |
| 205 | 0 | 0.039896 |
| 210 | 0 | 0.0141986 |
| 215 | 0 | 0.00309969 |
| 220 | 0 | 0.00049995 |
| 225 | 0 | 0.00039996 |
| 230 | 0 | 0 |
| 235 | 0 | 0 |
| 240 | 0 | 0 |
| 245 | 0 | 0 |
| 250 | 0 | 0 |
| 255 | 0 | 0 |
| 260 | 0 | 0 |
| 265 | 0 | 0 |
| 270 | 0 | 0 |
| 275 | 0 | 0 |
| 280 | 0 | 0 |
| 285 | 0 | 0 |
| 290 | 9.999e-05 | 0 |
| 295 | 0.00019998 | 0 |
| 300 | 0.00059994 | 0 |
| 305 | 0.0009999 | 0 |
| 310 | 0.00169983 | 0 |
| 315 | 0.00169983 | 0 |
| 320 | 0.00629937 | 0 |
| 325 | 0.0119988 | 0 |
| 330 | 0.0281972 | 0 |
| 335 | 0.0575942 | 0 |
| 340 | 0.0865913 | 0 |
| 345 | 0.111789 | 0 |
| 350 | 0.142686 | 0 |
| 355 | 0.173883 | 0 |
| 360 | 0.158984 | 0 |

(c) 293K-apoAB-theta-A24-J1/2

H(θ)

| θ | NF-apoA | NF-apoB |
| --- | --- | --- |
| 0 | 0 | 0 |
| 5 | 0 | 0 |
| 10 | 0 | 0 |
| 15 | 0 | 0 |
| 20 | 0 | 0 |
| 25 | 0 | 0 |
| 30 | 0.00039996 | 0 |
| 35 | 0.00459954 | 0 |
| 40 | 0.0388961 | 0.00049995 |
| 45 | 0.175382 | 0.00659934 |
| 50 | 0.318968 | 0.0381962 |
| 55 | 0.280772 | 0.089691 |
| 60 | 0.118988 | 0.142286 |
| 65 | 0.0390961 | 0.142286 |
| 70 | 0.0160984 | 0.116388 |
| 75 | 0.00519948 | 0.0632937 |
| 80 | 0.00159984 | 0.0323968 |
| 85 | 0 | 0.0234977 |
| 90 | 0 | 0.0341966 |
| 95 | 0 | 0.0555944 |
| 100 | 0 | 0.0652935 |
| 105 | 0 | 0.0651935 |
| 110 | 0 | 0.0487951 |
| 115 | 0 | 0.0416958 |
| 120 | 0 | 0.0243976 |
| 125 | 0 | 0.00849915 |
| 130 | 0 | 0.00079992 |
| 135 | 0 | 0.00039996 |
| 140 | 0 | 0 |
| 145 | 0 | 0 |
| 150 | 0 | 0 |
| 155 | 0 | 0 |
| 160 | 0 | 0 |
| 165 | 0 | 0 |
| 170 | 0 | 0 |
| 175 | 0 | 0 |
| 180 | 0 | 0 |
| 185 | 0 | 0 |
| 190 | 0 | 0 |
| 195 | 0 | 0 |
| 200 | 0 | 0 |
| 205 | 0 | 0 |
| 210 | 0 | 0 |
| 215 | 0 | 0 |
| 220 | 0 | 0 |
| 225 | 0 | 0 |
| 230 | 0 | 0 |
| 235 | 0 | 0 |
| 240 | 0 | 0 |
| 245 | 0 | 0 |
| 250 | 0 | 0 |
| 255 | 0 | 0 |
| 260 | 0 | 0 |
| 265 | 0 | 0 |
| 270 | 0 | 0 |
| 275 | 0 | 0 |
| 280 | 0 | 0 |
| 285 | 0 | 0 |
| 290 | 0 | 0 |
| 295 | 0 | 0 |
| 300 | 0 | 0 |
| 305 | 0 | 0 |
| 310 | 0 | 0 |
| 315 | 0 | 0 |
| 320 | 0 | 0 |
| 325 | 0 | 0 |
| 330 | 0 | 0 |
| 335 | 0 | 0 |
| 340 | 0 | 0 |
| 345 | 0 | 0 |
| 350 | 0 | 0 |
| 355 | 0 | 0 |
| 360 | 0 | 0 |

(d) 293K-apoAB-theta-U48-J2/3

H(θ)

| θ | NF-apoA | NF-apoB |
| --- | --- | --- |
| 0 | 0 | 0 |
| 5 | 0 | 0 |
| 10 | 0 | 0 |
| 15 | 0 | 0 |
| 20 | 0 | 0 |
| 25 | 0 | 0 |
| 30 | 0 | 0 |
| 35 | 0 | 0 |
| 40 | 0 | 0 |
| 45 | 0 | 0 |
| 50 | 0 | 0 |
| 55 | 0 | 0 |
| 60 | 0 | 0 |
| 65 | 0 | 0 |
| 70 | 0 | 0 |
| 75 | 0 | 0 |
| 80 | 0 | 0 |
| 85 | 0 | 0 |
| 90 | 0 | 0 |
| 95 | 0 | 0 |
| 100 | 0 | 0 |
| 105 | 0 | 0 |
| 110 | 0 | 0 |
| 115 | 0 | 0 |
| 120 | 0 | 0 |
| 125 | 0 | 0 |
| 130 | 0 | 0 |
| 135 | 0 | 0 |
| 140 | 0 | 0 |
| 145 | 0 | 0.00029997 |
| 150 | 0 | 0.00079992 |
| 155 | 0 | 0.00119988 |
| 160 | 0 | 0.00529947 |
| 165 | 0 | 0.0155984 |
| 170 | 0 | 0.0385961 |
| 175 | 0 | 0.0568943 |
| 180 | 0 | 0.0593941 |
| 185 | 0.00069993 | 0.0581942 |
| 190 | 0.00209979 | 0.0493951 |
| 195 | 0.00929907 | 0.0419958 |
| 200 | 0.0432957 | 0.0418958 |
| 205 | 0.20498 | 0.0405959 |
| 210 | 0.377362 | 0.0534947 |
| 215 | 0.268773 | 0.0564944 |
| 220 | 0.0753925 | 0.0549945 |
| 225 | 0.0134987 | 0.0632937 |
| 230 | 0.00239976 | 0.070093 |
| 235 | 0.00089991 | 0.0749925 |
| 240 | 0.00089991 | 0.070293 |
| 245 | 0.00029997 | 0.0513949 |
| 250 | 9.999e-05 | 0.030497 |
| 255 | 0 | 0.0168983 |
| 260 | 0 | 0.0111989 |
| 265 | 0 | 0.00769923 |
| 270 | 0 | 0.00529947 |
| 275 | 0 | 0.00519948 |
| 280 | 0 | 0.00439956 |
| 285 | 0 | 0.00429957 |
| 290 | 0 | 0.00269973 |
| 295 | 0 | 0.00309969 |
| 300 | 0 | 0.00209979 |
| 305 | 0 | 0.00059994 |
| 310 | 0 | 0.00039996 |
| 315 | 0 | 0.00029997 |
| 320 | 0 | 9.999e-05 |
| 325 | 0 | 0 |
| 330 | 0 | 0 |
| 335 | 0 | 0 |
| 340 | 0 | 0 |
| 345 | 0 | 0 |
| 350 | 0 | 0 |
| 355 | 0 | 0 |
| 360 | 0 | 0 |

(e) 293K-apoAB-theta-C50-J2/3

H(θ)

| θ | NF-apoA | NF-apoB |
| --- | --- | --- |
| 0 | 0 | 0 |
| 5 | 0 | 0 |
| 10 | 0 | 0 |
| 15 | 0 | 0 |
| 20 | 0 | 0 |
| 25 | 0 | 0 |
| 30 | 0 | 0 |
| 35 | 0 | 0 |
| 40 | 0 | 0 |
| 45 | 0 | 0 |
| 50 | 0 | 0 |
| 55 | 0 | 0 |
| 60 | 9.999e-05 | 0 |
| 65 | 0.00049995 | 9.999e-05 |
| 70 | 0.00119988 | 0.00019998 |
| 75 | 0.00289971 | 0.00129987 |
| 80 | 0.00319968 | 0.00159984 |
| 85 | 0.00289971 | 0.00549945 |
| 90 | 0.00469953 | 0.0177982 |
| 95 | 0.00729927 | 0.0323968 |
| 100 | 0.0166983 | 0.0455954 |
| 105 | 0.0275972 | 0.0437956 |
| 110 | 0.039696 | 0.0372963 |
| 115 | 0.0494951 | 0.0261974 |
| 120 | 0.0483952 | 0.0169983 |
| 125 | 0.060394 | 0.0135986 |
| 130 | 0.0950905 | 0.0150985 |
| 135 | 0.131487 | 0.0135986 |
| 140 | 0.151485 | 0.0162984 |
| 145 | 0.138686 | 0.0181982 |
| 150 | 0.10289 | 0.0242976 |
| 155 | 0.0623938 | 0.0313969 |
| 160 | 0.0318968 | 0.029997 |
| 165 | 0.0133987 | 0.0322968 |
| 170 | 0.00439956 | 0.0273973 |
| 175 | 0.00229977 | 0.0186981 |
| 180 | 0.00059994 | 0.0150985 |
| 185 | 0.00029997 | 0.0163984 |
| 190 | 0 | 0.0218978 |
| 195 | 0 | 0.0291971 |
| 200 | 0 | 0.0381962 |
| 205 | 0 | 0.0561944 |
| 210 | 0 | 0.0635936 |
| 215 | 0 | 0.0693931 |
| 220 | 0 | 0.0647935 |
| 225 | 0 | 0.0516948 |
| 230 | 0 | 0.040096 |
| 235 | 0 | 0.0236976 |
| 240 | 0 | 0.0172983 |
| 245 | 0 | 0.00919908 |
| 250 | 0 | 0.00619938 |
| 255 | 0 | 0.00269973 |
| 260 | 0 | 0.0019998 |
| 265 | 0 | 0.00129987 |
| 270 | 0 | 0.0009999 |
| 275 | 0 | 0.00019998 |
| 280 | 0 | 0.00019998 |
| 285 | 0 | 0 |
| 290 | 0 | 0 |
| 295 | 0 | 0 |
| 300 | 0 | 0 |
| 305 | 0 | 0 |
| 310 | 0 | 0 |
| 315 | 0 | 0 |
| 320 | 0 | 0 |
| 325 | 0 | 0 |
| 330 | 0 | 0 |
| 335 | 0 | 0 |
| 340 | 0 | 0 |
| 345 | 0 | 0 |
| 350 | 0 | 0 |
| 355 | 0 | 0 |
| 360 | 0 | 0 |

(f) 293K-apoAB-theta-U51-J2/3

H(θ)

| θ | NF-apoA | NF-apoB |
| --- | --- | --- |
| 0 | 0.00039996 | 0.00239976 |
| 5 | 0.00909909 | 0.0935906 |
| 10 | 0.00379962 | 0.0808919 |
| 15 | 0.00049995 | 0.0605939 |
| 20 | 0.00059994 | 0.0418958 |
| 25 | 0 | 0.0331967 |
| 30 | 0 | 0.0207979 |
| 35 | 0 | 0.0215978 |
| 40 | 0 | 0.010199 |
| 45 | 0 | 0.00979902 |
| 50 | 0 | 0.00729927 |
| 55 | 0 | 0.00809919 |
| 60 | 0 | 0.0109989 |
| 65 | 0 | 0.0145985 |
| 70 | 0 | 0.0206979 |
| 75 | 0 | 0.0169983 |
| 80 | 0 | 0.010199 |
| 85 | 0 | 0.00479952 |
| 90 | 0 | 0.00229977 |
| 95 | 0 | 0.00089991 |
| 100 | 0 | 0.00029997 |
| 105 | 0 | 0.00039996 |
| 110 | 0 | 0 |
| 115 | 0 | 0 |
| 120 | 0 | 0 |
| 125 | 0 | 9.999e-05 |
| 130 | 0 | 0 |
| 135 | 0 | 0 |
| 140 | 0 | 0 |
| 145 | 0 | 0 |
| 150 | 0 | 0 |
| 155 | 0 | 0 |
| 160 | 0 | 0 |
| 165 | 0 | 0 |
| 170 | 0 | 0 |
| 175 | 0 | 0 |
| 180 | 0 | 0 |
| 185 | 0 | 0 |
| 190 | 0 | 0 |
| 195 | 0 | 0 |
| 200 | 0 | 0 |
| 205 | 0 | 0 |
| 210 | 0 | 0 |
| 215 | 0 | 0 |
| 220 | 0 | 0 |
| 225 | 0 | 0 |
| 230 | 0 | 0 |
| 235 | 0 | 0 |
| 240 | 0 | 0 |
| 245 | 0 | 0 |
| 250 | 0 | 0 |
| 255 | 0 | 0 |
| 260 | 0 | 0 |
| 265 | 0 | 0 |
| 270 | 0.00049995 | 0 |
| 275 | 0.00069993 | 0 |
| 280 | 0.00159984 | 0 |
| 285 | 0.00309969 | 0.00059994 |
| 290 | 0.00789921 | 0.00289971 |
| 295 | 0.0132987 | 0.00659934 |
| 300 | 0.0237976 | 0.0141986 |
| 305 | 0.0370963 | 0.0225977 |
| 310 | 0.0538946 | 0.030397 |
| 315 | 0.0761924 | 0.0372963 |
| 320 | 0.0914909 | 0.0380962 |
| 325 | 0.108889 | 0.0321968 |
| 330 | 0.120788 | 0.0314969 |
| 335 | 0.118788 | 0.0219978 |
| 340 | 0.112089 | 0.0254975 |
| 345 | 0.089591 | 0.0316968 |
| 350 | 0.0662934 | 0.0556944 |
| 355 | 0.0370963 | 0.0806919 |
| 360 | 0.0224978 | 0.0953905 |

(g) 293K-apoAB-theta-C61-Loop3

H(θ)

| θ | NF-apoA | NF-apoB |
| --- | --- | --- |
| 0 | 0 | 0 |
| 5 | 0 | 0 |
| 10 | 0 | 0 |
| 15 | 0 | 0 |
| 20 | 0 | 0 |
| 25 | 0 | 0 |
| 30 | 0 | 0 |
| 35 | 0 | 0 |
| 40 | 0 | 0 |
| 45 | 0 | 0 |
| 50 | 0 | 0 |
| 55 | 0 | 0 |
| 60 | 0 | 0 |
| 65 | 0 | 0 |
| 70 | 0 | 0 |
| 75 | 0 | 0 |
| 80 | 0 | 0 |
| 85 | 0 | 0 |
| 90 | 0 | 0 |
| 95 | 0 | 0 |
| 100 | 0 | 0 |
| 105 | 0 | 0 |
| 110 | 0 | 0 |
| 115 | 0 | 0 |
| 120 | 0 | 0 |
| 125 | 0 | 0 |
| 130 | 0 | 0 |
| 135 | 0 | 0 |
| 140 | 0 | 9.999e-05 |
| 145 | 0 | 0.00019998 |
| 150 | 0 | 0.00139986 |
| 155 | 0 | 0.00389961 |
| 160 | 0 | 0.00619938 |
| 165 | 0 | 0.0179982 |
| 170 | 0 | 0.0312969 |
| 175 | 0 | 0.0441956 |
| 180 | 0 | 0.0439956 |
| 185 | 0 | 0.0341966 |
| 190 | 0 | 0.0241976 |
| 195 | 0 | 0.0176982 |
| 200 | 9.999e-05 | 0.0145985 |
| 205 | 9.999e-05 | 0.0206979 |
| 210 | 0.00019998 | 0.0224978 |
| 215 | 0.00239976 | 0.0243976 |
| 220 | 0.0114989 | 0.0260974 |
| 225 | 0.0364964 | 0.0338966 |
| 230 | 0.0910909 | 0.0472953 |
| 235 | 0.170083 | 0.0727927 |
| 240 | 0.233177 | 0.10139 |
| 245 | 0.214979 | 0.116988 |
| 250 | 0.137286 | 0.10309 |
| 255 | 0.0672933 | 0.080292 |
| 260 | 0.0255974 | 0.0471953 |
| 265 | 0.00849915 | 0.0269973 |
| 270 | 0.00109989 | 0.0140986 |
| 275 | 9.999e-05 | 0.00749925 |
| 280 | 0 | 0.00619938 |
| 285 | 0 | 0.00289971 |
| 290 | 0 | 0.00369963 |
| 295 | 0 | 0.00109989 |
| 300 | 0 | 0.00059994 |
| 305 | 0 | 0.00019998 |
| 310 | 0 | 9.999e-05 |
| 315 | 0 | 0 |
| 320 | 0 | 0 |
| 325 | 0 | 0 |
| 330 | 0 | 0 |
| 335 | 0 | 0 |
| 340 | 0 | 0 |
| 345 | 0 | 0 |
| 350 | 0 | 0 |
| 355 | 0 | 0 |
| 360 | 0 | 0 |

(h) 293K-apoAB-theta-U63-Loop3

H(θ)

| θ | NF-apoA | NF-apoB |
| --- | --- | --- |
| 0 | 0 | 0 |
| 5 | 0 | 0 |
| 10 | 0 | 0 |
| 15 | 0 | 0 |
| 20 | 0 | 0.00019998 |
| 25 | 0 | 0.00049995 |
| 30 | 0 | 0.00479952 |
| 35 | 0 | 0.0275972 |
| 40 | 0 | 0.0626937 |
| 45 | 0 | 0.0756924 |
| 50 | 0 | 0.070093 |
| 55 | 0 | 0.039696 |
| 60 | 0 | 0.0185981 |
| 65 | 0 | 0.00949905 |
| 70 | 9.999e-05 | 0.00539946 |
| 75 | 0.00239976 | 0.00829917 |
| 80 | 0.0113989 | 0.0216978 |
| 85 | 0.0447955 | 0.0439956 |
| 90 | 0.0966903 | 0.0810919 |
| 95 | 0.171283 | 0.122788 |
| 100 | 0.20128 | 0.124888 |
| 105 | 0.192181 | 0.10409 |
| 110 | 0.137686 | 0.0774923 |
| 115 | 0.0810919 | 0.0463954 |
| 120 | 0.0388961 | 0.0279972 |
| 125 | 0.0128987 | 0.0129987 |
| 130 | 0.00549945 | 0.0079992 |
| 135 | 0.00179982 | 0.00369963 |
| 140 | 0.00159984 | 0.00129987 |
| 145 | 0.00029997 | 0.00049995 |
| 150 | 9.999e-05 | 0 |
| 155 | 0 | 0 |
| 160 | 0 | 0 |
| 165 | 0 | 0 |
| 170 | 0 | 0 |
| 175 | 0 | 0 |
| 180 | 0 | 0 |
| 185 | 0 | 0 |
| 190 | 0 | 0 |
| 195 | 0 | 0 |
| 200 | 0 | 0 |
| 205 | 0 | 0 |
| 210 | 0 | 0 |
| 215 | 0 | 0 |
| 220 | 0 | 0 |
| 225 | 0 | 0 |
| 230 | 0 | 0 |
| 235 | 0 | 0 |
| 240 | 0 | 0 |
| 245 | 0 | 0 |
| 250 | 0 | 0 |
| 255 | 0 | 0 |
| 260 | 0 | 0 |
| 265 | 0 | 0 |
| 270 | 0 | 0 |
| 275 | 0 | 0 |
| 280 | 0 | 0 |
| 285 | 0 | 0 |
| 290 | 0 | 0 |
| 295 | 0 | 0 |
| 300 | 0 | 0 |
| 305 | 0 | 0 |
| 310 | 0 | 0 |
| 315 | 0 | 0 |
| 320 | 0 | 0 |
| 325 | 0 | 0 |
| 330 | 0 | 0 |
| 335 | 0 | 0 |
| 340 | 0 | 0 |
| 345 | 0 | 0 |
| 350 | 0 | 0 |
| 355 | 0 | 0 |
| 360 | 0 | 0 |

(i) 293K-apoAB-theta-U68-Stem2

H(θ)

| θ | NF-apoA | NF-apoB |
| --- | --- | --- |
| 0 | 0 | 0 |
| 5 | 0 | 0 |
| 10 | 0 | 0 |
| 15 | 0 | 0 |
| 20 | 0 | 0 |
| 25 | 0 | 0 |
| 30 | 0 | 0 |
| 35 | 0 | 0 |
| 40 | 0 | 0 |
| 45 | 0 | 0 |
| 50 | 0 | 0 |
| 55 | 0 | 0 |
| 60 | 0 | 0 |
| 65 | 0 | 0 |
| 70 | 0 | 0 |
| 75 | 0 | 0 |
| 80 | 0 | 0 |
| 85 | 0 | 0 |
| 90 | 0 | 0 |
| 95 | 0 | 0 |
| 100 | 0 | 0 |
| 105 | 0 | 0 |
| 110 | 0 | 0 |
| 115 | 0 | 0 |
| 120 | 0 | 0 |
| 125 | 0 | 0 |
| 130 | 0 | 0 |
| 135 | 0 | 0 |
| 140 | 0 | 0 |
| 145 | 0 | 0 |
| 150 | 0 | 0 |
| 155 | 0 | 0 |
| 160 | 0 | 0 |
| 165 | 0 | 0 |
| 170 | 0 | 0.00059994 |
| 175 | 0.00019998 | 0.00339966 |
| 180 | 0.00149985 | 0.0166983 |
| 185 | 0.00429957 | 0.0520948 |
| 190 | 0.0165983 | 0.0893911 |
| 195 | 0.0455954 | 0.108689 |
| 200 | 0.10269 | 0.107489 |
| 205 | 0.175282 | 0.106389 |
| 210 | 0.213879 | 0.114489 |
| 215 | 0.20398 | 0.121788 |
| 220 | 0.140986 | 0.108889 |
| 225 | 0.0651935 | 0.0769923 |
| 230 | 0.019698 | 0.0457954 |
| 235 | 0.00589941 | 0.0179982 |
| 240 | 0.00269973 | 0.00889911 |
| 245 | 0.0009999 | 0.00779922 |
| 250 | 0.00049995 | 0.00659934 |
| 255 | 0 | 0.00359964 |
| 260 | 0 | 0.00159984 |
| 265 | 0 | 0.00059994 |
| 270 | 0 | 9.999e-05 |
| 275 | 0 | 0 |
| 280 | 0 | 9.999e-05 |
| 285 | 0 | 0 |
| 290 | 0 | 0 |
| 295 | 0 | 0 |
| 300 | 0 | 0 |
| 305 | 0 | 0 |
| 310 | 0 | 0 |
| 315 | 0 | 0 |
| 320 | 0 | 0 |
| 325 | 0 | 0 |
| 330 | 0 | 0 |
| 335 | 0 | 0 |
| 340 | 0 | 0 |
| 345 | 0 | 0 |
| 350 | 0 | 0 |
| 355 | 0 | 0 |
| 360 | 0 | 0 |

(j) 293K-apoAB-theta-A73-J3/1

H(θ)

| θ | NF-apoA | NF-apoB |
| --- | --- | --- |
| 0 | 0 | 0 |
| 5 | 0 | 0 |
| 10 | 0 | 0 |
| 15 | 0 | 0 |
| 20 | 0 | 0 |
| 25 | 0 | 0 |
| 30 | 0 | 0 |
| 35 | 0 | 0 |
| 40 | 0 | 0 |
| 45 | 0 | 0 |
| 50 | 0 | 0 |
| 55 | 0 | 0 |
| 60 | 0 | 0 |
| 65 | 0 | 0 |
| 70 | 0 | 0 |
| 75 | 0 | 0 |
| 80 | 0 | 0 |
| 85 | 0 | 0 |
| 90 | 0 | 0 |
| 95 | 0 | 0 |
| 100 | 0 | 0 |
| 105 | 0 | 0 |
| 110 | 0 | 0 |
| 115 | 0 | 0 |
| 120 | 0 | 0 |
| 125 | 0 | 0 |
| 130 | 9.999e-05 | 0 |
| 135 | 0 | 0 |
| 140 | 0.00029997 | 0 |
| 145 | 0.00019998 | 0 |
| 150 | 0.00079992 | 0 |
| 155 | 0.00179982 | 0 |
| 160 | 0.00179982 | 0 |
| 165 | 0.0009999 | 0 |
| 170 | 0.00089991 | 0 |
| 175 | 0.00089991 | 0 |
| 180 | 0.00219978 | 0 |
| 185 | 0.00239976 | 0 |
| 190 | 0.00809919 | 9.999e-05 |
| 195 | 0.020498 | 0.00019998 |
| 200 | 0.0355964 | 0.00089991 |
| 205 | 0.0825917 | 0.00229977 |
| 210 | 0.161084 | 0.00729927 |
| 215 | 0.235876 | 0.0241976 |
| 220 | 0.227477 | 0.0546945 |
| 225 | 0.146085 | 0.112889 |
| 230 | 0.0542946 | 0.152285 |
| 235 | 0.0136986 | 0.146885 |
| 240 | 0.00189981 | 0.123288 |
| 245 | 0.00039996 | 0.116188 |
| 250 | 0 | 0.0974903 |
| 255 | 0 | 0.0818918 |
| 260 | 0 | 0.0510949 |
| 265 | 0 | 0.0207979 |
| 270 | 0 | 0.00649935 |
| 275 | 0 | 0.00079992 |
| 280 | 0 | 9.999e-05 |
| 285 | 0 | 9.999e-05 |
| 290 | 0 | 0 |
| 295 | 0 | 0 |
| 300 | 0 | 0 |
| 305 | 0 | 0 |
| 310 | 0 | 0 |
| 315 | 0 | 0 |
| 320 | 0 | 0 |
| 325 | 0 | 0 |
| 330 | 0 | 0 |
| 335 | 0 | 0 |
| 340 | 0 | 0 |
| 345 | 0 | 0 |
| 350 | 0 | 0 |
| 355 | 0 | 0 |
| 360 | 0 | 0 |

**Figure S9: Some representative Histogram distribution of pseudo-torsion angle (η) at 303K (apoA in blue and apoB in orange))**

(a) 303K-apoAB-eta-A24-J1/2

H(η)

| η | NF-apoA | NF-apoB |
| --- | --- | --- |
| 0 | 0 | 0 |
| 5 | 0 | 0 |
| 10 | 0 | 0 |
| 15 | 0 | 0 |
| 20 | 0 | 0 |
| 25 | 0 | 0 |
| 30 | 0 | 0 |
| 35 | 0 | 0 |
| 40 | 0 | 0 |
| 45 | 0 | 0 |
| 50 | 0 | 0 |
| 55 | 0 | 0 |
| 60 | 0 | 0 |
| 65 | 0 | 0 |
| 70 | 0 | 0 |
| 75 | 0 | 0 |
| 80 | 0 | 0 |
| 85 | 0 | 0 |
| 90 | 0 | 0 |
| 95 | 0 | 0 |
| 100 | 0 | 0 |
| 105 | 0 | 0 |
| 110 | 0 | 0 |
| 115 | 0 | 0 |
| 120 | 0 | 0 |
| 125 | 0 | 0 |
| 130 | 0 | 0 |
| 135 | 0 | 0 |
| 140 | 0 | 0 |
| 145 | 0 | 0 |
| 150 | 0 | 0 |
| 155 | 0 | 0 |
| 160 | 0 | 0.00019998 |
| 165 | 0 | 0.00029997 |
| 170 | 0 | 0.00159984 |
| 175 | 0 | 0.0108989 |
| 180 | 0 | 0.0286971 |
| 185 | 0 | 0.0506949 |
| 190 | 0 | 0.0767923 |
| 195 | 0 | 0.0991901 |
| 200 | 0 | 0.123088 |
| 205 | 0 | 0.144486 |
| 210 | 0 | 0.132187 |
| 215 | 0 | 0.0971903 |
| 220 | 0 | 0.0688931 |
| 225 | 0 | 0.0523948 |
| 230 | 0 | 0.0360964 |
| 235 | 0.00029997 | 0.0233977 |
| 240 | 0.00109989 | 0.0122988 |
| 245 | 0.00709929 | 0.00719928 |
| 250 | 0.0329967 | 0.00479952 |
| 255 | 0.10309 | 0.00529947 |
| 260 | 0.219278 | 0.00959904 |
| 265 | 0.280272 | 0.00749925 |
| 270 | 0.216078 | 0.00439956 |
| 275 | 0.10009 | 0.00229977 |
| 280 | 0.029897 | 0.00029997 |
| 285 | 0.00659934 | 0.00019998 |
| 290 | 0.00219978 | 0 |
| 295 | 0.00089991 | 0 |
| 300 | 9.999e-05 | 0 |
| 305 | 0 | 0 |
| 310 | 0 | 0 |
| 315 | 0 | 0 |
| 320 | 0 | 0 |
| 325 | 0 | 0 |
| 330 | 0 | 0 |
| 335 | 0 | 0 |
| 340 | 0 | 0 |
| 345 | 0 | 0 |
| 350 | 0 | 0 |
| 355 | 0 | 0 |
| 360 | 0 | 0 |

(b) 303K-apoAB-eta-U25-Stem2

H(η)

| η | NF-apoA | NF-apoB |
| --- | --- | --- |
| 0 | 0 | 0 |
| 5 | 0 | 0 |
| 10 | 0 | 0 |
| 15 | 0 | 0 |
| 20 | 0 | 0 |
| 25 | 0 | 0 |
| 30 | 0 | 0 |
| 35 | 0 | 0 |
| 40 | 0 | 0 |
| 45 | 0 | 0 |
| 50 | 0 | 0 |
| 55 | 0 | 0 |
| 60 | 0 | 0 |
| 65 | 0 | 0 |
| 70 | 0 | 0 |
| 75 | 0 | 0 |
| 80 | 0 | 0 |
| 85 | 0 | 0 |
| 90 | 0 | 0 |
| 95 | 0 | 0 |
| 100 | 0 | 0 |
| 105 | 0 | 0 |
| 110 | 0 | 0 |
| 115 | 0 | 0 |
| 120 | 0 | 0 |
| 125 | 0 | 0 |
| 130 | 0 | 0 |
| 135 | 0 | 0 |
| 140 | 0 | 0 |
| 145 | 9.999e-05 | 0 |
| 150 | 0.00059994 | 0.00089991 |
| 155 | 0.0126987 | 0.00589941 |
| 160 | 0.0753925 | 0.0171983 |
| 165 | 0.208179 | 0.0357964 |
| 170 | 0.309469 | 0.0618938 |
| 175 | 0.257574 | 0.0879912 |
| 180 | 0.10409 | 0.111589 |
| 185 | 0.0272973 | 0.117688 |
| 190 | 0.00409959 | 0.126387 |
| 195 | 0.00049995 | 0.112089 |
| 200 | 0 | 0.0976902 |
| 205 | 0 | 0.079992 |
| 210 | 0 | 0.0641936 |
| 215 | 0 | 0.0414959 |
| 220 | 0 | 0.0221978 |
| 225 | 0 | 0.0128987 |
| 230 | 0 | 0.00339966 |
| 235 | 0 | 0.00049995 |
| 240 | 0 | 0.00019998 |
| 245 | 0 | 0 |
| 250 | 0 | 0 |
| 255 | 0 | 0 |
| 260 | 0 | 0 |
| 265 | 0 | 0 |
| 270 | 0 | 0 |
| 275 | 0 | 0 |
| 280 | 0 | 0 |
| 285 | 0 | 0 |
| 290 | 0 | 0 |
| 295 | 0 | 0 |
| 300 | 0 | 0 |
| 305 | 0 | 0 |
| 310 | 0 | 0 |
| 315 | 0 | 0 |
| 320 | 0 | 0 |
| 325 | 0 | 0 |
| 330 | 0 | 0 |
| 335 | 0 | 0 |
| 340 | 0 | 0 |
| 345 | 0 | 0 |
| 350 | 0 | 0 |
| 355 | 0 | 0 |
| 360 | 0 | 0 |

(c) 303K-apoAB-eta-U48-J2/3

H(η)

| η | NF-apoA | NF-apoB |
| --- | --- | --- |
| 0 | 0.00119988 | 0.00029997 |
| 5 | 0.113589 | 0.00429957 |
| 10 | 0.219278 | 0.00549945 |
| 15 | 0.277772 | 0.00539946 |
| 20 | 0.209079 | 0.00839916 |
| 25 | 0.0930907 | 0.0119988 |
| 30 | 0.0241976 | 0.0152985 |
| 35 | 0.00469953 | 0.0206979 |
| 40 | 0.0019998 | 0.0248975 |
| 45 | 0.00079992 | 0.0314969 |
| 50 | 0.00019998 | 0.0405959 |
| 55 | 0 | 0.0462954 |
| 60 | 0 | 0.0529947 |
| 65 | 0 | 0.0542946 |
| 70 | 0 | 0.0586941 |
| 75 | 0 | 0.0581942 |
| 80 | 0 | 0.0490951 |
| 85 | 0 | 0.0329967 |
| 90 | 0 | 0.0210979 |
| 95 | 0 | 0.0106989 |
| 100 | 0 | 0.00709929 |
| 105 | 0 | 0.00409959 |
| 110 | 0 | 0.0029997 |
| 115 | 0 | 0.00429957 |
| 120 | 0 | 0.00989901 |
| 125 | 0 | 0.019998 |
| 130 | 0 | 0.0409959 |
| 135 | 0 | 0.0767923 |
| 140 | 0 | 0.0858914 |
| 145 | 0 | 0.0605939 |
| 150 | 0 | 0.0224978 |
| 155 | 0 | 0.010099 |
| 160 | 0 | 0.00409959 |
| 165 | 0 | 0.00129987 |
| 170 | 0 | 0 |
| 175 | 0 | 0 |
| 180 | 0 | 0 |
| 185 | 0 | 0 |
| 190 | 0 | 0 |
| 195 | 0 | 0 |
| 200 | 0 | 0 |
| 205 | 0 | 0 |
| 210 | 0 | 0 |
| 215 | 0 | 0 |
| 220 | 0 | 0 |
| 225 | 0 | 0 |
| 230 | 0 | 0 |
| 235 | 0 | 0 |
| 240 | 0 | 0 |
| 245 | 0 | 0 |
| 250 | 0 | 0.00089991 |
| 255 | 0 | 0.00209979 |
| 260 | 0 | 0.00539946 |
| 265 | 0 | 0.00629937 |
| 270 | 0 | 0.00409959 |
| 275 | 0 | 0.00259974 |
| 280 | 0 | 0.0019998 |
| 285 | 0 | 0.00289971 |
| 290 | 0 | 0.00119988 |
| 295 | 0 | 0.00289971 |
| 300 | 0 | 0.00249975 |
| 305 | 0 | 0.00379962 |
| 310 | 0 | 0.00479952 |
| 315 | 0 | 0.00569943 |
| 320 | 0 | 0.00529947 |
| 325 | 0 | 0.00559944 |
| 330 | 0.00019998 | 0.00529947 |
| 335 | 0.00039996 | 0.00549945 |
| 340 | 0.00019998 | 0.00609939 |
| 345 | 0.00159984 | 0.00559944 |
| 350 | 0.00259974 | 0.00479952 |
| 355 | 0.0113989 | 0.00609939 |
| 360 | 0.0376962 | 0.00459954 |

(d) 303K-apoAB-eta-U49-J2/3

H(η)

| η | NF-apoA | NF-apoB |
| --- | --- | --- |
| 0 | 0 | 0 |
| 5 | 0 | 0.00019998 |
| 10 | 0 | 9.999e-05 |
| 15 | 0 | 0 |
| 20 | 0 | 9.999e-05 |
| 25 | 0 | 0.00029997 |
| 30 | 0 | 0 |
| 35 | 0 | 9.999e-05 |
| 40 | 0 | 9.999e-05 |
| 45 | 0 | 0 |
| 50 | 0 | 0.00029997 |
| 55 | 0 | 0.00029997 |
| 60 | 0 | 0.00089991 |
| 65 | 0 | 0.00189981 |
| 70 | 0 | 0.00519948 |
| 75 | 0 | 0.00529947 |
| 80 | 0 | 0.0113989 |
| 85 | 0 | 0.0178982 |
| 90 | 0 | 0.0273973 |
| 95 | 0 | 0.0391961 |
| 100 | 0 | 0.0451955 |
| 105 | 0 | 0.0475952 |
| 110 | 0 | 0.0433957 |
| 115 | 0 | 0.0286971 |
| 120 | 0 | 0.0237976 |
| 125 | 0 | 0.0141986 |
| 130 | 0 | 0.0109989 |
| 135 | 0 | 0.00819918 |
| 140 | 0 | 0.00729927 |
| 145 | 0 | 0.00609939 |
| 150 | 0 | 0.00519948 |
| 155 | 0 | 0.00739926 |
| 160 | 0 | 0.00609939 |
| 165 | 0 | 0.00629937 |
| 170 | 0 | 0.00729927 |
| 175 | 0 | 0.00779922 |
| 180 | 0 | 0.00859914 |
| 185 | 0.00209979 | 0.0119988 |
| 190 | 0.0169983 | 0.0183982 |
| 195 | 0.0955904 | 0.0253975 |
| 200 | 0.261974 | 0.0371963 |
| 205 | 0.352365 | 0.0507949 |
| 210 | 0.205779 | 0.0727927 |
| 215 | 0.0572943 | 0.0859914 |
| 220 | 0.00739926 | 0.0789921 |
| 225 | 0.00049995 | 0.0551945 |
| 230 | 0 | 0.0446955 |
| 235 | 0 | 0.0420958 |
| 240 | 0 | 0.0338966 |
| 245 | 0 | 0.0227977 |
| 250 | 0 | 0.0118988 |
| 255 | 0 | 0.00829917 |
| 260 | 0 | 0.00229977 |
| 265 | 0 | 0.00159984 |
| 270 | 0 | 0.00059994 |
| 275 | 0 | 0 |
| 280 | 0 | 0 |
| 285 | 0 | 0 |
| 290 | 0 | 0 |
| 295 | 0 | 0 |
| 300 | 0 | 0 |
| 305 | 0 | 0 |
| 310 | 0 | 0 |
| 315 | 0 | 0 |
| 320 | 0 | 0 |
| 325 | 0 | 0 |
| 330 | 0 | 0 |
| 335 | 0 | 0.00019998 |
| 340 | 0 | 0 |
| 345 | 0 | 0 |
| 350 | 0 | 0 |
| 355 | 0 | 0 |
| 360 | 0 | 0 |

(e) 303K-apoAB-eta-U36-Loop2

H(η)

| η | NF-apoA | NF-apoB |
| --- | --- | --- |
| 0 | 0.00289971 | 0.00269973 |
| 5 | 0.181082 | 0.115488 |
| 10 | 0.174383 | 0.0961904 |
| 15 | 0.129887 | 0.080192 |
| 20 | 0.0808919 | 0.0939906 |
| 25 | 0.0424958 | 0.10009 |
| 30 | 0.0212979 | 0.114689 |
| 35 | 0.0126987 | 0.0959904 |
| 40 | 0.00919908 | 0.0645935 |
| 45 | 0.00519948 | 0.0333967 |
| 50 | 0.00329967 | 0.00959904 |
| 55 | 0.00059994 | 0.00189981 |
| 60 | 9.999e-05 | 0.00059994 |
| 65 | 0 | 9.999e-05 |
| 70 | 9.999e-05 | 0 |
| 75 | 0 | 0 |
| 80 | 0 | 0 |
| 85 | 0 | 0 |
| 90 | 0 | 0 |
| 95 | 0 | 0 |
| 100 | 0 | 0 |
| 105 | 0 | 0 |
| 110 | 0 | 0 |
| 115 | 0 | 0 |
| 120 | 0 | 0 |
| 125 | 0 | 0 |
| 130 | 0 | 0 |
| 135 | 0 | 0 |
| 140 | 0 | 0 |
| 145 | 0 | 0 |
| 150 | 0 | 0 |
| 155 | 0 | 0 |
| 160 | 0 | 0 |
| 165 | 0 | 0 |
| 170 | 0 | 0 |
| 175 | 0 | 0 |
| 180 | 0 | 0 |
| 185 | 0 | 0 |
| 190 | 0 | 0 |
| 195 | 0 | 0 |
| 200 | 0 | 0 |
| 205 | 0 | 0 |
| 210 | 0 | 0 |
| 215 | 0 | 0 |
| 220 | 0 | 0 |
| 225 | 0 | 0 |
| 230 | 0 | 0 |
| 235 | 0 | 0 |
| 240 | 0 | 0 |
| 245 | 0 | 0 |
| 250 | 0 | 0 |
| 255 | 0 | 0 |
| 260 | 0 | 0 |
| 265 | 0 | 0 |
| 270 | 0 | 0 |
| 275 | 0 | 0 |
| 280 | 0 | 0 |
| 285 | 0 | 0 |
| 290 | 0 | 0 |
| 295 | 0 | 0 |
| 300 | 0 | 0 |
| 305 | 0 | 0 |
| 310 | 0 | 0 |
| 315 | 0 | 0 |
| 320 | 0 | 0 |
| 325 | 0 | 9.999e-05 |
| 330 | 0.00089991 | 0.00029997 |
| 335 | 0.0019998 | 0.00089991 |
| 340 | 0.00739926 | 0.00239976 |
| 345 | 0.0227977 | 0.00819918 |
| 350 | 0.0520948 | 0.0253975 |
| 355 | 0.0974903 | 0.0561944 |
| 360 | 0.153185 | 0.0969903 |

**Figure S10: Some representative Histogram distribution of pseudo-torsion angle (θ) at 303K (apoA in blue and apoB in orange))**

(a) 303K-apoAB-theta-A17-Stem1

H(θ)

| θ | NF-apoA | NF-apoB |
| --- | --- | --- |
| 0 | 0 | 0 |
| 5 | 0 | 0 |
| 10 | 0 | 0 |
| 15 | 0 | 0 |
| 20 | 0 | 0 |
| 25 | 0 | 0 |
| 30 | 0 | 0 |
| 35 | 0 | 0 |
| 40 | 0 | 0 |
| 45 | 0 | 0 |
| 50 | 0 | 0 |
| 55 | 0 | 0 |
| 60 | 0 | 0 |
| 65 | 0 | 0 |
| 70 | 0 | 0 |
| 75 | 0 | 0 |
| 80 | 0 | 0 |
| 85 | 0 | 0 |
| 90 | 0 | 0 |
| 95 | 0 | 0 |
| 100 | 0 | 0 |
| 105 | 0 | 0 |
| 110 | 0 | 0 |
| 115 | 0 | 0 |
| 120 | 0 | 0 |
| 125 | 0 | 0 |
| 130 | 0 | 0 |
| 135 | 0 | 0 |
| 140 | 0 | 0 |
| 145 | 0 | 0 |
| 150 | 0 | 0 |
| 155 | 0 | 0 |
| 160 | 0 | 0 |
| 165 | 0 | 0 |
| 170 | 0.00019998 | 9.999e-05 |
| 175 | 0.00079992 | 0.00019998 |
| 180 | 0.00079992 | 0.00059994 |
| 185 | 0.00359964 | 0.0019998 |
| 190 | 0.00609939 | 0.00639936 |
| 195 | 0.00929907 | 0.0169983 |
| 200 | 0.0141986 | 0.049695 |
| 205 | 0.0175982 | 0.105089 |
| 210 | 0.0292971 | 0.169983 |
| 215 | 0.0543946 | 0.212079 |
| 220 | 0.0959904 | 0.20328 |
| 225 | 0.135286 | 0.134687 |
| 230 | 0.139086 | 0.0627937 |
| 235 | 0.108589 | 0.0246975 |
| 240 | 0.111189 | 0.00659934 |
| 245 | 0.125187 | 0.00309969 |
| 250 | 0.0888911 | 0.00119988 |
| 255 | 0.0418958 | 0.00039996 |
| 260 | 0.0137986 | 0 |
| 265 | 0.00329967 | 9.999e-05 |
| 270 | 0.00049995 | 0 |
| 275 | 0 | 0 |
| 280 | 0 | 0 |
| 285 | 0 | 0 |
| 290 | 0 | 0 |
| 295 | 0 | 0 |
| 300 | 0 | 0 |
| 305 | 0 | 0 |
| 310 | 0 | 0 |
| 315 | 0 | 0 |
| 320 | 0 | 0 |
| 325 | 0 | 0 |
| 330 | 0 | 0 |
| 335 | 0 | 0 |
| 340 | 0 | 0 |
| 345 | 0 | 0 |
| 350 | 0 | 0 |
| 355 | 0 | 0 |
| 360 | 0 | 0 |

**Figure S11: Some representative Histogram distribution of pseudo-torsion angle (η) at 313K (apoA in blue and apoB in orange))**

**Figure S12: Some representative Histogram distribution of pseudo-torsion angle (θ) at 313K (apoA in blue and apoB in orange))**

**Figure S13: Some representative Histogram distribution of pseudo-torsion angle (η) at 323K (apoA in blue and apoB in orange))**

**Figure S14: Some representative Histogram distribution of pseudo-torsion angle (θ) at 323K (apoA in blue and apoB in orange))**

**Figure S15: Some representative Histogram distribution of pseudo-torsion angle (η) at 373K (apoA in blue and apoB in orange))**

**Figure S16: Some representative Histogram distribution of pseudo-torsion angle (θ) at 373K (apoA in blue and apoB in orange))**

Supplementary table

Numbering according to PDB file

Table S1 (a): The changes in conformational thermodynamics (kJ/mol) in the residues of Stem1 of apoA adenine riboswitch with respect to apoB adenine riboswitch at 283K

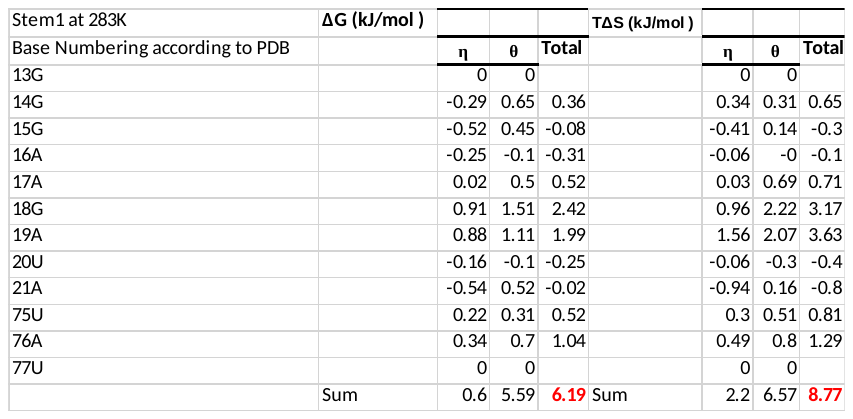

Table S1 (b): The changes in conformational thermodynamics (kJ/mol) in the residues of Stem1 of apoA adenine riboswitch with respect to apoB adenine riboswitch at 293K

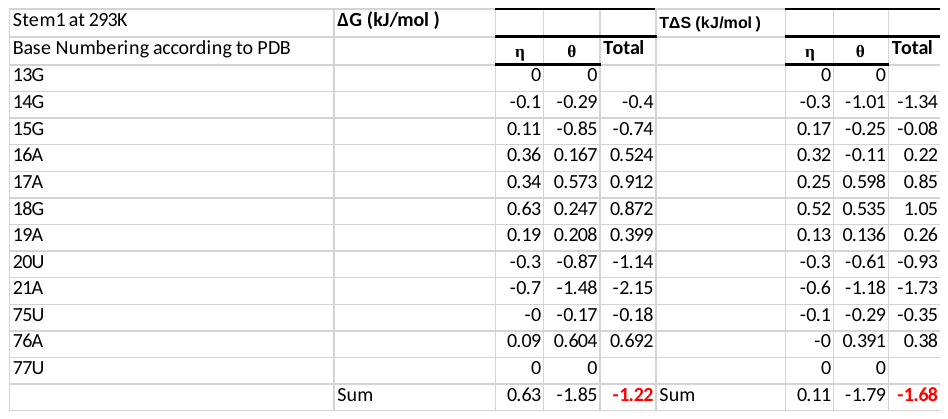

Table S1 (c): The changes in conformational thermodynamics (kJ/mol) in the residues of Stem1 of apoA adenine riboswitch with respect to apoB adenine riboswitch at 303K

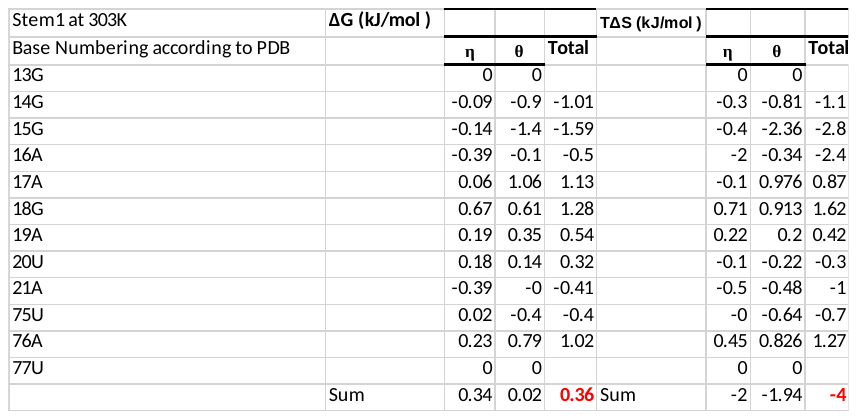

Table S1 (d): The changes in conformational thermodynamics (kJ/mol) in the residues of Stem1 of apoA adenine riboswitch with respect to apoB adenine riboswitch at 313K

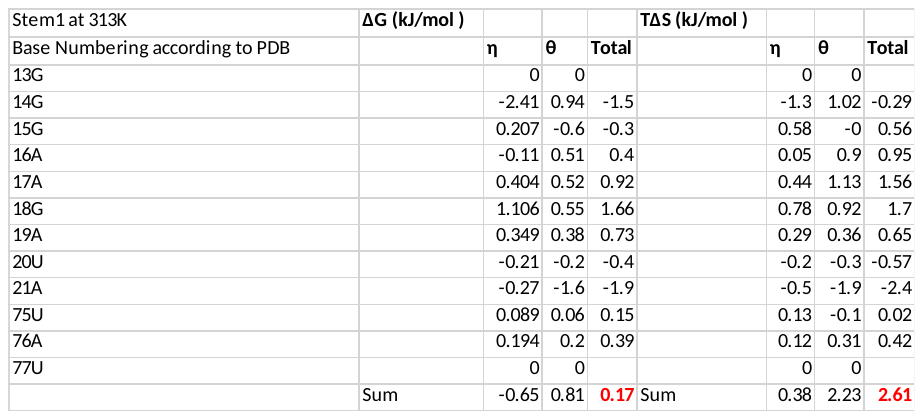

Table S1 (e): The changes in conformational thermodynamics (kJ/mol) in the residues of Stem1 of apoA adenine riboswitch with respect to apoB adenine riboswitch at 323K

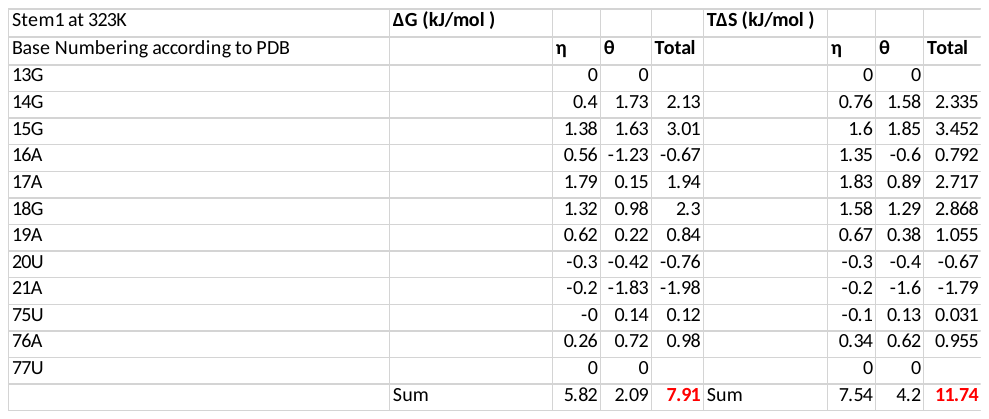

Table S1 (f): The changes in conformational thermodynamics (kJ/mol) in the residues of Stem1 of apoA adenine riboswitch with respect to apoB adenine riboswitch at 373K

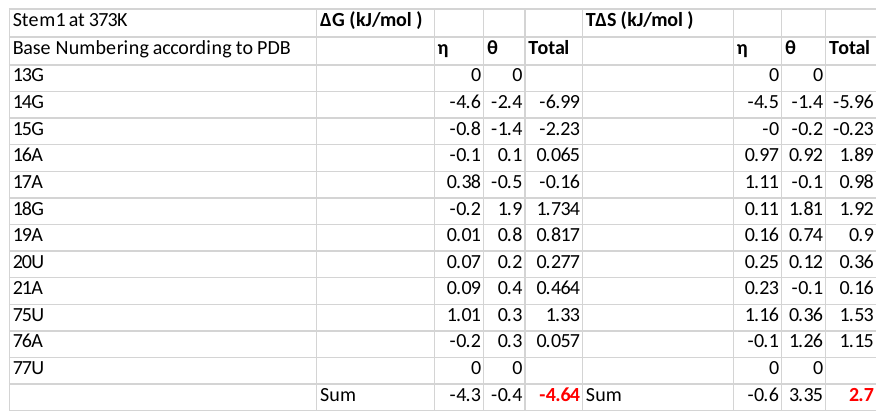

Table S2 (a): The changes in conformational thermodynamics (kJ/mol) in the residues of Stem2 of apoA adenine riboswitch with respect to apoB adenine riboswitch at 283K

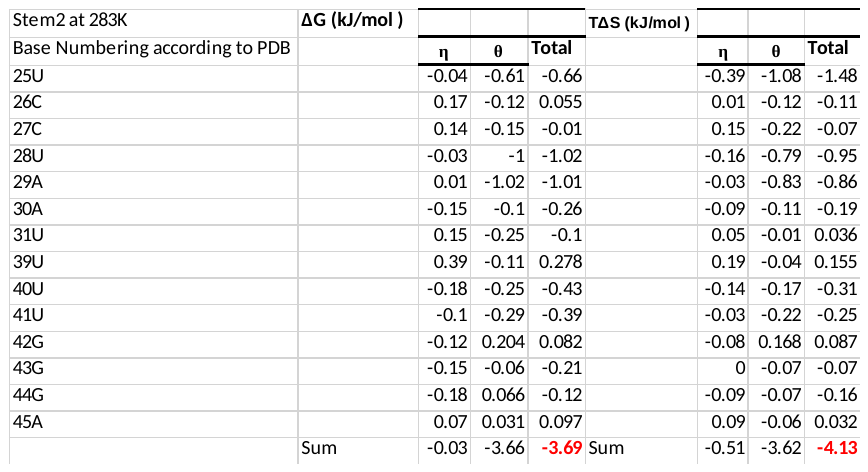

Table S2 (b): The changes in conformational thermodynamics (kJ/mol) in the residues of Stem2 of apoA adenine riboswitch with respect to apoB adenine riboswitch at 293K

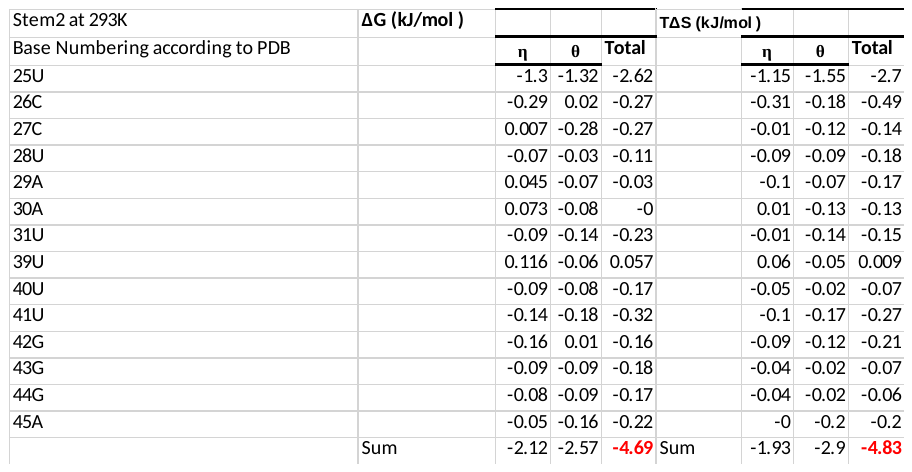

Table S2 (c): The changes in conformational thermodynamics (kJ/mol) in the residues of Stem2 of apoA adenine riboswitch with respect to apoB adenine riboswitch at 303K

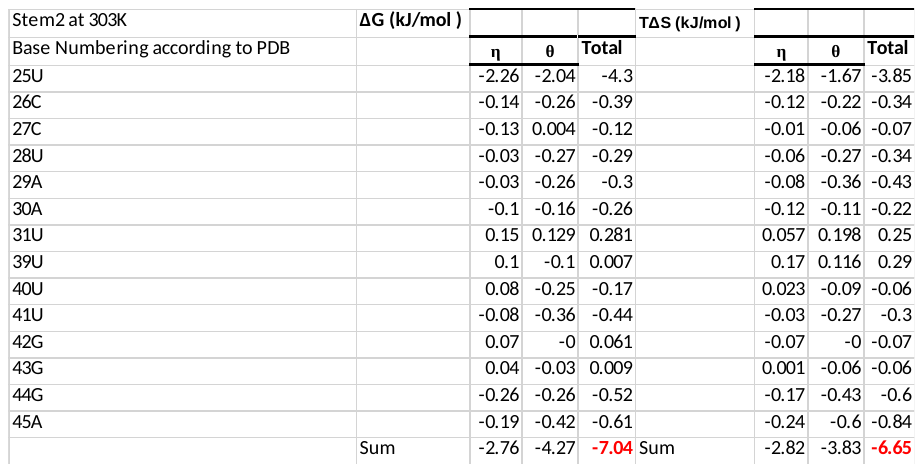

Table S2 (d): The changes in conformational thermodynamics (kJ/mol) in the residues of Stem2 of apoA adenine riboswitch with respect to apoB adenine riboswitch at 313K

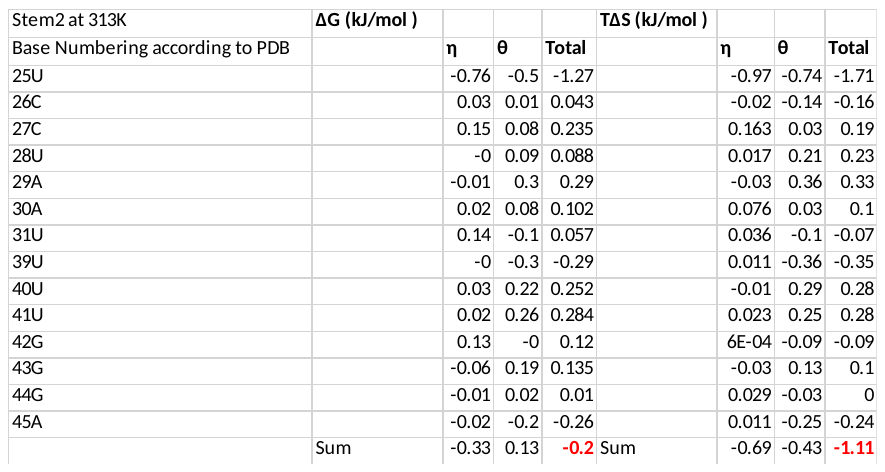

Table S2 (e): The changes in conformational thermodynamics (kJ/mol) in the residues of Stem2 of apoA adenine riboswitch with respect to apoB adenine riboswitch at 323K

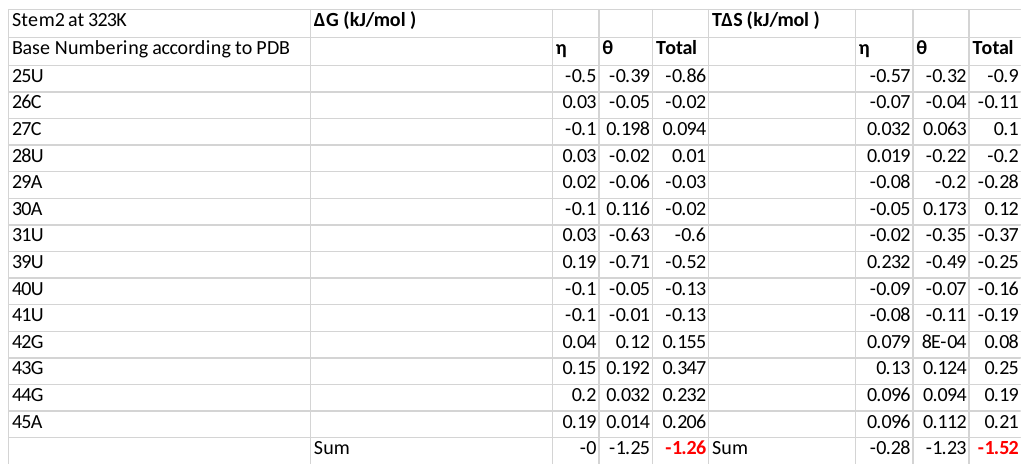

Table S2 (f): The changes in conformational thermodynamics (kJ/mol) in the residues of Stem2 of apoA adenine riboswitch with respect to apoB adenine riboswitch at 373K

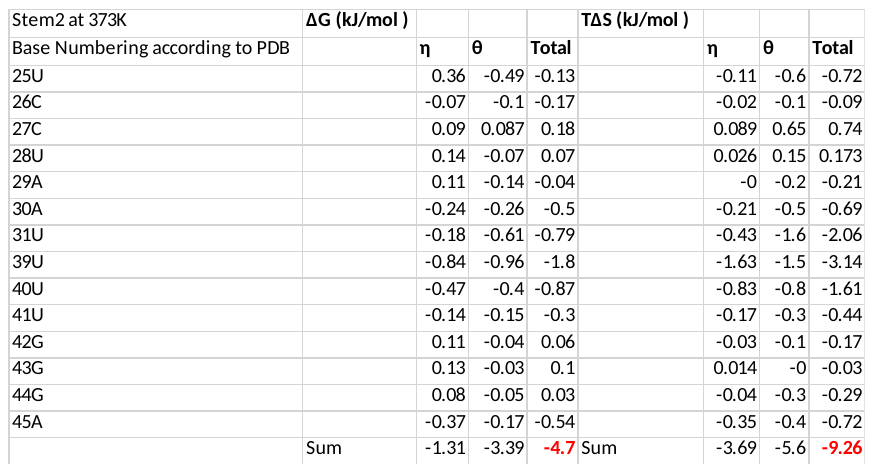

Table S3 (a): The changes in conformational thermodynamics (kJ/mol) in the residues of Stem3 of apoA adenine riboswitch with respect to apoB adenine riboswitch at 283K

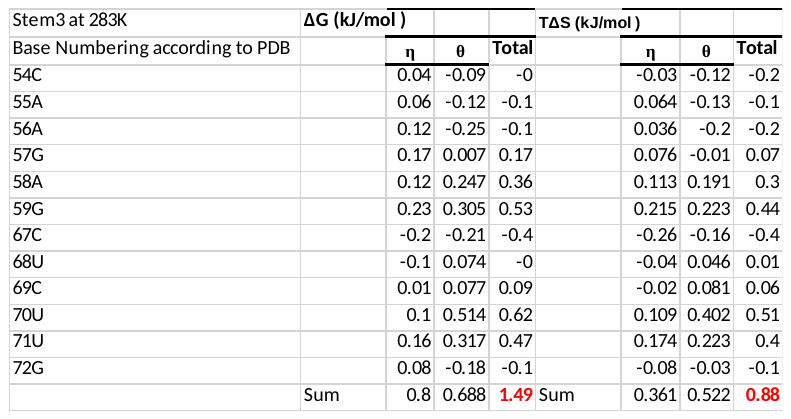

Table S3 (b): The changes in conformational thermodynamics (kJ/mol) in the residues of Stem3 of apoA adenine riboswitch with respect to apoB adenine riboswitch at 293K

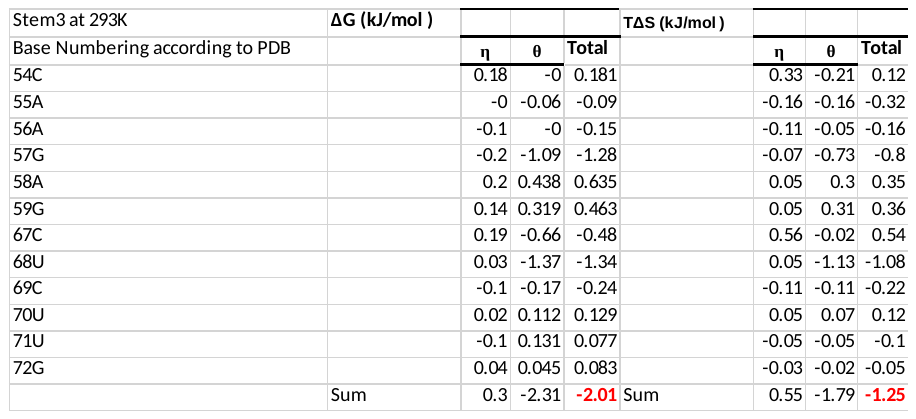

Table S3 (c): The changes in conformational thermodynamics (kJ/mol) in the residues of Stem3 of apoA adenine riboswitch with respect to apoB adenine riboswitch at 303K

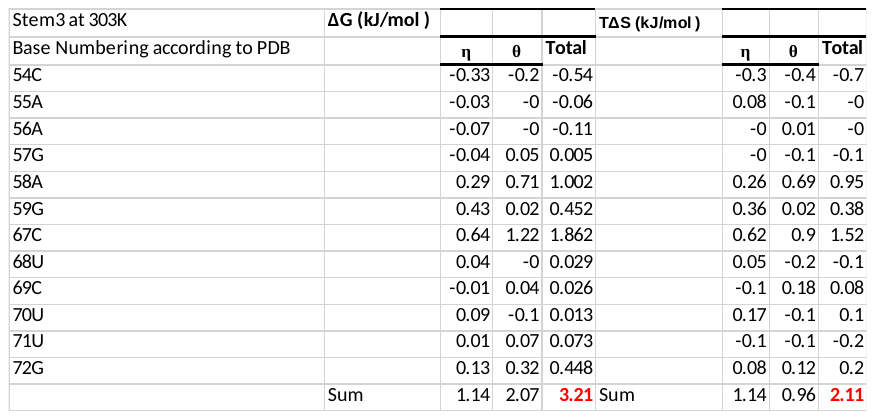

Table S3 (d): The changes in conformational thermodynamics (kJ/mol) in the residues of Stem3 of apoA adenine riboswitch with respect to apoB adenine riboswitch at 313K

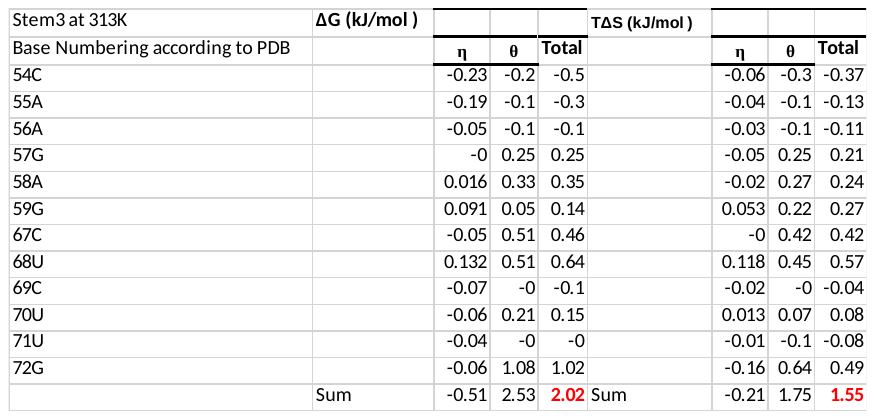

Table S3 (e): The changes in conformational thermodynamics (kJ/mol) in the residues of Stem3 of apoA adenine riboswitch with respect to apoB adenine riboswitch at 323K

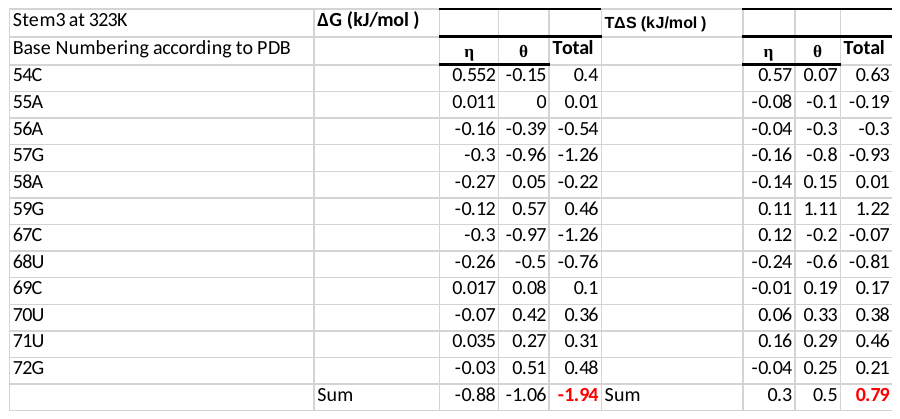

Table S3 (f): The changes in conformational thermodynamics (kJ/mol) in the residues of Stem3 of apoA adenine riboswitch with respect to apoB adenine riboswitch at 373K

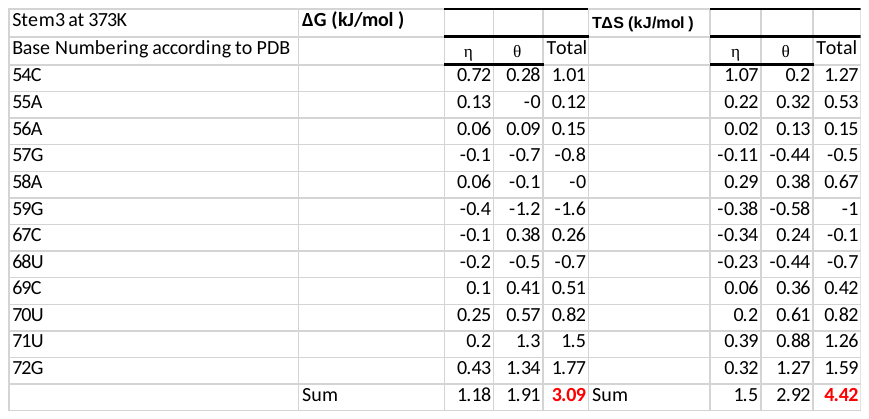

Table S4 (a): The changes in conformational thermodynamics (kJ/mol) in the residues of Loop2 of apoA adenine riboswitch with respect to apoB adenine riboswitch at 283K

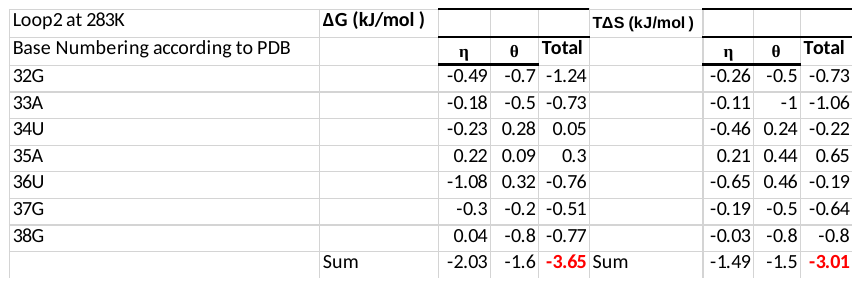

Table S4 (b): The changes in conformational thermodynamics (kJ/mol) in the residues of Loop2 of apoA adenine riboswitch with respect to apoB adenine riboswitch at 293K

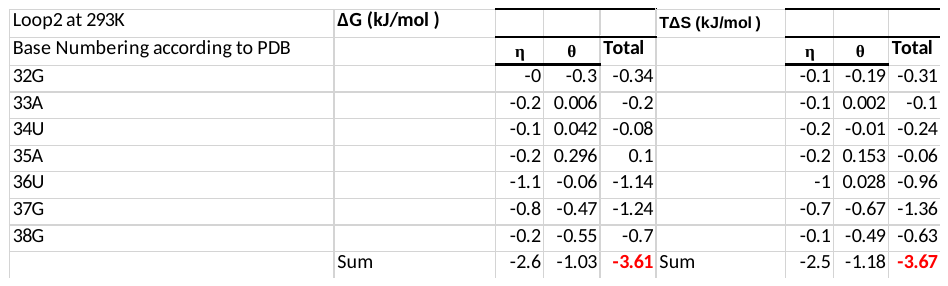

Table S4 (c): The changes in conformational thermodynamics (kJ/mol) in the residues of Loop2 of apoA adenine riboswitch with respect to apoB adenine riboswitch at 303K

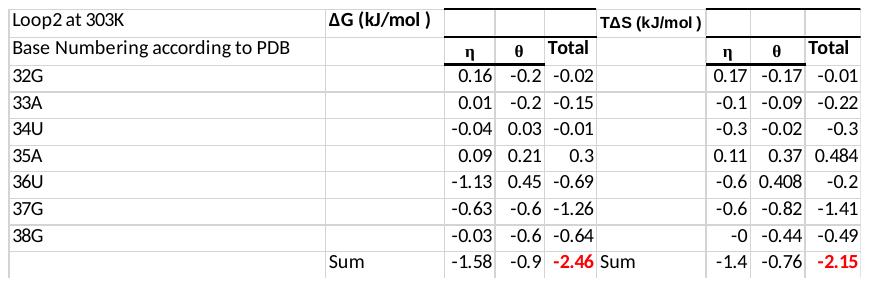

Table S4 (d): The changes in conformational thermodynamics (kJ/mol) in the residues of Loop2 of apoA adenine riboswitch with respect to apoB adenine riboswitch at 313K

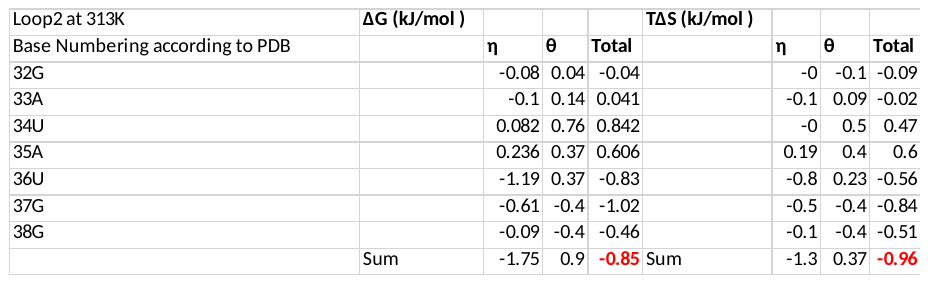

Table S4 (e): The changes in conformational thermodynamics (kJ/mol) in the residues of Loop2 of apoA adenine riboswitch with respect to apoB adenine riboswitch at 323K

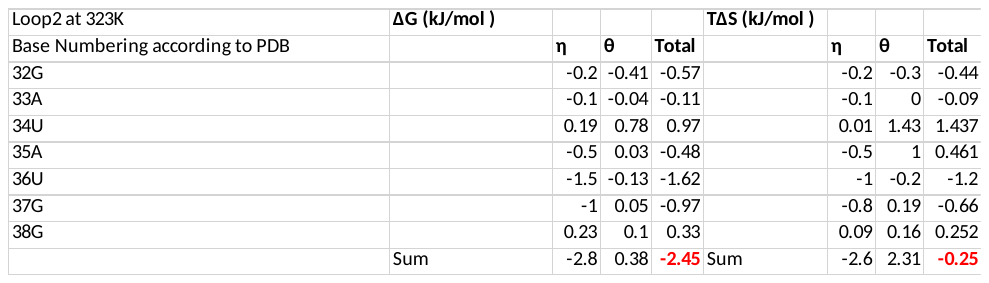

Table S4 (f): The changes in conformational thermodynamics (kJ/mol) in the residues of Loop2 of apoA adenine riboswitch with respect to apoB adenine riboswitch at 373K

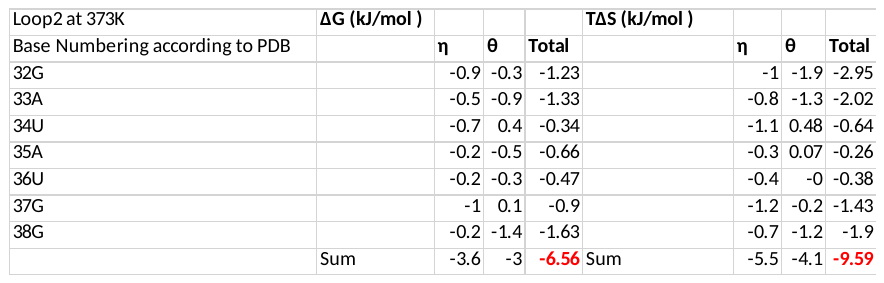

Table S5 (a): The changes in conformational thermodynamics (kJ/mol) in the residues of Loop3 of apoA adenine riboswitch with respect to apoB adenine riboswitch at 283K

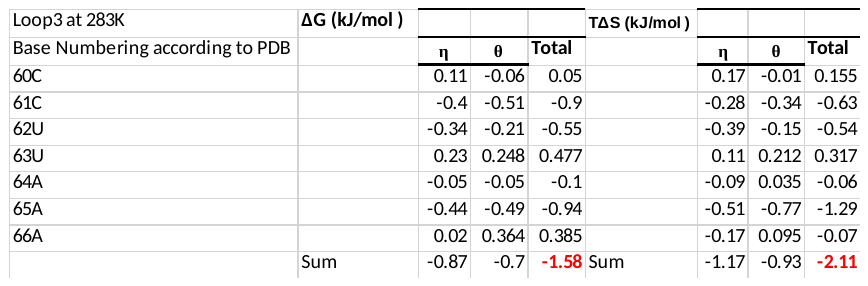

Table S5 (b): The changes in conformational thermodynamics (kJ/mol) in the residues of Loop3 of apoA adenine riboswitch with respect to apoB adenine riboswitch at 293K

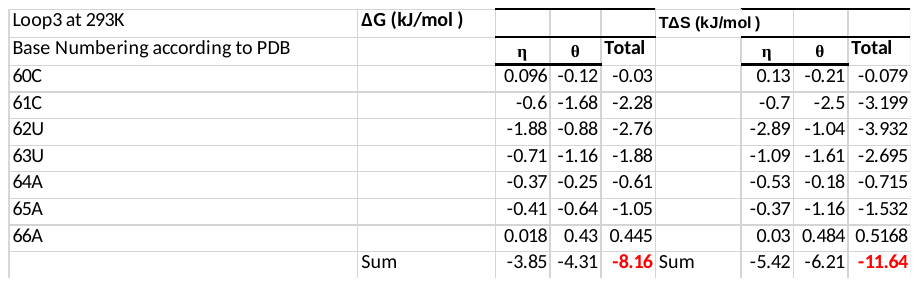

Table S5 (c): The changes in conformational thermodynamics (kJ/mol) in the residues of Loop3 of apoA adenine riboswitch with respect to apoB adenine riboswitch at 303K

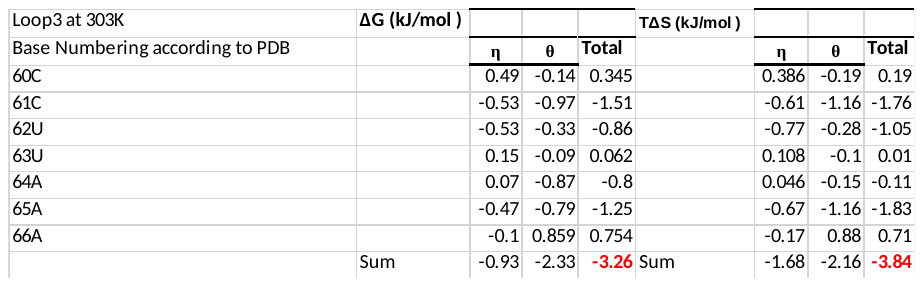

Table S5 (d): The changes in conformational thermodynamics (kJ/mol) in the residues of Loop3 of apoA adenine riboswitch with respect to apoB adenine riboswitch at 313K

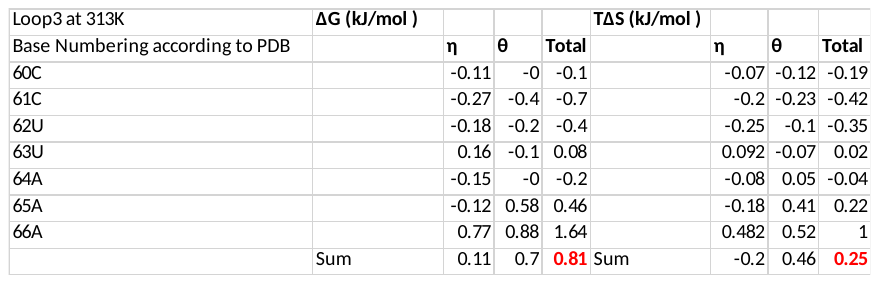

Table S5 (e): The changes in conformational thermodynamics (kJ/mol) in the residues of Loop3 of apoA adenine riboswitch with respect to apoB adenine riboswitch at 323K

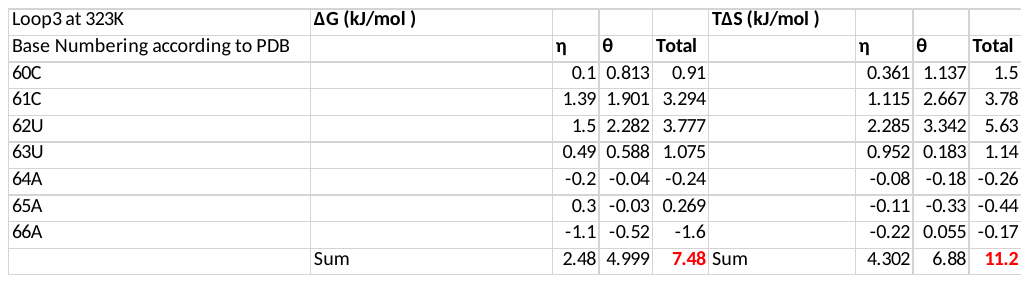

Table S5 (f): The changes in conformational thermodynamics (kJ/mol) in the residues of Loop3 of apoA adenine riboswitch with respect to apoB adenine riboswitch at 373K

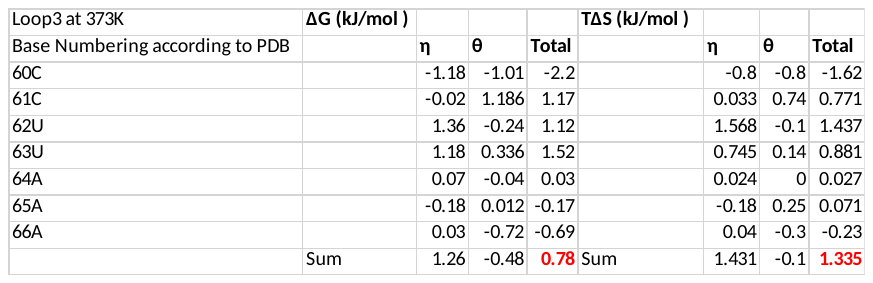

Table S6 (a): The changes in conformational thermodynamics (kJ/mol) in the residues of J_1/2_ Hinge of apoA adenine riboswitch with respect to apoB adenine riboswitch at 283K

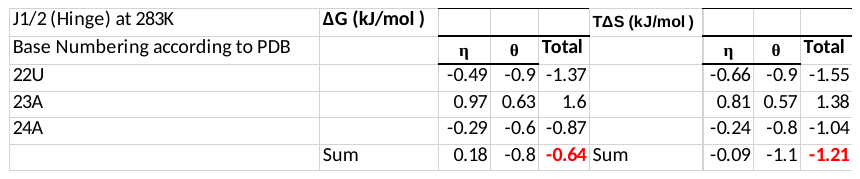

Table S6 (b): The changes in conformational thermodynamics (kJ/mol) in the residues of J_1/2_ Hinge of apoA adenine riboswitch with respect to apoB adenine riboswitch at 293K

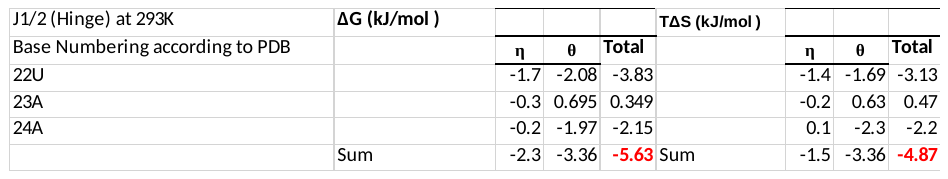

Table S6 (c): The changes in conformational thermodynamics (kJ/mol) in the residues of J_1/2_ Hinge of apoA adenine riboswitch with respect to apoB adenine riboswitch at 303K

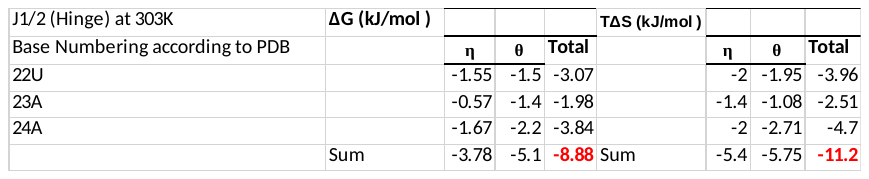

Table S6 (d): The changes in conformational thermodynamics (kJ/mol) in the residues of J_1/2_ Hinge of apoA adenine riboswitch with respect to apoB adenine riboswitch at 313K

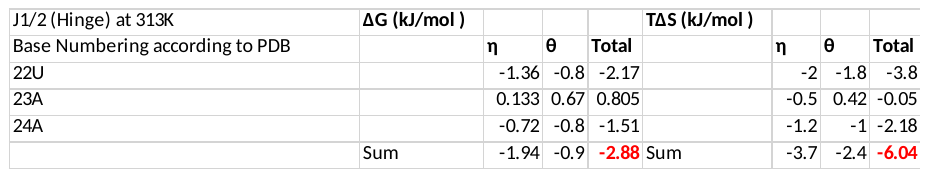

Table S6 (e): The changes in conformational thermodynamics (kJ/mol) in the residues of J_1/2_ Hinge of apoA adenine riboswitch with respect to apoB adenine riboswitch at 323K

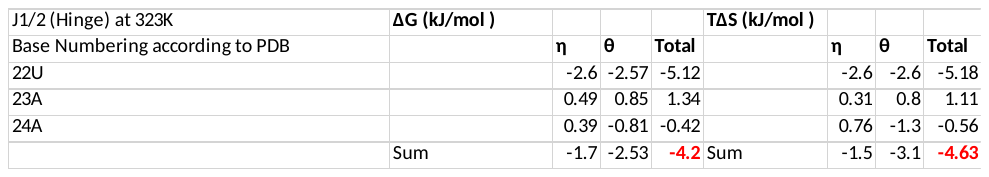

Table S6 (f): The changes in conformational thermodynamics (kJ/mol) in the residues of J_1/2_ Hinge of apoA adenine riboswitch with respect to apoB adenine riboswitch at 373K

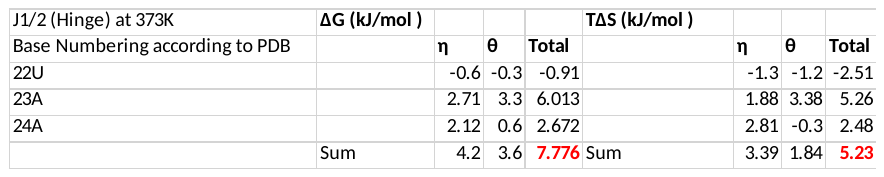

Table S7 (a): The changes in conformational thermodynamics (kJ/mol) in the residues of J_2/3_ Latch of apoA adenine riboswitch with respect to apoB adenine riboswitch at 283K

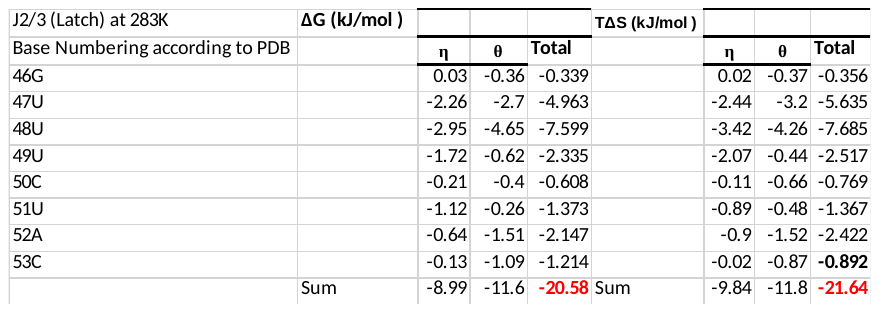

Table S7 (b): The changes in conformational thermodynamics (kJ/mol) in the residues of J_2/3_ Latch of apoA adenine riboswitch with respect to apoB adenine riboswitch at 293K

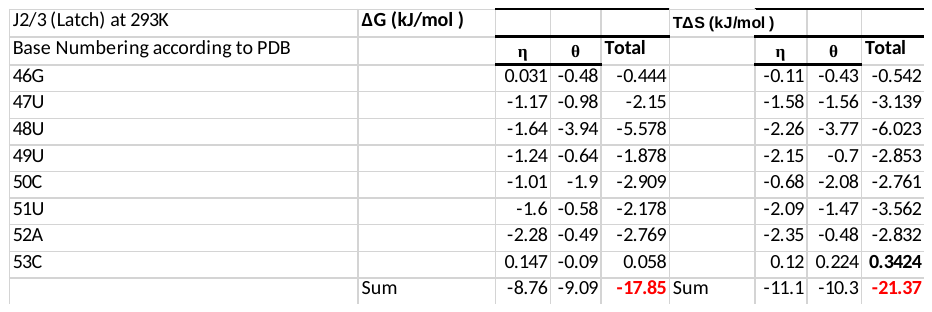

Table S7 (c): The changes in conformational thermodynamics (kJ/mol) in the residues of J_2/3_ Latch of apoA adenine riboswitch with respect to apoB adenine riboswitch at 303K

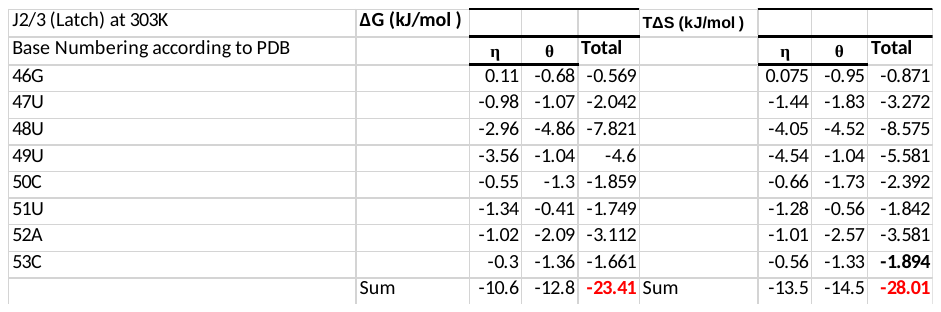

Table S7 (d): The changes in conformational thermodynamics (kJ/mol) in the residues of J_2/3_ Latch of apoA adenine riboswitch with respect to apoB adenine riboswitch at 313K

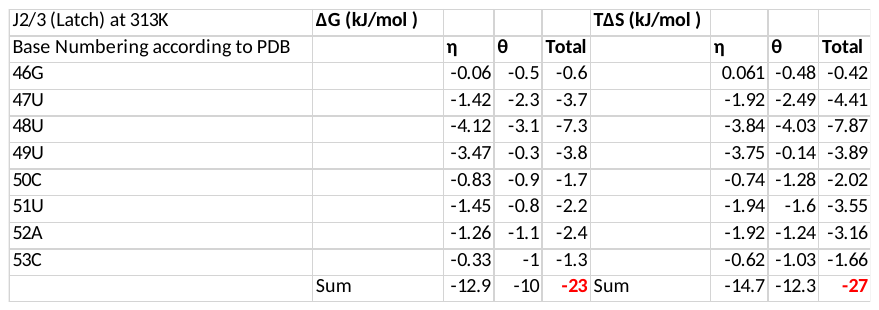

Table S7 (e): The changes in conformational thermodynamics (kJ/mol) in the residues of J_2/3_ Latch of apoA adenine riboswitch with respect to apoB adenine riboswitch at 323K

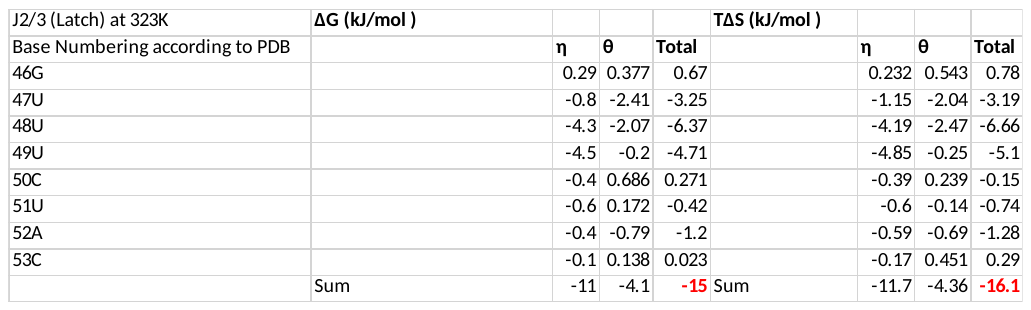

Table S7 (f): The changes in conformational thermodynamics (kJ/mol) in the residues of J_2/3_ Latch Hinge of apoA adenine riboswitch with respect to apoB adenine riboswitch at 373K

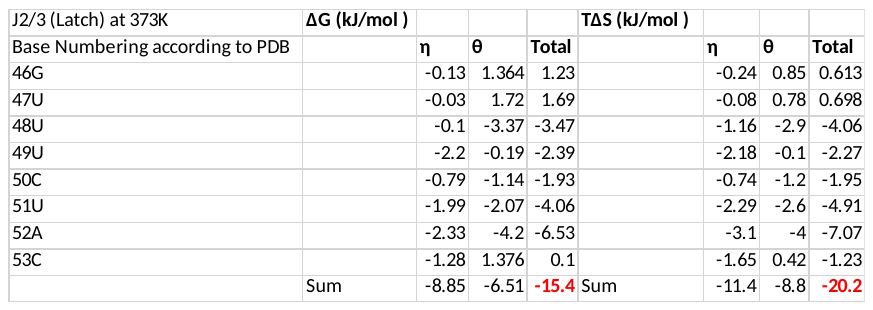
